# Intranasal SARS-CoV-2 RBD decorated nanoparticle vaccine enhances viral clearance in the Syrian hamster model

**DOI:** 10.1101/2022.10.27.514054

**Authors:** DR Patel, AM Minns, DG Sim, CJ Field, AE Kerr, T Heinly, EH Luley, R.M. Rossi, C. Bator, I.M. Moustafa, SL Hafenstein, SE Lindner, TC Sutton

## Abstract

Multiple vaccines have been developed and licensed for SARS-CoV-2. While these vaccines reduce disease severity, they do not prevent infection, and SARS-CoV-2 continues to spread and evolve. To prevent infection and limit transmission, vaccines must be developed that induce immunity in the respiratory tract. Therefore, we performed proof-of-principle vaccination studies with an intranasal nanoparticle vaccine against SARS-CoV-2. The vaccine candidate consisted of the self-assembling 60-subunit I3-01 protein scaffold covalently decorated with the SARS-CoV-2 receptor binding domain (RBD) using the SpyCatcher-SpyTag system. We verified the intended antigen display features by reconstructing the I3-01 scaffold to 3.4A using cryo-EM, and then demonstrated that the scaffold was highly saturated when grafted with RBD. Using this RBD-grafted SpyCage scaffold (RBD+SpyCage), we performed two unadjuvanted intranasal vaccination studies in the “gold-standard” preclinical Syrian hamster model. Hamsters received two vaccinations 28 days apart, and were then challenged 28 days post-boost with SARS-CoV-2. The initial study focused on assessing the immunogenicity of RBD+SpyCage, which indicated that vaccination of hamsters induced a non-neutralizing antibody response that enhanced viral clearance but did not prevent infection. In an expanded study, we demonstrated that covalent bonding of RBD to the scaffold was required to induce an antibody response. Consistent with the initial study, animals vaccinated with RBD+SpyCage more rapidly cleared SARS-CoV-2 from both the upper and lower respiratory tract. These findings demonstrate the intranasal SpyCage vaccine platform can induce protection against SARS-CoV-2 and, with additional modifications to improve immunogenicity, is a versatile platform for the development of intranasal vaccines targeting respiratory pathogens.

**IMPORTANCE:** Despite the availability of efficacious COVID vaccines that reduce disease severity, SARS-CoV-2 continues to spread. To limit SARS-CoV-2 transmission, the next generation of vaccines must induce immunity in the mucosa of the upper respiratory tract. Therefore, we performed proof-of-principle, unadjuvanted intranasal vaccination studies with a recombinant protein nanoparticle scaffold, SpyCage, decorated with the receptor-binding domain (RBD) of the S protein (SpyCage+RBD). We show that SpyCage+RBD was immunogenic and enhanced SARS-CoV-2 clearance from the nose and lungs of Syrian hamsters. Moreover, covalent grafting of the RBD to the scaffold was required to induce an immune response when given via the intranasal route. These proof-of-concept findings indicate that with further enhancements to immunogenicity (e.g., adjuvant incorporation, antigen optimization), the SpyCage scaffold has potential as a versatile, intranasal vaccine platform for respiratory pathogens.

## INTRODUCTION

Severe acute respiratory syndrome coronavirus-2 (SARS-CoV-2) is the etiological agent of the coronavirus disease 2019 (COVID-19)^1^ pandemic. SARS-CoV-2 is an enveloped betacoronavirus with a non-segmented positive-sense single-stranded RNA genome. The genome encodes 4 structural proteins: spike (S), membrane (M), envelope (E), and nucleocapsid (N), as well as multiple non-structural proteins^2^. The S protein is the major surface protein and mediates viral entry and fusion. The receptor-binding domain (RBD) of the S protein binds the host receptor angiotensin-converting enzyme 2 (ACE2), leading to endocytosis of the virion and infection of the host^2,3^. Importantly, antibody responses against SARS-CoV-2 in humans and experimentally infected animals are predominantly directed toward the S protein. Moreover, titers of RBD-binding antibodies correlate with neutralizing activity, and RBD is considered the immunodominant region of the S protein ^4,5^. Therefore, RBD represents a suitable immunogen for vaccine development, and blocking this domain has the potential to prevent infection.

Prior to the SARS-CoV-2 pandemic, vaccines targeting coronaviruses in humans had not been advanced through late-stage clinical trials. Development of multiple SARS-CoV-2 vaccine candidates was enabled by rapid sequencing of the viral genome as well as pre-existing knowledge about vaccination against severe acute respiratory syndrome coronavirus (SARS-CoV) and Middle Eastern respiratory syndrome coronavirus (MERS-CoV)^6^. Currently, there are at least 12 vaccines approved for human use ^7,8^. Licensed vaccines such as CoronaVac and QazCovid-in contain inactivated virus ^9–13^, while vaccines developed by Pfizer/BioNTech and Moderna consist of mRNA encoding the pre-fusion S protein enclosed in a lipid nanoparticle^14^. The Novavax vaccine contains recombinant S protein, and vaccines from Johnson & Johnson and AstraZeneca use viral vectors to deliver DNA encoding the S protein^15^. Importantly, all of these licensed vaccines are delivered by intramuscular (*i.m.*) injection, and these vaccines have been shown to reduce the severity of SARS-CoV-2 infection^11,16,17^; however, these vaccines do not prevent infection, and vaccinated individuals can develop symptomatic infections and transmit the virus onwards.

Intramuscular vaccination induces a systemic immune response with high titers of IgG antibodies that enter the lungs to reduce viral replication and disease severity^6,18^. However, the delivery of vaccines via the *i.m*. route does not induce a strong mucosal immune response^6^, and a mucosal response is required to prevent infection of the upper respiratory tract and limit transmission. In contrast to *i.m*. administered vaccines, an efficacious intranasal vaccine has the potential to protect mucosal surfaces via the induction of secretory IgA antibodies and mucosal T cells. Moreover, these vaccines can also induce a serum IgG response that can impart similar disease reductions as observed for existing vaccines (reviewed in ^19^). However, on-going analyses of licensed SARS-CoV-2 vaccine efficacy has shown that vaccine-induced immunity wanes over time, resulting in breakthrough infections ^20–22^. In addition, intranasal administration of mRNA vaccines to mice did not confer protection against SARS-CoV-2 challenge^23^. As a result, there is a growing need to develop a second generation of SARS-CoV-2 vaccines that can be administered through intranasal routes to induce protective mucosal immunity ^24,25^.

To date, a limited number of intranasal vaccine candidates have been developed against SARS-CoV-2. Most of these candidates are viral vectors or live-attenuated vaccines; however, there have been safety concerns with viral vectored SARS-CoV-2 vaccines, and their administration is limited to individuals older than 18 years of age^26,27^. Moreover, the only licensed live-attenuated intranasal vaccine is against influenza, and due to safety concerns and poor immunogenicity in older individuals, its use is restricted to individuals 2-49 years of age^28^. Therefore, there is a need to develop intranasal vaccines that would be suitable for individuals of all ages.

To address this gap, we adapted the I3-01 self-assembling protein into a nanoparticle bearing a flexible SpyCatcher domain (SpyCage) to display SARS-CoV-2 RBD/SpyTag (RBD+SpyCage) as an intranasal vaccine. The I3-01-based platform has been shown to be an excellent immunization scaffold to present a variety of antigens from viral (SARS-CoV-2, influenza, EBV, CSFV) and parasitic (*Plasmodium*) pathogens that reproducibly boosts immune responses as compared to the unscaffolded antigen^29–36^. However, these trials have been restricted to *i.m*. injections with immune responses as endpoint readouts, with a few notable studies proceeding through challenges with live pathogens^35,37^.

Here we performed proof-of-principle studies to evaluate RBD grafted to the SpyCage scaffold (RBD+SpyCage) as an intranasal vaccine in the “gold-standard” Syrian hamster model. Syrian hamsters are highly permissive to SARS-CoV-2 infection, and the virus efficiently transmits in these animals by direct contact and respiratory droplets (*i.e*., airborne transmission)^38–40^. We performed two separate efficacy studies in which hamsters were given a prime and boost intranasal vaccination and challenged with SARS-CoV-2. We demonstrated covalent grafting of RBD to SpyCage was required to induce an IgG antibody response in vaccinated animals. Upon SARS-CoV-2 challenge, regardless of vaccination status, all hamsters became infected and exhibited weight loss; however, animals vaccinated with RBD+SpyCage more rapidly cleared the virus from both the upper and lower respiratory tract with evidence of reduced lung pathology. Collectively, these studies demonstrate the potential for SpyCage as the basis of an intranasal vaccine platform for SARS-CoV-2 and possibly other respiratory pathogens.

## MATERIALS AND METHODS

### Production and Purification of Apo Cage and SpyCage

The Apo Cage scaffold is based upon the 6xHis/I3-01 protein described previously by Hsia and colleagues^41,42^. The SpyCage scaffold consists of a genetic fusion of a 6xHis tag, the SpyCatcher domain, a flexible linker, and the I3-01 protein^41,43^. These proteins were expressed in the *E. coli* BL21 (DE3) CodonPlus strain bearing either plasmid pSL1013 (Apo Cage) or pSL1040 (SpyCage) using a modified pET28 vector. Cultures were grown in LB media at 37°C to an OD600 of ~0.5, at which point protein expression was induced by the addition of 0.5mM IPTG (final concentration) for 2.5 hours. Cell pellets were suspended in 50mL of resuspension buffer (50 mM Tris-Cl pH8.0 at room temperature(RT), 500 mM NaCl) per 1L of culture, and cells were lysed by sonication using a disruptor horn attachment, using three pulses of 30 seconds each at 70% amplitude and 50% duty cycle (model 450 Branson Digital Sonifier). The crude extract was spun at 15,500 *xg* for 20 minutes at 4C, and the soluble fraction was then incubated in batch with 2 ml of equilibrated Ni-NTA resin for 1 hour at 4C. The resin was applied to a gravity flow column and washed with 50 mL of resuspension buffer followed by 50 mL of Mid-Imidazole buffer (25 mM Tris-Cl pH7.5@RT, 500 mM NaCl, 50 mM Imidazole, 250 mM dextrose, 10% v/v glycerol). Apo cage and SpyCage protein were eluted using elution buffer (50 mM Tris-Cl pH8.0@RT, 500 mM NaCl, 300 mM imidazole), and then were exhaustively dialyzed into 50 mM Tris-Cl pH8.0@RT, 500 mM NaCl, 1 mM DTT, 10% v/v glycerol. The dialyzed material was then concentrated to ~2.0 mg/ml using Amicon Ultra Centrifugal Filters (Fisher Scientific Cat#: UFC9-003-08) and snap-frozen in liquid nitrogen for long-term storage at −80C. Complete plasmid sequences are provided in Supp File 1.

### Production and Purification of SARS-CoV-2 Spike RBD

The Receptor-Binding Domain (RBD) of SARS-CoV2-2 Spike protein was produced with and without a C-terminal SpyTag for covalent attachment to SpyCage using plasmid pSL1515 and pSL1510 respectively^43^. Plasmid DNA was purified (Qiagen HiSpeed Maxiprep Kit) precipitated with ethanol, and resuspended in water before transfection using the Expi293 Expression System (ThermoFisher, Expi293F cells, Expi293 Media, and the ExpiFectamine 293 Transfection Kit) by the Penn State Sartorius Cell Culture Facility as per manufacturer instructions. Briefly, cells maintained in log phase growth at 37°C and 8% CO2 in baffled flasks shaking at 120-130 rpm were transfected at a concentration of 5 × 10^6^/ml, and were supplemented by the addition of ExpiFectamine 293 Transfection Enhancer 1 & 2 approximately 20 hours post-transfection. Culture supernatant was harvested by centrifugation (274 *xg*, 5 minutes, RT) on day three, and was incubated in batch with Ni-NTA (ThermoSci HisPur) resin pre-equilibrated in 1xPBS at 4°C for 1 hour on a nutator. The resin was then applied to a gravity flow column and was washed four times with 10 column volumes of wash buffer (57 mM NaH2PO4 pH 6.3@RT, 30 mM NaCl, 20 mM imidazole). Protein was eluted with 4 column volumes of elution buffer (57 mM NaH2PO4 pH 7.9@RT, 30 mM NaCl, 235 mM imidazole). Eluted protein was dialyzed to completion in 1xPBS and snap-frozen in liquid nitrogen for long-term storage at −80°C. Complete plasmid sequences are provided in Supp File 1.

### Covalent Bonding of SARS-CoV-2 Spike RBD to SpyCage

Purified SpyCage was dialyzed into 1xPBS with 1 mM DTT, and then mixed with purified SARS-CoV-2 Spike RBD either with (RBD+SpyTag) or without (RBD only) at a 1.2:1 molar ratio of RBD to SpyCage monomer in a buffer consisting of 1xPBS and 1mM DTT. The binding reaction was allowed to go to completion by incubation for 3 hours at RT. The extent of SpyCage saturation was assessed by SDS-PAGE as previously described^43^. The binding reaction was then dialyzed into 1xPBS and stored at −80C until use in immunization efforts.

### Cryo-EM specimen preparation and data collection

Purified apo cage protein complex based upon I3-01^41,42^ was first assessed by negative staining to check sample quality and concentration before preparing TEM grids for data collection. Briefly, a 3.5 ul aliquot was applied to a glow-discharged Cu-grid coated with a thin film of continuous carbon, washed, stained with 0.75% w/v uranyl formate for 15 sec, blotted, air-dried, and loaded on EFI Tecnai G2 Spirit BioTwin microscope (120 kV) for imaging.

TEM grids (QUANTAFOIL R2/1; QUANTAFOIL, Germany) were plasma cleaned using a PELCO Glow Discharge System (Ted Pella, Redding CA). Aliquots of 3.5 ul of the apo cage sample at approximately 0.1 mg/ml were applied to the grids, blotted for 2 sec, and then plunge-frozen in liquid ethane using a vitrification robot (Vitrobot, Thermo Fisher). Grids were stored in liquid nitrogen until the date of screening and data collection. Data was acquired on a Thermo Fisher Titan Krios electron microscope (300 kV) equipped with Falcon 3EC direct detection camera. EPU software (V 2.13.0.3175REL) was used to set up data acquisition at a nominal magnification of x59,000 and a physical pixel size of 1.11 Å/pixel. A total of 1,220 micrographs were recorded as movies (stacks of 39 frames) at an exposure rate of 1.15 e/Å^2^/frame and a total exposure time of 69.8 s. The nominal defocus range of −1.2 to −3.0 μm was applied during data collection.

### Cryo-EM Image processing

Image analysis was performed using cryoSPARC software package (v3.3.2)^44^. Aligned movie stacks were generated from raw micrographs after correcting for stage drift and anisotropic motion using patch motion correction. Parameters of the contrast transfer function (CTF) were estimated for each aligned movie in patch mode. Manually selected 283 particles from 11 micrographs were used to train a Topaz model for particle picking; a box of 420×420 pixel size was used for particle extraction^45^. The trained model was applied to pick 129,792 particles from 1,202 micrographs. Further cleaning of the data using 2D classification resulted in 63,430 particles for subsequent data processing. A map from an *ab initio* model (generated using 10,000 particles) along with the selected clean particles were subjected to homogenous refinement in cryoSPARC. Local motion correction^46^ of the refined particles followed by homogenous refinement with higher-order CTF terms enabled (including beam-tilt, spherical aberration, trefoil and tetrafoil) and icosahedral symmetry (I1) enforced resulted in a final map at 3.4 Å resolution.

### Cryo-EM Model building

The initial model of the apo cage monomer was extracted from the published I3-01 model^41,42^. The monomer model was manually fitted into the 3.4 Å map in ChimeraX^47^; a full icosahedral model of apo cage was generated from the asymmetric unit. PHENIX real-space refinement was used to refine the model against the sharpened map with non-crystallographic symmetry parameters applied^48^. The refined model was visually inspected in Coot and validated by MolProbity^49,50^. All figures of the protein structure and cryo-EM map were created using ChimeraX.

### Culture of SARS-CoV-2

The SARS-CoV-2/USA/WA1/2020 isolate was received from The World Reference Center for Emerging Viruses and Arboviruses (WRCEVA), University of Texas Medical Branch at Galveston (UTMB). The virus was obtained at passage 4 and was sub-cultured once on Vero E6/TMPRSS2 cells (Japanese Collection of Research Bioresources Cell Bank). All titrations of virus stocks and tissue homogenates were performed on Vero E6 cells (ATCC) cultured in Dulbecco’s modified Eagle Medium (Cytiva) supplemented with 10% FBS, 4 mM L-glutamine, 1 mM sodium pyruvate (Corning), 1X non-essential amino acids and 1X antibiotic and antimycotic (Corning) at 37° C with 5% CO_2_. For culture of the VeroE6/TMPRSS2 cells 1 mg/mL geneticin was added to the media, and the FBS was reduced to 5%. To determine the titer of viral stocks, the tissue culture infectious dose 50% (TCID50) was determined by inoculating cells grown in 24-well plates with serial dilutions of the virus. The plates were incubated at 37°C with 5% CO_2_ and scored for cytopathic effect at 96 hours post-infection. The TCID_50_ was then calculated using the method of Reed and Muench^51^.

### Vaccination and Challenge Experiments

Equal numbers of male and female, six to eight-week-old Syrian hamsters (HsdHan:AURA, Envigo, Haslett, MI) were used for all studies. After acclimatization, animals were implanted with a subcutaneous transponder chip (Bio Medic Data Systems), and a pre-vaccination blood sample was collected. For intranasal vaccination and virus inoculation, animals were sedated and intranasally inoculated with a vaccine candidate (70 ul in 1xPBS) or SARS-CoV-2 (100 ul in DMEM). For all experimental procedures, hamsters were sedated with 150 mg/kg ketamine, 7.5 mg/kg xylazine, and 0.015 mg/kg atropine via intraperitoneal injection. After completion of the procedure, hamsters were given 1 mg/kg atipamezole subcutaneously. For tissue collection and at the end of each study, hamsters were humanely euthanized via CO_2_ asphyxiation.

#### Trial 1: Evaluation of immunogenicity and efficacy of the SpyCage-RBD vaccine candidate

To evaluate the immunogenicity and efficacy of the SpyCage RBD vaccine, groups of hamsters (n=14/group) were intranasally vaccinated with PBS (mock), SpyCage (15 ug), SARS-CoV-2 RBD (10 ug), or SARS-CoV-2 RBD (10 ug) bound to SpyCage (15 ug, “RBD+SpyCage”). Animals received a primary (1°) vaccination and a secondary (2°) vaccination 28 days later. The vaccine was administered without adjuvants. Blood samples were collected via gingival vein from 6 animals (3 males and 3 females) per group on days 14, 26, 42, and 55 post-1° vaccination. Blood samples were centrifuged at 1000 *x g* for 10 minutes at RT, and serum was collected and stored at −20°C. On day 56 post-1° vaccination, all animals were intranasally inoculated with 10^5^ TCID_50_ SARS-CoV-2/USA/WA1/2020. On days 3 and 6 post-infection (day 59 and 62 post-1° vaccination), lung and nasal turbinate tissues were collected (n=4/group, 2 males and 2 females) and stored at −80°C. The remaining six hamsters were monitored for weight loss until day 14 (day 70 post-primary vaccination).

#### Trial 2: Assessment of the requirement for grafting of RBD to SpyCage

Groups of hamsters (n=18/group) were intranasally vaccinated with PBS (mock), SpyCage (15 ug), SARS-CoV-2 RBD (10 ug), SARS-CoV-2 RBD without SpyTag (10 ug) mixed with SpyCage (15 ug) (*i.e*., RBD could not covalently bond to SpyCage, “RBD|SpyCage”), and SARS-CoV-2 RBD (10 ug) grafted to SpyCage (15 ug, “RBD+SpyCage”). The vaccination and blood collection protocols were the same as in the initial study, and on day 56 post-1° vaccination, animals were challenged with 1000 TCID_50_ of SARS-CoV-2. On days 3, 5, and 7 post-challenge (days 59, 61, and 63 post-primary vaccination), lung and nasal turbinates were collected (n=4/group (2 males and 2 females)). One lung lobe was fixed with 10% v/v normal buffered formalin and the remaining lung lobes and nasal turbinates were stored at −80°C. The remaining six hamsters/group were monitored for weight loss until day 14 post-SARS-CoV-2 challenge (day 70 post-primary vaccination). All animals were euthanized on day 15 post-SARS-CoV-2 challenge.

### Viral titration of tissue samples

Collected lungs and nasal turbinates were homogenized in 2% FBS-DMEM containing 2X antibiotic and antimycotic using an Omni tissue homogenizer. The homogenates were centrifuged at 1000 *x g* for 10 minutes at 4°C and the supernatant was titrated to determine the tissue culture infectious dose 50% (TCID50) on Vero E6 cells as previously described^52^.

### Microneutralization Assay

To determine titers of neutralizing antibodies, microneutralization assays were performed on Vero E6 cells as previously described^52^.

### ELISA

To assess the levels of RBD-binding IgG and IgA antibodies, ELISA assays were performed according to a protocol generously provided by Dr. Sabra Klein, Johns Hopkins, School of Public Health^53,54^.

### Histopathology

Formalin-fixed lung samples were processed and stained with haematoxylin and eosin as previously described^54^. Slides were scored by a board-certified veterinary pathologist using established methods^55^. Each animal was scored for the extent of lesions (0-4), alveolar damage (0-3), bronchial damage (0-3), blood vessel damage (0-3), hemorrhage (0-2), and type II pneumocyte hyperplasia (0-2). For each animal, a total pathology score was obtained by calculating the sum of scores.

### Biocontainment and Animal Care and Use

All experiments using SARS-CoV-2 were conducted in an animal biosafety level 3 enhanced laboratory. This facility is approved by the US Department of Agriculture and the Centers for Disease Control and Prevention. All animal studies were conducted in compliance with the Institutional Animal Care and Use Committee under protocol number 202001440.

### Statistical Analysis

Prism GraphPad (v9.0) was used to perform all statistical analyses with p<0.05 considered significant. Weight loss and viral titers at each time point were evaluated for normality by D’Agostino & Pearson test. For data sets that passed the normality test, one-way ANOVA with post-hoc Tukey’s test was performed. When data sets did not pass the normality test, Kruskal-Wallis tests with a post-hoc Dunn’s multiple comparison test were performed. Histopathological scores were also compared using non-parametric Kruskal-Wallis tests with a post-hoc Dunn’s multiple comparison.

## RESULTS

### Cryo-EM reconstruction and refinement of an atomic model of the apo cage scaffold

To establish a robust, multimeric, spherical protein-based scaffold for intranasal immunizations that would mimic the size of a viral particle, we selected a wireframe dodecahedron based upon the previously described I3-01 protein, which was designed to self-assemble from 60 monomers^41,42^. To validate this scaffold structurally, we assessed a purified sample of this apo cage complex by electron microscopy. Samples were first quality controlled by negative staining with uranyl formate to assess particle integrity and concentration, and were then vitrified on gold grids for cryo-EM data collection on our home-source Titan Krios electron microscope equipped with a Falcon 3EC direct detection camera. Data processing and all aspects of the cryo-EM workflow were conducted in cryoSPARC software (v.3.3.2)^44^.

To create an experimentally determined high-resolution model, we collected a cryo-EM dataset with 1,220 recorded movies to yield 1,202 processed micrographs with good-quality ice. These were used to auto-pick and extract 129,792 particles using a box size of 420×420 pixels (calibrated pixel size = 1.11 Å) (**Fig. 1A**, **Supplemental Table 1**). Then 2D classification was used to clean up the data by removing junk particles (**Fig. 1B**) to produce a total of 63,430 particles for cryo-EM map reconstruction and refinement while imposing icosahedral symmetry. The refinement produced a cryo-EM map at an average resolution of 3.4 Å and an estimated local resolution in the range of 3.0 Å to 5.0 Å (**Fig. 1C**). Map resolution was determined based on the gold-standard criterion that applies a Fourier Shell Correlation (FSC) cutoff value of 0.143 (**Fig. 1D**)^56^. The 3.4 Å resolution map showed the typical features of the designed apo cage with an average diameter of 25 nm and the trimeric protein units occupying the vertices of the pentameric faces of the dodecahedron. The I3-01 design PDB of this protein structure was modified and used to initiate model building^41,42^. The map density was clear enough to build the atomic structure using the published model of I3-01 to provide the starting coordinates. Real-space refinement of the icosahedral model in PHENIX resulted in a 3.4 Å resolution model (**Fig. 1E**) with a cross-correlation value for model vs. map of 0.77 (CC masked, **Supplemental Table 1**). The geometrical parameters of the refined model checked by MolProbity revealed a good quality model with a MolProbity score of 1.5, with approximately 98% of residues in the favored region and no residues in the disallowed region of the Ramachandran plot. Quality-check parameters of the model and map-model agreement are listed in **Supplemental Table 1**. The 3.4 Å resolution map showed clear density for most side chains of the amino acids constituting the apo cage protein (aa 22-222) (**Fig. 1F)**. However, as expected for structures solved in the range of 3.0 to 4.0 Å resolution^57^, directionality of some carbonyl groups could not be resolved unambiguously.

**Figure 1.**
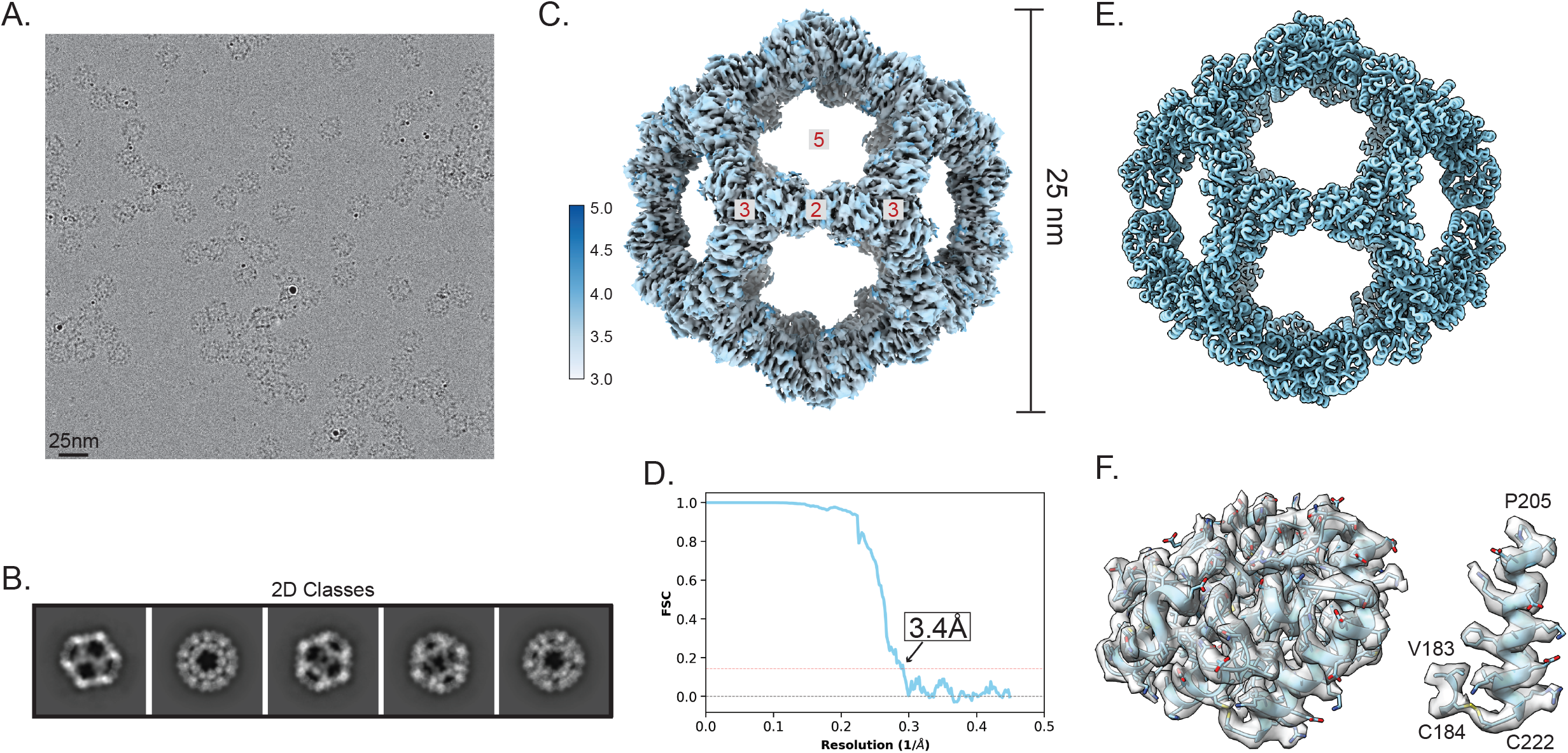
Single-particle analysis and Cryo-EM reconstruction of the apo cage used for immunizations. (A) A representative cryo-EM micrograph of apo cage particles (scale bar, 25 nm) is provided. (B) Examples of representative class averages from a 2D classification of the particles extracted from cryo-EM micrographs are provided. (C) A reconstructed icosahedral map of the apo cage structure is colored according to the estimated local resolution; color key is shown to the left of the map. Red numbers in gray boxes on the structural model indicate the two-, three- and five-fold symmetry axes of the dodecahedron. Apo cage particles have an approximate diameter of 25 nm. (D) A fourier shell correlation (FSC) curve of the reconstructed map using gold-standard refinement in cryoSPARC is presented. An approximate map resolution of 3.4 Å based on 0.143 FSC cutoff is indicated. (E) An atomic model of the apo cage was built by applying icosahedral symmetry in ChimeraX to an asymmetric unit fitted to the density of the map shown in (C). (F) Left: A portion of the map covering a single I3-01 monomer is rendered as a transparent surface, with the fitted model (aa 22-222) shown as a light blue cartoon with side chains represented as sticks. Right: A close-up view of residues Val183, Cys184 and the C-terminal helix (aa 205-222) showing clear density of the assigned sidechains is shown with the map contoured at level 0.9 in ChimeraX. The quality of density is sufficient to observe the disulfide bond between Cys184 and the C-terminal Cys222.

Comparing the solved cryo-EM structure and that from the computationally designed I3-01 model^41,42^ showed that the two structures are almost identical [r.m.s. values of 0.57 Å for 200 Cα atoms and 1.29 Å for all non-hydrogen atoms], but the sidechain atoms of surface residues had minor differences between our experimental and the computationally designed structures.

### Covalent Bonding of SARS-CoV-2 Spike RBD to SpyCage

As the wireframe cage scaffold was robust, spherical, symmetrical, and could outwardly present up to 60 fused proteins-of-interest, we selected it for further modification for antigen display. Because the genetic fusion of antigens directly to protein-based scaffolds can influence expression levels, solubility, and purification conditions needed, we leveraged the SpyTag/SpyCatcher system to covalently link antigens-of-interest to the scaffold following its purification^43^. This approach enables substantial versatility to load different proteins and their variants without modifying the scaffold itself and has been used for a variety of viral and parasitic pathogens to enhance immune responses^29–31,34–36,58^. To this end, we appended a SpyCatcher domain with a flexible linker to the N-terminus of I3-01 so that all 60 subunits bear this capture domain (schematic in **Fig. 2A**). We observed that this self-assembling scaffold fused with SpyCatcher capture domains displayed excellent solubility and stability profiles when expressed in *E. coli*, as it expressed to high levels and did not precipitate in standard laboratory conditions (**Fig. 2B**). A comparable arrangement has been described for the mi3 variant of I3-01, which also exhibited favorable display properties^30^. This scaffold, which we have termed SpyCage, was advanced for all immunization studies presented here.

**Figure 2.**
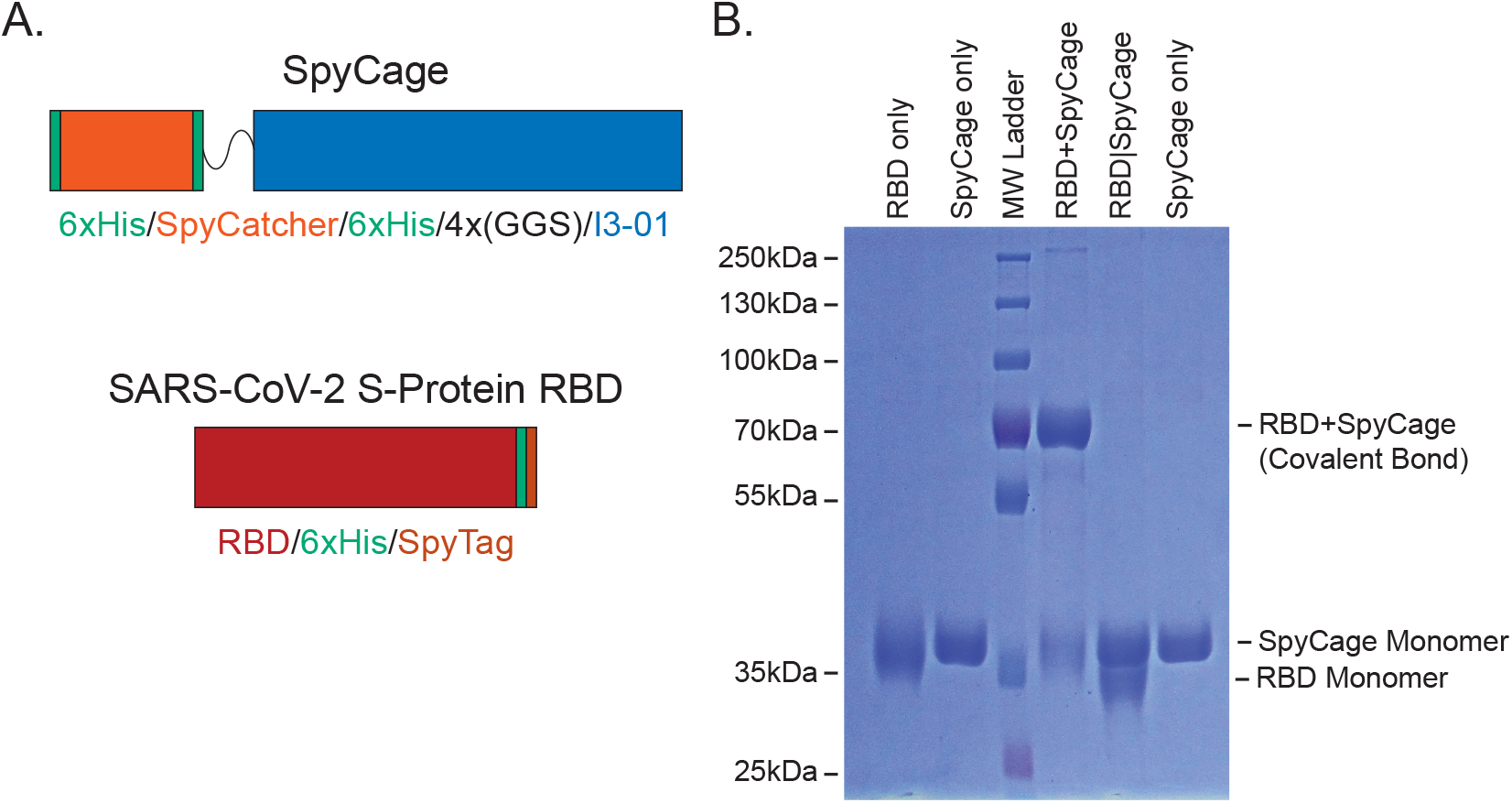
Display of RBD via the SpyCage scaffold. (A) A schematic of the SpyCage scaffold illustrates 6xHis purification tags, the SpyCatcher capture domain, a flexible linker, and a C-terminal I3-01 variant used to create the self-assembling protein wireframe platform. A schematic of RBD with a C-terminal 6xHis tag and with/without a C-terminal SpyTag is shown. (B) SDS-PAGE of SpyCage, RBD, SpyCage+RBD, and SpyCage|RBD preparations. The covalent bonding (“grafting”) of RBD to SpyCage is evident when RBD bears a SpyTag, but not in its absence as per a mobility shift. SpyCage approaches saturation with RBD at a 1-to-1.2 molar ratio of SpyCage-to-RBD, as has been seen with other antigens of comparable shape and mass.

The use of SpyTag/SpyCatcher elements permits the versatile loading of antigens that are independently expressed in an ideal expression system for that protein, thus ensuring that proper post-translational modifications and processing events occur. Here, using 293F suspension cells, we produced the receptor-binding domain (RBD) of SARS-CoV-2 Spike protein with a C-terminal 6xHis tag, and either with or without an additional C-terminal SpyTag (**Fig. 2A**). Secreted protein was purified from the culture supernatant to >99% purity via Ni-NTA affinity chromatography (**Fig. 2B**). Mixing of SpyCage with RBD (“RBD+SpyCage”)at a 1 to 1.2 molar ratio in 1xPBS led to the formation of a covalent bond that could be detected by a mobility shift by SDS-PAGE with >95% saturation of the scaffold with RBD (**Fig. 2B**). This molar ratio consistently achieves this maximal degree of saturation for RBD and other comparably sized, globular antigens (data not shown). In contrast, when SpyCage was mixed with RBD lacking a SpyTag (“RBD|SpyCage”), no covalent linkage formed between the RBD and SpyCage, as intended (**Fig. 2B**). This second combination creates an admixture (“RBD|SpyCage”) that permits testing of the effect that covalent bonding of the antigen to the scaffold has upon efficacy. From this we concluded that SpyCage is a stable and saturable antigen display platform capable of presenting antigens-of-interest in a versatile mix-and-go format. As SpyCage has similar structural properties as a virus particle and displays 5-15 copies of an antigen on a single face of the scaffold (up to a total of 60 antigens per particle), we hypothesized SpyCage grafted with RBD would be a greatly improved vaccine candidate. As there is an urgent need for the development of vaccines inducing mucosal immunity, we proceeded to evaluate SpyCage as an intranasal vaccine.

### RBD grafting to SpyCage is required to induce an antibody response

To assess the immunogenicity of RBD+SpyCage as an intranasal vaccine candidate (*i.e*., Trial 1), hamsters were given a 1° and 2° intranasal vaccine consisting of PBS (mock), SpyCage, RBD, or RBD+SpyCage. The two vaccine doses were administered 28 days apart and the vaccines were formulated without adjuvants. Serum samples were collected prior to each vaccination and the day before viral challenge. To assess the antibody response, levels of RBD-binding IgG antibodies in the serum were quantified by ELISA and neutralizing antibodies were assayed by a microneutralization assay. We found that only the RBD+SpyCage vaccinated animals developed an IgG antibody response (**Fig. S1A**). On day 28, one animal had IgG antibodies against RBD, while on day 55, 4/6 animals given RBD+SpyCage developed an antibody response (**Fig. S1A)**; however, while these antibodies were able to bind RBD, they did not exhibit neutralizing activity (**Fig. S1B**).

Subsequently, we conducted an expanded vaccine study (designated Trial 2) to determine if grafting of RBD directly to SpyCage through covalent bonding was required to induce an antibody response. Animals were vaccinated according to the same regimen; however, an additional group in which RBD without SpyTag was mixed with SpyCage (“RBD|SpyCage”) was included to test the importance of the bonding of RBD to SpyCage. Therefore, the experimental groups consisted of animals given the following vaccines: 1) Mock (PBS), 2) RBD, 3) SpyCage, 4) RBD mixed with but not bound to SpyCage (RBD|SpyCage), and 5) RBD grafted to SpyCage (RBD+SpyCage). In both vaccination studies, animals were monitored for 7 days post-1° and 2° vaccination for adverse effects. None of the animals exhibited weight loss or clinical signs, indicating the vaccine was well-tolerated (data not shown).

Evaluation of the antibody response by ELISA and microneutralization assay (**Fig. 3**) showed that none of the mock, SpyCage, RBD alone, or RBD|SpyCage immunized animals developed RBD-directed antibodies on day 28 or 55. In contrast, animals vaccinated with RBD+SpyCage developed IgG antibodies against RBD on day 28 post-1° vaccination (2/6 animals positive) and on day 55 (5/6 animals positive) (**Fig. 3A**). While most animals in the RBD+SpyCage group developed IgG antibodies by day 55, only one animal developed an IgA antibody response (**Fig. 3B**). This was one of the two animals that developed IgG antibodies early (day 28) and was the animal with the highest IgG antibody titer on day 55 (1: 12800) (**Fig. 3A**). In addition, the serum from this animal exhibited neutralizing activity (**Fig. 3C**), while none of the other animals developed a neutralizing antibody response. Thus, while we did not observe 100% seroconversion, the RBD+SpyCage vaccine candidate was reproducibly capable of inducing IgG antibodies. As one animal developed both a neutralizing antibody response and IgA antibodies indicative of mucosal immunity, our findings indicate that, with additional modifications to enhance immunogenicity, intranasal vaccination with RBD+SpyCage could induce mucosal immunity. Furthermore, these findings demonstrate the RBD+SpyCage vaccine provides a substantial boost in antibody responses compared to RBD alone or an RBD|SpyCage admixture where the covalent bond needed for grafting cannot form. To address the immunogenicity of SpyCage itself, we also evaluated the antibody response to SpyCage by ELISA (**Fig. 3D)**. All animals given a vaccine containing SpyCage developed IgG antibodies directed towards the scaffold. While there were no statistically significant differences between different vaccine groups that received the SpyCage, the RBD|SpyCage admixture increased the antibody response 2-fold compared to animals given SpyCage alone, and RBD+SpyCage further increased the response by 2-fold (*i.e*., 4-fold relative to SpyCage alone). These findings indicate that grafting the antigen to SpyCage enhances the antibody response to both the scaffold and the antigen.

**Figure 3.**
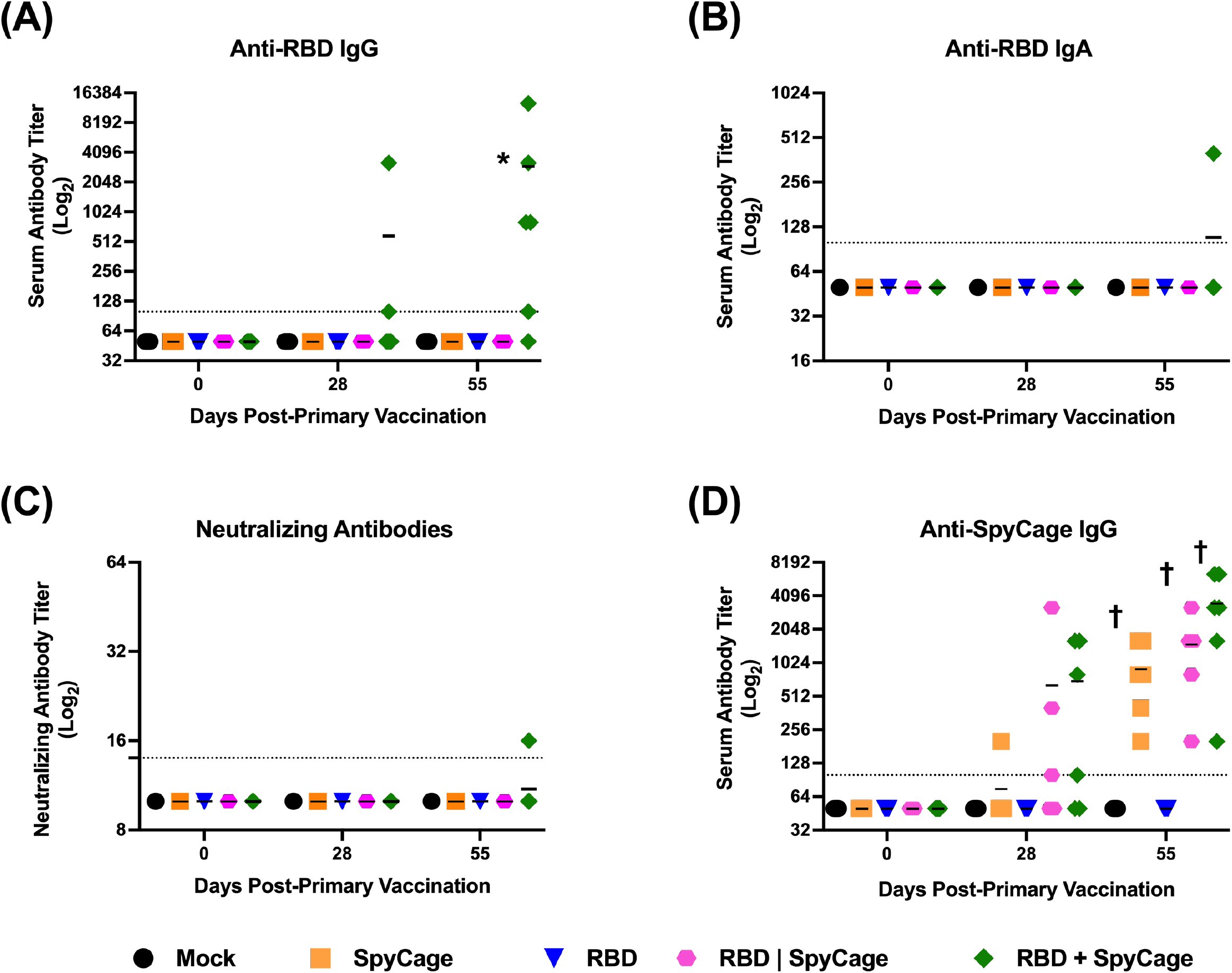
Binding and neutralizing antibody responses to intranasal vaccination with RBD+SpyCage. Antibody titers were measured in serum samples on days 0, 28, and 55, prior to primary vaccination, boost vaccination, and viral challenge, respectively. Plotted are (A) anti-RBD IgG, (B) IgA titers, (C) neutralizing antibody titers against SARS-CoV-2, and (D) IgG antibody titers against the SpyCage scaffold. * significantly different from all other groups by Kruskal-Wallis with Dunn’s multiple comparison. † significantly different from mock and RBD groups.

### Intranasal vaccination with RBD+SpyCage enhances clearance of SARS-CoV-2 from the respiratory tract

To assess the efficacy of the RBD+SpyCage vaccine candidate, in both vaccine trials of this study, we assessed whether RBD+SpyCage enhanced viral clearance and reduced clinical illness. In the second trial, we also evaluated the effect of vaccination on reducing lung pathology. In Trial 1, vaccinated hamsters were challenged with 10^5^ TCID50 SARS-CoV-2 on day 56 post-1° vaccination (day 28 post-2° vaccination). After viral challenge, animals were monitored for weight loss for 14 days, and on days 3 and 5 post-infection (p.i.), lung and nasal turbinate samples were collected from a subset of animals (n=4/group/timepoint) (**Fig. S2**). After viral challenge, animals in all experimental groups lost weight (**Fig. S2A**) and there were no statistically significant differences between the groups; however, the animals that received RBD+SpyCage had reduced weight loss, and by the end of the study, these animals exceeded their pre-challenge weight. When we evaluated viral titers in the lungs and nasal turbinates, on day 3 p.i. all experimental groups had high titers of replicating virus in these tissues with no significant differences between groups. On day 6 p.i., viral titers in both tissues were reduced for all groups; however, while the mock and SpyCage-vaccinated animals had replicating virus in the nose and lungs, no replicating virus was recovered from the RBD only and RBD+SpyCage vaccinated animals (**Fig. S2 B,C**). These findings suggested vaccination with RBD or RBD+SpyCage facilitated viral clearance.

While vaccination with either RBD only or RBD+SpyCage reduced viral load on day 6, only the RBD+SpyCage vaccinated animals developed an IgG antibody response. We therefore performed a second, expanded vaccination study (i.e., Trial 2) to evaluate if grafting of RBD to SpyCage via covalent bonding was required for protection. Here we repeated the vaccination study with an additional group of animals that were vaccinated with an admixture of RBD and SpyCage where the covalent bond needed for grafting could not form (RBD|SpyCage). As the 10^5^ TCID_50_ challenge dose used in the initial study is 2-3 orders of magnitude higher than the estimated infectious dose for humans (*i.e*., 100-1000 infectious units)^59,60^, we reduced the challenge dose to 1000 TCID50 of SARS-CoV-2. This challenge dose was previously shown to induce weight loss in hamsters^61^, which we also verified with our virus stock (**Fig. S3)**. Finally, to evaluate the dynamics of viral clearance more comprehensively, we also modified the time points of tissue collection such that tissues were collected on days 3, 5, and 7 p.i. After viral challenge, the animals in all experimental groups lost weight, with peak weight-loss at day 6 or 7. However, while there were no statistically significant differences in weight loss between experimental groups (**Fig. 4A**), consistent with our first study, we also observed that animals vaccinated with RBD+SpyCage trended toward reduced weight loss compared to the other groups (**Fig. 4A**). We next evaluated viral replication in the nasal turbinates and lungs. On day 3 p.i., mean viral load in the nasal turbinates and lungs for all groups were comparable, with titers greater than 10^5^ and 10^6^ TCID50/gm in each tissue, respectively (**Fig. 4 B,C**). However, on day 5 p.i. the mean viral titer in the nasal turbinates for the RBD+SpyCage group was significantly lower (25 TCID50/gm) compared to the other experimental groups (titer range: 881 – 3955 TCID50/gm) (**Fig. 4B**). Similarly, in the lungs, RBD+SpyCage vaccinated animals also had significantly lower titers (1183 TCID50/gm) (titer range for other experimental groups:11,988 – 59,356 TCID50/gm) (**Fig. 4C**). On day 7 p.i., replicating virus was not detected in the nasal turbinates or lungs from any of the experimental groups (**Fig. 4 B,C**). Therefore, while viral titers on days 3 and 7 in the nasal turbinates and lungs were comparable for all groups, on day 5 viral titers in the RBD+SpyCage vaccinated animals were more than 10-fold lower, indicating that the RBD+SpyCage vaccinated animals more rapidly cleared SARS-CoV-2 from both the upper and lower respiratory tract.

**Figure 4.**
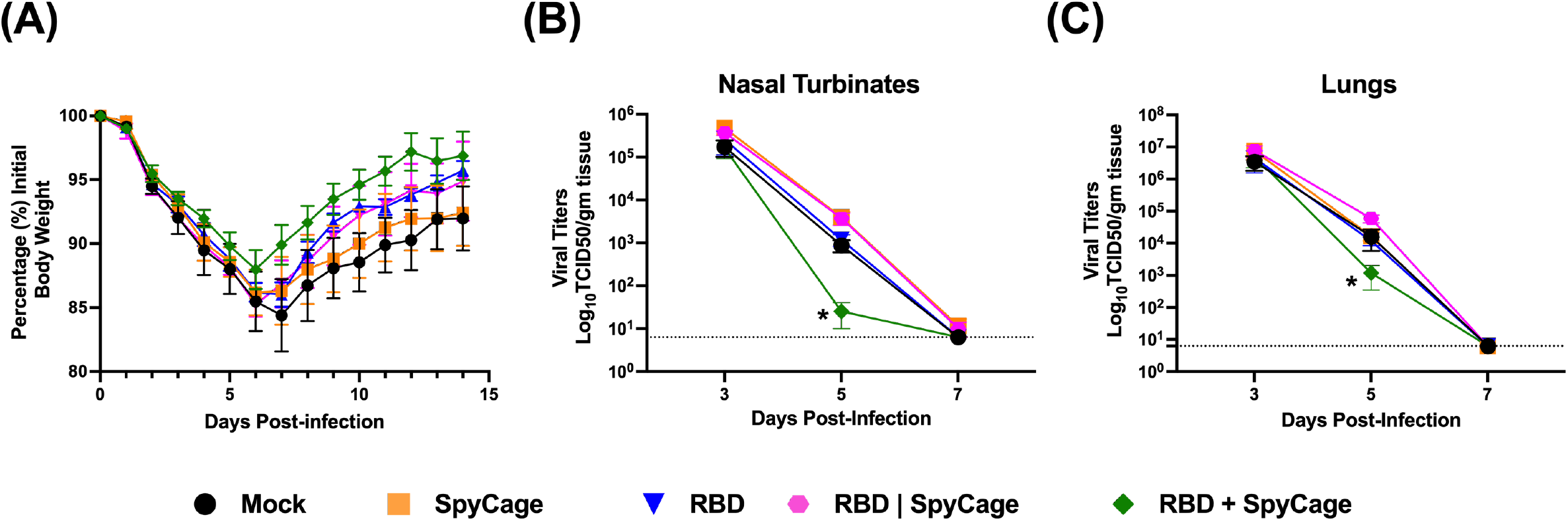
Weight loss and viral titers in the nasal turbinates and lungs after SARS-CoV-2 challenge of vaccinated hamsters. After viral challenge, hamsters were monitored for (A) weight loss, and viral titers were evaluated in (B) nasal turbinates and (C) lung tissues on days 3, 5 and 7 post-infection. *significantly different from RBD and RBD|Spycage. **significantly different from RBD|SpyCage. Non-parametric Kruskal-Wallis test with Dunn’s multiple comparisons were used to determine significant differences.

Last, we performed a histopathology analysis to determine if the RBD+SpyCage vaccinated animals had reduced lung inflammation and damage. Lung tissue sections were blinded and scored for the extent of lesions, alveolar, bronchial, and blood vessel damage, as well as hemorrhage and type II pneumocyte hyperplasia. These scores were then combined to give a total pathology score. Representative images of lung pathology and inflammation from each group are shown in **Fig. 5A**. The largest differences in pathology scores were observed in the total pathology score and the extent of lesions (**Fig. 5 B,C)**, with additional scores reported in **Fig. S4**. On day 3, all groups exhibited similar pathology. For the mock vaccinated animals, the total pathology score and extent of lesions peaked on day 5 and then declined on day 7. The RBD+SpyCage vaccinated animals exhibited the lowest total pathology and extent of lesions scores compared to all other groups on both days 5 and 7. Animals receiving SpyCage, RBD, or the SpyCage|RBD admixture had intermediate scores between the mock and SpyCage+RBD groups on day 5, and had pathology scores comparable to mock infected animals on day 7. While the pathology scoring shows a trend towards reduced pathology with the SpyCage+RBD group, this difference was not statistically significant due to the limited number of animals used at each time point. Future studies can leverage these observed effect sizes to establish expanded group sizes to determine if these promising trends are maintained.

**Figure 5.**
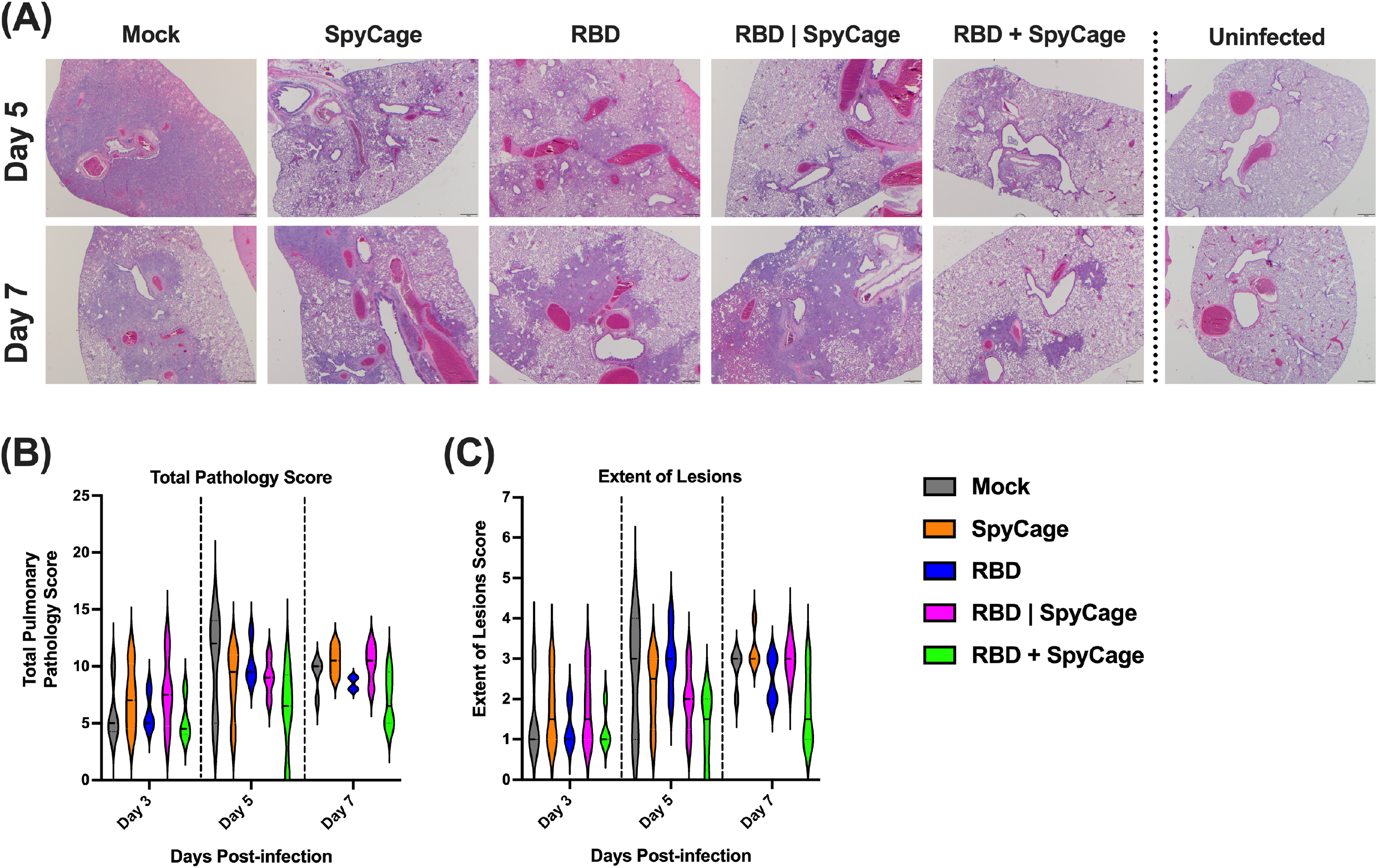
SARS-CoV-2 induced lung pathology in vaccinated hamsters. On days 3, 5, and 7, post-infection, lung tissues were processed for H&E staining and scored by a veterinary pathologist. Panel (A) displays representative images from each group of hamsters on days 5 and 7 post-infection. This panel also includes images of uninfected hamster lung tissues (far right panels). Panels (B) and (C) display total pathology scores and the extent of lesions scoring, respectively.

## DISCUSSION

The development of efficacious intranasal vaccines has the potential to prevent SARS-CoV-2 infections and reduce transmission^62^. To date several pre-clinical intranasal vaccine candidates have been developed ^63^; however, most candidates are live-attenuated or viral vector vaccines. Due to safety concerns, the administration of these vaccines is often limited to healthy adults (18-55 years old) and/or older children. In contrast, recombinant protein or inactivated vaccines are widely used in individuals of all ages. Therefore, we sought to develop a recombinant protein-based intranasal vaccine.

Because immune responses can be enhanced when antigens-of-interest are displayed on a scaffold, we established design criteria for an intranasal vaccine candidate. These criteria included 1) rigid bodies, 2) a spherical shape of ~20-30 nm in diameter, and 3) genetically accessible N- and/or C-termini presented in an outward-facing manner. I3-01 met all these criteria and was selected as the strongest candidate. To date, only a lower resolution cryo-EM reconstruction and a computational model of I3-01 have been published^41^. Therefore, we used both negative stain TEM and cryo-EM to resolve the I3-01-based scaffold (apo cage) to a 3.4A average resolution and validated that the experimentally derived atomic model closely matched the computationally designed protein (**Fig. 1**). We then proceeded with using I3-01 to create SpyCage by further modifying I3-01 to bear an N-terminal SpyCatcher domain with a 12 amino acid flexible linker to reduce steric hindrance and permit greater saturation of antigens. As anticipated, the SpyTag/SpyCatcher system enabled rapid, covalent linkage of RBD to SpyCage (RBD+SpyCage) to near saturation as seen by a mobility shift by SDS-PAGE (**Fig. 2**). Importantly, the RBD+SpyCage preparation remained highly soluble and stable over time, which further supports its feasibility as a vaccine candidate.

Subsequently, we assessed the immunogenicity of the RBD+SpyCage vaccine in the gold standard Syrian hamster model (*i.e*., Trial 1). Animals were given an unadjuvanted prime-boost intranasal vaccination 28 days apart, and then challenged 28 days later (day 56 post-primary vaccination) with SARS-CoV-2. On day 26 post-primary vaccination, 1 of 6 RBD+SpyCage vaccinated hamsters developed serum IgG antibodies, and by day 55, most animals (4/6) had a measurable IgG response. None of the animals developed neutralizing antibodies (**Fig. S1**). Following viral challenge, all animals, regardless of vaccination status, lost weight, although there was a trend towards reduced weight loss and earlier recovery in the RBD+SpyCage vaccinated animals (**Fig. S2**). Based on these outcomes, we next sought to determine the properties of the vaccine that enhanced immunogenicity.

Studies on intramuscular vaccination have shown that presenting viral antigens on the surface of particles enhances the immune presentation and protective efficacy of vaccines^64^. Therefore, we expanded upon our initial study and compared RBD+SpyCage (in which RBD is covalently bound to the scaffold) to an RBD|SpyCage admixture lacking this covalent attachment. Covalent grafting was shown to be a requirement for immunogenicity, as 2/6 and 5/6 animals given RBD+SpyCage developed an IgG response on days 28 and 55, respectively against RBD (**Fig 3**). The one animal that exhibited the highest IgG titers on day 28 and day 55, also developed an IgA response, and these antibodies exhibited neutralizing activity. In contrast, animals vaccinated with RBD|SpyCage (*i.e*., RBD mixed with, but not covalently bound to SpyCage) did not develop an antibody response to RBD. Further studies are warranted to explore strategies to enhance the immunogenicity of RBD+SpyCage to match the neutralizing IgG and IgA response we observed in this one animal.

As a final component of our antibody analyses, we evaluated the response generated against the SpyCage scaffold. All animals that received SpyCage as a component of the vaccine developed antibodies directed towards the scaffold. Our results do not suggest this immunity interfered with the immune response to RBD, as boost vaccination increased the response to RBD. However, in future studies, if immunity against the scaffold interferes with immunogenicity for other vaccine antigens, alternative scaffolds can be evaluated for subsequent intranasal vaccinations^65^. Importantly, assessment of the immune response in hamsters is currently hampered by a limited number of species-specific reagents. Reagents to assess mucosal and cellular immunity are lacking. Using our existing protocol, we previously performed ELISA on matched convalescent hamster serum and nasal wash samples^54^. While we detected serum IgA antibodies, we could not detect antibodies in the nasal wash (data not shown). Therefore, as new reagents are developed for hamsters, it will be important to expand the immunological assessment of vaccine responses.

Covalent grafting to SpyCage was also a requirement for vaccine efficacy. Upon SARS-CoV-2 challenge in Trial 2, all animals lost weight; however, compared to the other groups, RBD+SpyCage vaccinated animals had a trend towards reduced weight loss and reduced lung pathology. This was associated with significantly reduced levels of replicating virus in the respiratory tract on day 5, indicating rapid viral clearance in the RBD+SpyCage group relative to the RBD|SpyCage group **(Fig 4)**. Future studies could pinpoint a more comprehensive view of the dynamics and changes in viral clearance. Collectively, the induction of non-neutralizing RBD-binding antibodies in association with accelerated viral clearance and trends towards reduced disease severity and pathology suggests alternative antibody-mediated mechanisms (*e.g*., antibody-dependent cell-mediated cytotoxicity (ADCC) or antibody-dependent cellular phagocytosis (ADCP)), and/or that T cell mediated immunity contributed to protection. To our knowledge, this is the first report of a scaffolded antigen being used as an intranasal protein-based vaccine. Other groups have used the I3-01 scaffold successfully as a vaccine platform to display antigens for influenza, SARS-CoV-2, and *Plasmodium*, but have only explored intramuscular administration^30,37,58,66–69^. Several groups have expressed the RBD or S-protein on I3-01 and evaluated the immunogenicity of these scaffolded antigens as intramuscular vaccines in animal models^37,58,66–68^. In these studies, the vaccine candidates were administered with an adjuvant (*e.g*., Addavax, Alum, CpG), and potent neutralizing antibody responses were induced in mice, hamsters, pigs, or non-human primates^37,58,66–68^. Given both the route of administration and the inclusion of adjuvants, this is the expected antibody response. In comparison, we administered the RBD+SpyCage vaccine only via the intranasal route and without an adjuvant, which induced a non-neutralizing IgG antibody response. Prior studies have evaluated the protective efficacy of intramuscular vaccination with RBD grafted to I3-01 against SARS-CoV-2 challenge^37,58^. When hamsters were intramuscularly vaccinated and challenged with SARS-CoV-2, consistent with our results, all animals lost weight, and the RBD-I3-01 vaccinated animals (designated “RBD-VLP” in that study) had reduced weight loss relative to animals vaccinated with RBD alone^37^. Unfortunately, in this study, viral titers were not evaluated in lung or nasal turbinate samples after viral infection, precluding a comparison with our findings. In another study, when transgenic K18-hACE2 mice were vaccinated with a similar construct, SARS-CoV-2 Beta RBD-mi3, and challenged, all vaccinated mice survived a lethal challenge while only 20% of control animals survived. In parallel, with enhanced survival, no replicating virus was detected in the lungs of the RBD-mi3 vaccinated animals^58^. Similarly, when Rhesus macaques were vaccinated with the same construct and challenged with SARS-CoV-2, at both day 2 and 4 p.i. significantly lower titers of virus were detected in nasal swabs compared to unimmunized controls. Moreover, in RBD-mi3 vaccinated animals, replicating virus was not recovered from bronchioloalveolar lavage fluids (BAL) (i.e., 0/4), while 3/4 unimmunized controls had between 10^3^ – 10^6^ TCID50/mL of SARS-CoV-2 in the BAL^58^. In addition to I3-01, the bipartite I53-50 icosahedral scaffold consisting of 120-subunit proteins has also been decorated with either the RBD or S-protein and utilized as an intramuscular vaccine^29,70^. Consistent with intramuscular vaccination with I3-01, intramuscular vaccination with RBD or S-protein grafted to I53-50 combined with an adjuvant induced a neutralizing antibody response in mice, rabbits, or macaques ^29,70^. In these studies, only macaques vaccinated with S-protein on I53-50 nanoparticles were challenged with SARS-CoV-2. Following viral challenge, relative to unimmunized controls, vaccinated animals had reduced clinical manifestations associated with significantly reduced viral titers in the upper airways and BAL from day 1 until resolution on day 7 p.i.^29^. In contrast to these studies, when we challenged the RBD+SpyCage vaccinated animals, we did not observe an initial reduction in SARS-CoV-2 replication as all animals had similar titers on day 3 p.i.; however, the RBD+SpyCage vaccinated animals had reduced titers on day 5 indicating accelerated viral clearance. We also observed trends towards reduced weight loss and reduced pathology; however, these were not statistically significant. The reduced efficacy observed in our studies relative to intramuscular vaccination is most likely due to a lack of a neutralizing antibody response following intranasal vaccination. In our studies, we purposefully did not include an adjuvant as there are no licensed intranasal adjuvants for human use; however, future development of the intranasal RBD+SpyCage vaccine warrants the inclusion of intranasal adjuvants to enhance the quality of the antibody response and vaccine efficacy. In addition, to more closely model immunity in the human population, a recent study explored intranasal vaccination with recombinant S-protein as a booster in animals previously given intramuscular vaccines^71^. This approach enhanced protection, and based on our findings, we posit that such an intranasal boost would benefit from scaffolding the S-protein or its derivatives. Collectively, we demonstrate intranasal vaccination with RBD grafted to SpyCage induced a serum IgG response in hamsters. Upon viral challenge, this response was associated with enhanced viral clearance from both the upper and lower respiratory tract. RBD+SpyCage vaccinated animals also exhibited non-significant reductions in weight loss and lung pathology consistent with a non-neutralizing antibody response. We further show the immunogenicity and efficacy of the RBD+SpyCage vaccine required that RBD was covalently linked to the SpyCage scaffold. These studies demonstrate the potential for intranasal delivery of SpyCage scaffolded antigens as a vaccine platform, and additional vaccine development is warranted with the inclusion of intranasal adjuvants to enhance immunogenicity. Moreover, given the relative ease with which vaccine antigens can be grafted to the scaffold and the potential to induce mucosal immunity, SpyCage-derived intranasal vaccines can be developed to target other respiratory viruses, and if successful, this platform could also be used as a rapid response vaccine platform to target novel or pandemic pathogens.

## DATA AVAILABILITY

The icosahedral map of the solved apo cage structure is deposited in EM data bank under accession code EMD-27812. The apo cage atomic model is deposited in the PDB data bank under ID 8E01. <u>A validation report is provided as Supp File 2 for peer review purposes</u>.

## ACKNOWLEDGMENTS

SH, SEL, and TCS planned all experiments with assistance from DRP and AM. AM and SEL designed and purified recombinant proteins and vaccine candidates. RMR expressed the RBD protein. All cryo-EM studies were conducted by CB, IMM, and SH. Vaccination and challenge studies were conducted by DRP with assistance from DGS, CJF, AK, and TH. EHL performed histopathology and analysis of lung samples. DRP, SH, SEL and TCS prepared the final manuscript.

The molecular graphics and analyses were performed with UCSF ChimeraX, developed by the Resource for Biocomputing, Visualization, and Informatics at the University of California, San Francisco, with support from National Institutes of Health R01-GM129325 and the Office of Cyber Infrastructure and Computational Biology, National Institute of Allergy and Infectious Diseases. We would like to acknowledge Dr. David Baker from the University of Washington for providing I3-01_HisR4 plasmid, Dr. Florian Krammer from Mount Sinai School of Medicine, for providing the pCAGGS plasmid for SARS-CoV-2 Spike RBD expression, and Dr. Sabra Klein, Johns Hopkins, School of Public Health for generously providing the IgG and IgA ELISA protocols. We also acknowledge Bob Ashley at Penn State College of Medicine for pilot electron microscopy efforts. Lastly, we would also like to acknowledge the Sartorius Huck Cell Culture Facility for early access and production of SARS-CoV-2 Spike RBD protein.

## DISCLOSURES

We wish to disclose the following intellectual property claims related to this study:

**Lindner, S.E**. & **Hafenstein, S**. US Patent Application 16/494,502 “Versatile Display for Proteins”

**Lindner, S.E**., **Hafenstein, S**., & Butler, N. PCT/US2020/033785 “Specific Selection of Immune Cells Using Versatile Display Scaffolds”

**Lindner, S.E**., **Sutton, T.C**., **Hafenstein, S**., & Butler, N. US Trademark Application 9063755 “SPYCAGE”

