## Supplemental Figures, Table, and Files for "Intranasal SARS-CoV-2 RBD decorated nanoparticle vaccine enhances viral clearance in the Syrian hamster model"

#### SUPPLEMENTAL FIGURES, TABLES, AND FILES

**Supplemental Figure 1. Evaluation of the immunogenicity of RBD+SpyCage determined by serum IgG and neutralizing antibody titers.** Shown are the antibody titers from Trial 1 that evaluated immunogenicity of the RBD+SpyCage vaccine in hamsters. Plotted are (A) IgG antibody titers in serum samples collected on days 0, 26, and 55, and (B) neutralizing antibody titers against SARS-CoV-2 on day 0 and 55. * significantly different from all other groups, ($p<0.03$) determined by Kruskal-Wallis test with Dunn's multiple comparison correction.

**Supplemental Figure 2. Weight loss and viral titers in the nasal turbinates and lungs after SARS-CoV-2 challenge of vaccinated hamsters.** After viral challenge, hamsters in Trial 1 (immunogenicity study) hamsters were monitored for (A) weight loss, and viral titers were evaluated in (B) lung tissues and (C) nasal turbinates on days 3 and 6 post-infection.

**Supplemental Figure 3. Weight loss in unvaccinated hamsters after challenge with $10^5$, $10^4$, and $10^3$ TCID50 of SARS-CoV-2.** To determine if lower challenge doses of SARS-CoV-2 would induce weight loss, equal numbers of male and female hamsters (n=3/sex, n=6/group) were intranasally inoculated with $10^5$, $10^4$, and $10^3$ TCID50 of SARS-CoV-2/USA/WA1/2020. Weight loss was then monitored for 14 days until the animals recovered.

**Supplemental Figure 4. Lung histopathology scoring for multiple parameters in vaccinated hamsters challenged with SARS-CoV-2.** On days 3, 5, and 7 post-infection, lung tissues were processed for H&E staining and scored by a veterinary pathologist. Panels display scoring for (A) Type II Pneumocyte Hyperplasia, (B) Alveoli Pathology, (C) Hemorrhage, (D) Blood Vessels Pathology, and (E) Bronchi Pathology. Black horizontal lines indicate median score with the distribution of scores displayed as violin plots.

**Supplementary Table 1: Cryo-EM data collection, processing, and model refinement statistics.**

**Supplemental File 1. Sequences of plasmids used in this study.**

**Supplemental File 2. A validation report for the cryo-EM reconstruction of the apo scaffold based on I3-01 is provided for peer review purposes.**

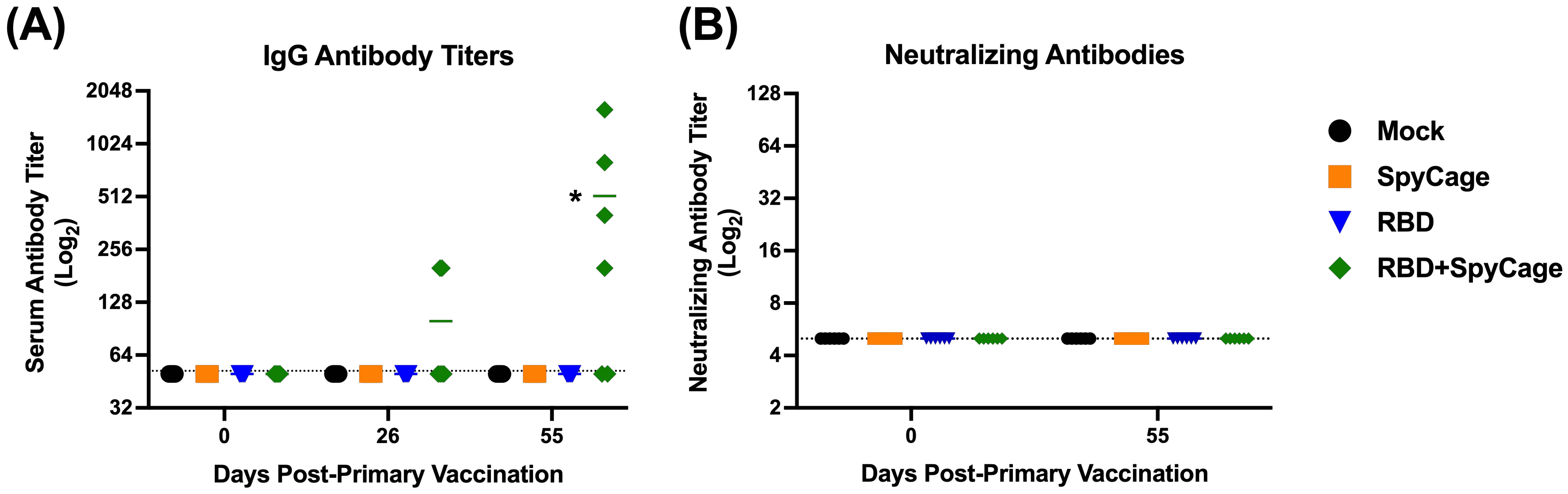

(A)
IgG Antibody Titers
Serum Antibody Titer (Log2)
2048
1024
512
256
128
64
32
*
0
26
55
Days Post-Primary Vaccination
(B)
Neutralizing Antibodies
Neutralizing Antibody Titer (Log2)
128
64
32
16
8
4
2
0
55
Days Post-Primary Vaccination
Mock
SpyCage
RBD
RBD+SpyCage

Figure S1

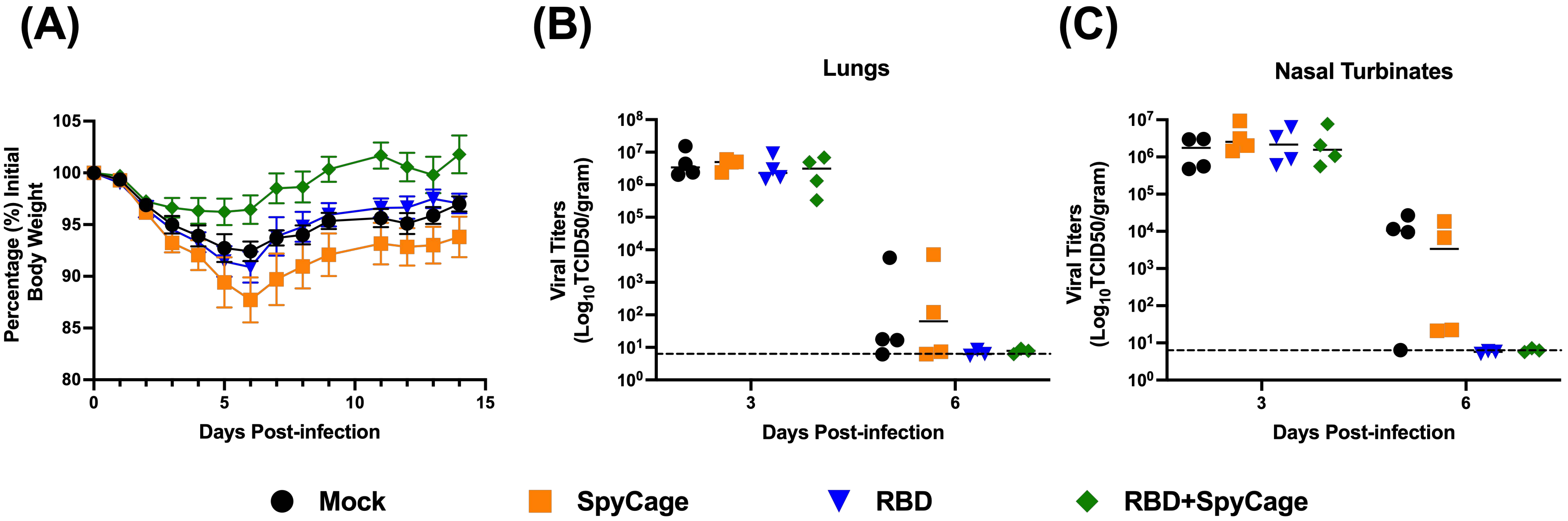
(A)
(B)
(C)
Lungs
Nasal Turbinates
Percentage (%) Initial Body Weight
105
100
95
90
85
80
0
5
10
15
Days Post-infection
Viral Titers ($Log_{10}$TCID50/gram)
$10^8$
$10^7$
$10^6$
$10^5$
$10^4$
$10^3$
$10^2$
$10^1$
$10^0$
3
6
Days Post-infection
Viral Titers ($Log_{10}$TCID50/gram)
$10^7$
$10^6$
$10^5$
$10^4$
$10^3$
$10^2$
$10^1$
$10^0$
3
6
Days Post-infection
Mock
SpyCage
RBD
RBD+SpyCage

Figure S2

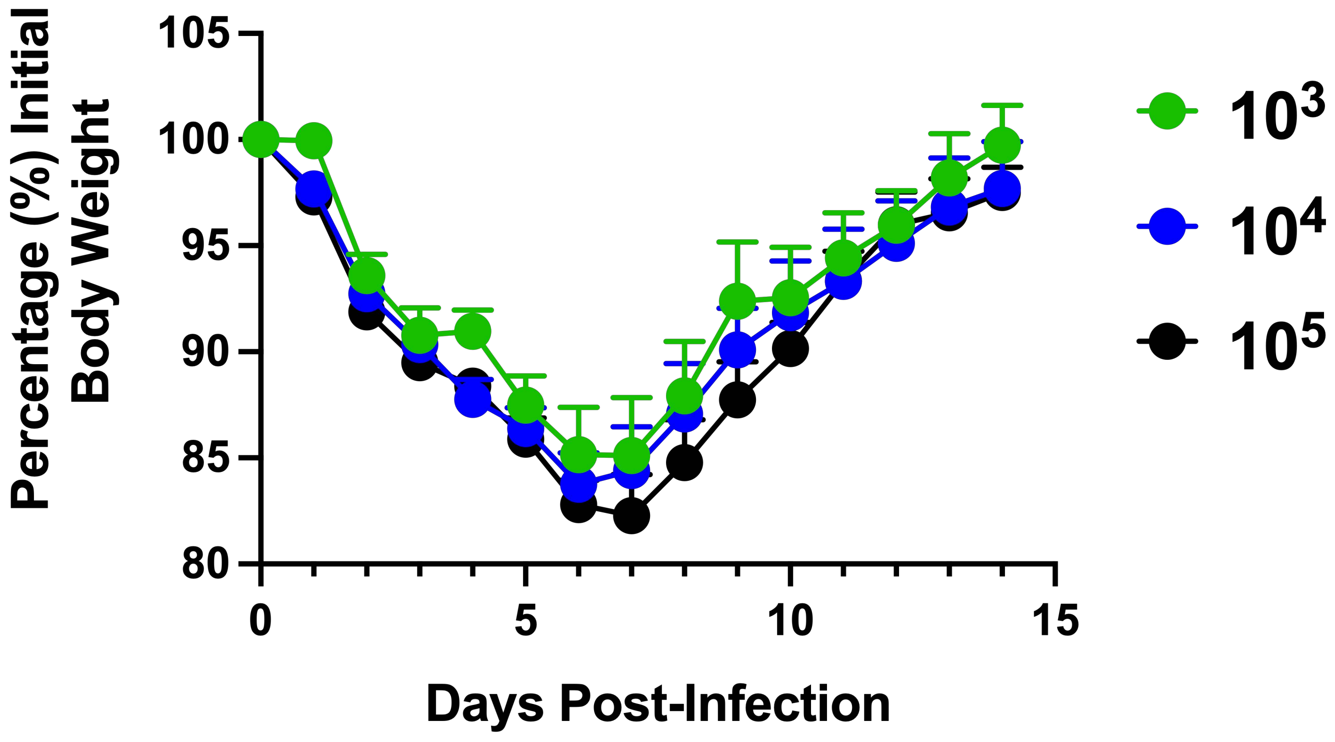
Percentage (%) Initial Body Weight
105
100
95
90
85
80
0
5
10
15
Days Post-Infection
10^3
10^4
10^5

Figure S3

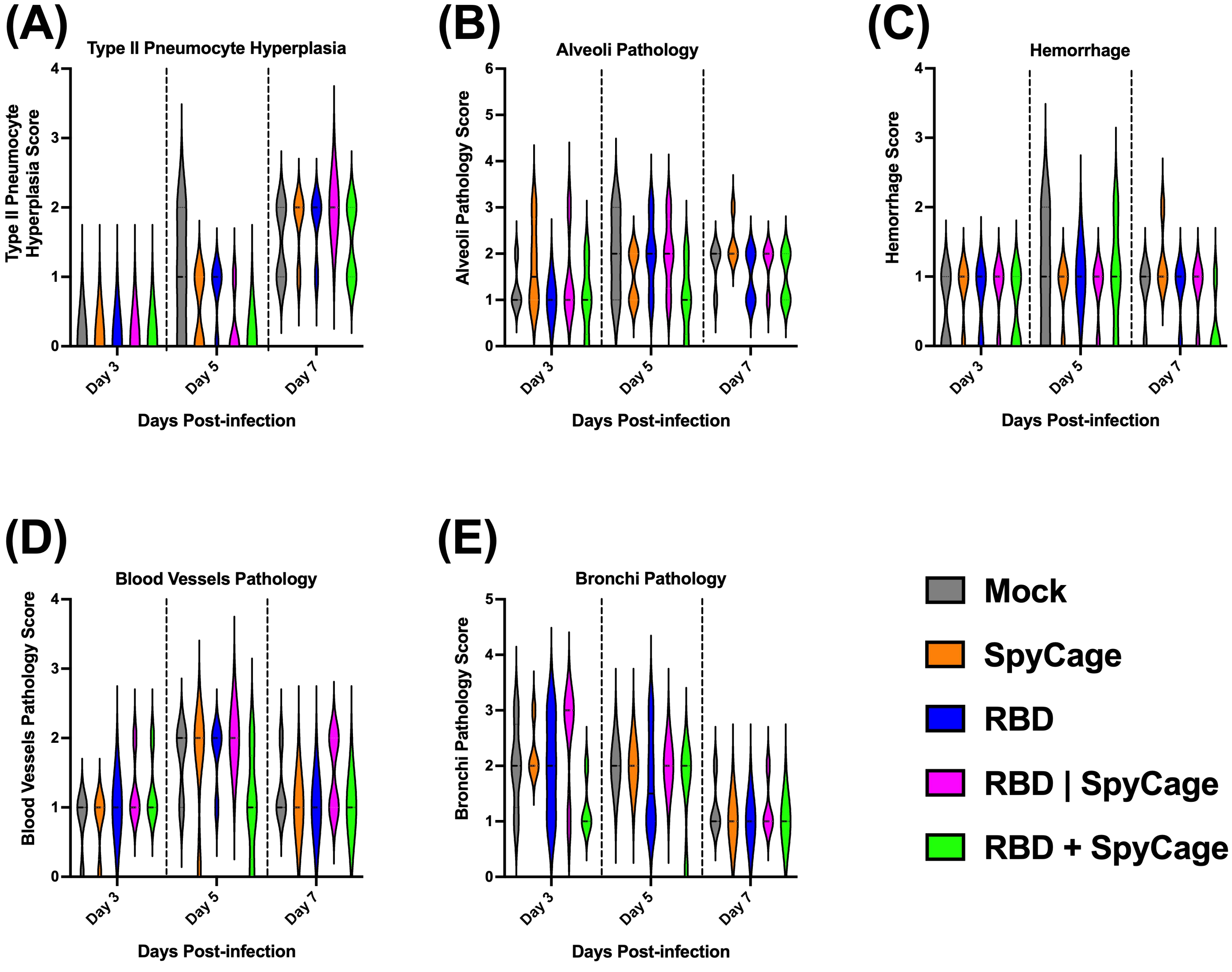
(A)
Type II Pneumocyte Hyperplasia
Type II Pneumocyte Hyperplasia Score
0
1
2
3
4
Day 3
Day 5
Day 7
Days Post-infection
(B)
Alveoli Pathology
Alveoli Pathology Score
0
1
2
3
4
5
6
Day 3
Day 5
Day 7
Days Post-infection
(C)
Hemorrhage
Hemorrhage Score
0
1
2
3
4
Day 3
Day 5
Day 7
Days Post-infection
(D)
Blood Vessels Pathology
Blood Vessels Pathology Score
0
1
2
3
4
Day 3
Day 5
Day 7
Days Post-infection
(E)
Bronchi Pathology
Bronchi Pathology Score
0
1
2
3
4
5
Day 3
Day 5
Day 7
Days Post-infection
Mock
SpyCage
RBD
RBD | SpyCage
RBD + SpyCage

Figure S4

**Supplementary Table 1: Cryo-EM data collection, processing, and model refinement statistics**

| | |
|---|---|
| **Data collection and processing** | |
| Microscope | Titan Krios |
| Detector | Falcon 3EC |
| Voltage (kV) | 300 |
| Recording mode | Counting |
| Nominal magnification | x59,000 |
| Electron exposure ($e^-/Å^2$) | 45 |
| No. of frames | 39 |
| Defocus range (µm) | -1.2 to -3.0 |
| Pixel size (Å) | 1.11 |
| Symmetry | Icosahedral |
| Initial particles (no.) | 129,792 |
| Final particles (no.) | 63,430 |
| Map resolution (Å) | 3.4 |
| FSC Threshold | 0.143 |
| Map sharpening B-factor ($Å^2$) | -196.1 |
| **Model refinement** | |
| Initial model used | I3-01 |
| FSC model vs. map at FSC=0.5 | 3.77 |
| CC model vs. map (masked) | 0.77 |
| Average B-factor ($Å^2$) | 69.1 |
| Bond length rmsd (Å) | 0.002 |
| Bond angle rmsd ($^o$) | 0.562 |
| **Model composition** | |
| No. of chains | 60 |
| Non-hydrogen atoms/chain | 1,519 |
| protein residues/chain | 201 (aa 22-222) |
| **Validation** | |
| Molprobity score | 1.5 |
| Clashscore | 9.4 |
| Poor rotamers (%) | none |
| **Ramachandran plot** | |
| Favored (%) | 97.99 |
| Allowed (%) | 2.01 |
| Disallowed (%) | none |

**Supplemental File 1: Sequences of plasmids used in this study.**

```
>pSL1013_Apo_Cage_Expression_Plasmid
LOCUS       pSL1013               5916 bp    DNA       circular     20-JUL-2022
DEFINITION  .
ACCESSION
VERSION
SOURCE      .
  ORGANISM  .
COMMENT     pET29b(+) from 1 to 5370
COMMENT     pET29b(+)
COMMENT     ApEinfo:methylated:0
FEATURES             Location/Qualifiers
     misc_feature    5076..5750
                     /locus_tag="6xHis_4xGGS_I3-01"
                     /label="6xHis_4xGGS_I3-01"
                     /ApEinfo_label="6xHis_4xGGS_I3-01"
                     /ApEinfo_fwdcolor="cyan"
                     /ApEinfo_revcolor="green"
                     /ApEinfo_graphicformat="arrow_data {{0 1 2 0 0 -1} {} 0}
                     width 5 offset 0"
     rep_origin      1421..2103
                     /locus_tag="ColE1 origin"
                     /label="ColE1 origin"
                     /ApEinfo_label="ColE1 origin"
                     /ApEinfo_fwdcolor="gray50"
                     /ApEinfo_revcolor="gray50"
                     /ApEinfo_graphicformat="arrow_data {{0 1 2 0 0 -1} {} 0}
                     width 5 offset 0"
     rep_origin      29..335
                     /locus_tag="F1 ori"
                     /label="F1 ori"
                     /ApEinfo_label="F1 ori"
                     /ApEinfo_fwdcolor="gray50"
                     /ApEinfo_revcolor="gray50"
                     /ApEinfo_graphicformat="arrow_data {{0 1 2 0 0 -1} {} 0}
                     width 5 offset 0"
     rep_origin      12..467
                     /locus_tag="M13 origin"
                     /label="M13 origin"
                     /ApEinfo_label="M13 origin"
                     /ApEinfo_fwdcolor="gray50"
                     /ApEinfo_revcolor="gray50"
                     /ApEinfo_graphicformat="arrow_data {{0 1 2 0 0 -1} {} 0}
                     width 5 offset 0"
     misc_binding    5007..5029
                     /locus_tag="LacO"
                     /label="LacO"
                     /ApEinfo_label="LacO"
                     /ApEinfo_fwdcolor="#6495ed"
                     /ApEinfo_revcolor="#6495ed"
                     /ApEinfo_graphicformat="arrow_data {{0 1 2 0 0 -1} {} 0}
                     width 5 offset 0"
     CDS             complement(3518..4474)
                     /locus_tag="LacI"
                     /label="LacI"
                     /ApEinfo_label="LacI"
                     /ApEinfo_fwdcolor="gray50"
                     /ApEinfo_revcolor="gray50"
```

/ApEinfo_graphicformat="arrow_data {{0 1 2 0 0 -1} {} 0}
width 5 offset 0"
CDS complement(563..1375)
/locus_tag="KanR"
/label="KanR"
/ApEinfo_label="KanR"
/ApEinfo_fwdcolor="yellow"
/ApEinfo_revcolor="yellow"
/ApEinfo_graphicformat="arrow_data {{0 1 2 0 0 -1} {} 0}
width 5 offset 0"
misc_feature 5788..5916
/locus_tag="T7 Terminator"
/label="T7 Terminator"
/ApEinfo_label="T7 Terminator"
/ApEinfo_fwdcolor="#8080ff"
/ApEinfo_revcolor="#8080ff"
/ApEinfo_graphicformat="arrow_data {{0 1 2 0 0 -1} {} 0}
width 5 offset 0"
misc_feature 4988..5006
/locus_tag="T7prom"
/label="T7prom"
/ApEinfo_label="T7prom"
/ApEinfo_fwdcolor="#8080ff"
/ApEinfo_revcolor="#8080ff"
/ApEinfo_graphicformat="arrow_data {{0 1 2 0 0 -1} {} 0}
width 5 offset 0"
misc_feature 5760..5777
/locus_tag="6xHis Epitope Tag (not translated)"
/label="6xHis Epitope Tag (not translated)"
/ApEinfo_label="6xHis Epitope Tag (not translated)"
/ApEinfo_fwdcolor="#0080c0"
/ApEinfo_revcolor="green"
/ApEinfo_graphicformat="arrow_data {{0 1 2 0 0 -1} {} 0}
width 5 offset 0"
ORIGIN
1 tggcgaatgg gacgcgccct gtagcggcgc attaagcgcg gcgggtgtgg tggttacgcg
61 cagcgtgacc gctacacttg ccagcgccct agcgcccgct cctttcgctt tcttcccttc
121 ctttctcgcc acgttcgccg gctttccccg tcaagctcta aatcgggggc tccctttagg
181 gttccgattt agtgctttac ggcacctcga ccccaaaaaa cttgattagg gtgatggttc
241 acgtagtggg ccatcgccct gatagacggt ttttcgccct ttgacgttgg agtccacgtt
301 ctttaatagt ggactcttgt tccaaactgg aacaacactc aaccctatct cggtctattc
361 ttttgattta taagggattt tgccgatttc ggcctattgg ttaaaaaatg agctgattta
421 acaaaaattt aacgcgaatt ttaacaaaat attaacgttt acaatttcag gtggcacttt
481 tcggggaaat gtgcgcggaa cccctatttg tttatttttc taaatacatt caaatatgta
541 tccgctcatg aattaattct tagaaaaact catcgagcat caaatgaaac tgcaatttat
601 tcatatcagg attatcaata ccatattttt gaaaaagccg tttctgtaat gaaggagaaa
661 actcaccgag gcagttccat aggatggcaa gatcctggta tcggtctgcg attccgactc
721 gtccaacatc aatacaacct attaatttcc cctcgtcaaa aataaggtta tcaagtgaga
781 aatcaccatg agtgacgact gaatccggtg agaatggcaa aagtttatgc atttctttcc
841 agacttgttc aacaggccag ccattacgct cgtcatcaaa atcactcgca tcaaccaaac
901 cgttattcat tcgtgattgc gcctgagcga gacgaaatac gcgatcgctg ttaaaaggac
961 aattacaaac aggaatcgaa tgcaaccggc gcaggaacac tgccagcgca tcaacaatat
1021 tttcacctga atcaggatat tcttctaata cctggaatgc tgttttcccg gggatcgcag
1081 tggtgagtaa ccatgcatca tcaggagtac ggataaaatg cttgatggtc ggaagaggca
1141 taaattccgt cagccagttt agtctgacca tctcatctgt aacatcattg gcaacgctac
1201 ctttgccatg tttcagaaac aactctggcg catcgggctt cccatacaat cgatagattg
1261 tcgcacctga ttgcccgaca ttatcgcgag cccatttata cccatataaa tcagcatcca
1321 tgttggaatt taatcgcggc ctagagcaag acgtttcccg ttgaatatgg ctcataacac

1381 cccttgtatt actgtttatg taagcagaca gttttattgt tcatgaccaa aatcccttaa
1441 cgtgagtttt cgttccactg agcgtcagac cccgtagaaa agatcaaagg atcttcttga
1501 gatccttttt ttctgcgcgt aatctgctgc ttgcaaacaa aaaaaccacc gctaccagcg
1561 gtggtttgtt tgccggatca agagctacca actctttttc cgaaggtaac tggcttcagc
1621 agagcgcaga taccaaatac tgtccttcta gtgtagccgt agttaggcca ccacttcaag
1681 aactctgtag caccgcctac atacctcgct ctgctaatcc tgttaccagt ggctgctgcc
1741 agtggcgata agtcgtgtct taccgggttg gactcaagac gatagttacc ggataaggcg
1801 cagcggtcgg gctgaacggg gggttcgtgc acacagccca gcttggagcg aacgacctac
1861 accgaactga gatacctaca gcgtgagcta tgagaaagcg ccacgcttcc cgaagggaga
1921 aaggcggaca ggtatccggt aagcggcagg gtcggaacag gagagcgcac gagggagctt
1981 ccagggggaa acgcctggta tctttatagt cctgtcgggt ttcgccacct ctgacttgag
2041 cgtcgatttt tgtgatgctc gtcagggggg cggagcctat ggaaaaacgc cagcaacgcg
2101 gcctttttac ggttcctggc cttttgctgg ccttttgctc acatgttctt tcctgcgtta
2161 tcccctgatt ctgtggataa ccgtattacc gcctttgagt gagctgatac cgctcgccgc
2221 agccgaacga ccgagcgcag cgagtcagtg agcgaggaag cggaagagcg cctgatgcgg
2281 tattttctcc ttacgcatct gtgcggtatt tcacaccgca tatatggtgc actctcagta
2341 caatctgctc tgatgccgca tagttaagcc agtatacact ccgctatcgc tacgtgactg
2401 ggtcatggct gcgccccgac acccgccaac acccgctgac gcgccctgac gggcttgtct
2461 gctcccggca tccgcttaca gacaagctgt gaccgtctcc gggagctgca tgtgtcagag
2521 gttttcaccg tcatcaccga aacgcgcgag gcagctgcgg taaagctcat cagcgtggtc
2581 gtgaagcgat tcacagatgt ctgcctgttc atccgcgtcc agctcgttga gtttctccag
2641 aagcgttaat gtctggcttc tgataaagcg ggccatgtta agggcggttt tttcctgttt
2701 ggtcactgat gcctccgtgt aagggggatt tctgttcatg gggtaatga taccgatgaa
2761 acgagagagg atgctcacga tacgggttac tgatgatgaa catgcccggt tactggaacg
2821 ttgtgagggt aaacaactgg cggtatggat gcggcgggac cagagaaaaa tcactcaggg
2881 tcaatgccag cgcttcgtta atacagatgt aggtgttcca cagggtagcc agcagcatcc
2941 tgcgatgcag atccggaaca taatggtgca gggcgctgac ttccgcgttt ccagacttta
3001 cgaaacacgg aaaccgaaga ccattcatgt tgttgctcag gtcgcagacg ttttgcagca
3061 gcagtcgctt cacgttcgct cgcgtatcgg tgattcattc tgctaaccag taaggcaacc
3121 ccgccagcct agccgggtcc tcaacgacag gagcacgatc atgcgcaccc gtggggccgc
3181 catgccggcg ataatggcct gcttctcgcc gaaacgtttg gtggcgggac cagtgacgaa
3241 ggcttgagcg agggcgtgca agattccgaa taccgcaagc gacaggccga tcatcgtcgc
3301 gctccagcga aagcggtcct cgccgaaaat gacccagagc gctgccggca cctgtcctac
3361 gagttgcatg ataaagaaga cagtcataag tgcggcgacg atagtcatgc cccgcgccca
3421 ccggaaggag ctgactgggt tgaaggctct caagggcatc ggtcgagatc ccggtgccta
3481 atgagtgagc taacttacat taattgcgtt gcgctcactg cccgctttcc agtcgggaaa
3541 cctgtcgtgc cagctgcatt aatgaatcgg ccaacgcgcg gggagaggcg gtttgcgtat
3601 tgggcgccag ggtggttttt cttttcacca gtgagacggg caacagctga ttgcccttca
3661 ccgcctggcc ctgagagagt tgcagcaagc ggtccacgct ggtttgcccc agcaggcgaa
3721 aatcctgttt gatggtggtt aacggcggga tataacatga gctgtcttcg gtatcgtcgt
3781 atcccactac cgagatgtcc gcaccaacgc gcagcccgga ctcggtaatg gcgcgcattg
3841 cgcccagcgc catctgatcg ttggcaacca gcatcgcagt gggaacgatg ccctcattca
3901 gcatttgcat ggtttgttga aaaccggaca tggcactcca gtcgccttcc cgttccgcta
3961 tcggctgaat ttgattgcga gtgagatatt tatgccagcc agccagacgc agacgcgccg
4021 agacagaact taatgggccc gctaacagcg cgatttgctg gtgacccaat gcgaccagat
4081 gctccacgcc cagtcgcgta ccgtcttcat gggagaaaat aatactgttg atgggtgtct
4141 ggtcagagac atcaagaaat aacgccggaa cattagtgca ggcagcttcc acagcaatgg
4201 catcctggtc atccagcgga tagttaatga tcagcccact gacgcgttgc gcgagaagat
4261 tgtgcaccgc cgctttacag gcttcgacgc cgcttcgttc taccatcgac accaccacgc
4321 tggcacccag ttgatcggcg cgagatttaa tcgccgcgac aatttgcgac ggcgcgtgca
4381 gggccagact ggaggtggca acgccaatca gcaacgactg tttgcccgcc agttgttgtg
4441 ccacgcggtt gggaatgtaa ttcagctccg ccatcgccgc ttccactttt tcccgcgttt
4501 tcgcagaaac gtggctggcc tggttcacca cgcgggaaac ggtctgataa gagacaccgg
4561 catactctgc gacatcgtat aacgttactg gtttcacatt caccaccctg aattgactct
4621 cttccgggcg ctatcatgcc ataccgcgaa aggttttgcg ccattcgatg gtgtccggga
4681 tctcgacgct ctcccttatg cgactcctgc attaggaagc agcccagtag taggttgagg
4741 ccgttgagca ccgccgccgc aaggaatggt gcatgcaagg agatggcgcc caacagtccc
4801 ccggccacgg ggcctgccac catacccacg ccgaaacaag cgctcatgag cccgaagtgg

4861 cgagcccgat cttccccatc ggtgatgtcg gcgatatagg cgccagcaac cgcacctgtg
4921 gcgccggtga tgccggccac gatgcgtccg gcgtagagga tcgagatcga tctcgatccc
4981 gcgaaattaa tacgactcac tataggggaa ttgtgagcgg ataacaattc ccctctagaa
5041 ataattttgt ttaactttaa gaaggagata taCATATGCA CCACCACCAC CACCACGGCG
5101 GCAGCGGCGG CAGCGGCGGT AGCGGCGGTA GCATGAAGAT GGAAGAGCTG TTCAAGAAAC
5161 ACAAGATCGT TGCCGTGCTG CGTGCCAATA GTGTGGAAGA AGCGAAAAAG AAAGCGCTGG
5221 CGGTTTTCCT GGGCGGCGTT CATCTGATTG AAATTACCTT TACCGTGCCG GATGCGGATA
5281 CCGTGATTAA GGAACTGAGC TTTCTGAAGG AAATGGGCGC GATTATTGGT GCGGGCACCG
5341 TGACCAGCGT GGAGCAGTGC CGTAAAGCGG TGGAAAGTGG CGCCGAATTC ATTGTGAGTC
5401 CGCACCTGGA CGAGGAAATT AGCCAATTTT GCAAGGAGAA GGGTGTGTTC TATATGCCAG
5461 GCGTTATGAC CCCGACCGAA CTGGTGAAAG CCATGAAACT GGGCCATACC ATCTTAAAAC
5521 TGTTTCCGGG TGAGGTGGTG GGTCCGCAGT TTGTTAAAGC GATGAAAGGT CCGTTTCCGA
5581 ATGTGAAATT TGTGCCAACC GGCGGTGTTA ATCTGGACAA TGTGTGCGAA TGGTTCAAAG
5641 CGGGCGTGCT GGCCGTGGGC GTGGGCAGCG CGTTAGTGAA AGGCACCCCG GTGGAAGTGG
5701 CGGAAAAGGC CAAGGCGTTC GTTGAGAAGA TTCGTGGCTG CACCGAATAA TAGCTCGAGc
5761 accaccacca ccaccactga gatccggctg ctaacaaagc ccgaaaggaa gctgagttgg
5821 ctgctgccac cgctgagcaa taactagcat aaccccttgg ggcctctaaa cgggtcttga
5881 ggggtttttt gctgaaagga ggaactatat ccggat
//

```
>pSL1040_SpyCage-Flexible_Expression_Plasmid
LOCUS       pSL1040                6259 bp    DNA       circular     12-MAY-2022
DEFINITION  .
ACCESSION
VERSION
SOURCE      .
  ORGANISM  .
COMMENT     pSL0045+I3-01
COMMENT     ApEinfo:methylated:0
FEATURES             Location/Qualifiers
     CDS             5071..5388
                     /codon_start=1
                     /note="SpyCatcher"
                     /translation="MGSSHHHHHHGSGDSATHIKFSKRDEDGKELAGATMELRDSSGK
                     T
                     ISTWISDGQVKDFYLYPGKYTFVETAAPDGYEVATAITFTVNEQGQVTVNGKATKGDA
                     H
                     IGSGGSGGSGANKPMQPITSTANKIVWSDPTRLSTTFSASLLRQRVKVGIAELNNVSG
                     Q
                     YVSVYKRPAPKPEGCADACVIMPNENQSIRTVISGSAENLATLKAEWETHKRNVDTLF
                     A SGNAGLGFLDPTAAIVSSDTTA"
                     /locus_tag="SpyCatcher"
                     /label="SpyCatcher"
                     /ApEinfo_label="SpyCatcher"
                     /ApEinfo_fwdcolor="pink"
                     /ApEinfo_revcolor="pink"
                     /ApEinfo_graphicformat="arrow_data {{0 1 2 0 0 -1} {} 0}
                     width 5 offset 0"
     primer_bind     4984..5003
                     /locus_tag="T7"
                     /label="T7"
                     /ApEinfo_label="T7"
                     /ApEinfo_fwdcolor="cyan"
                     /ApEinfo_revcolor="green"
                     /ApEinfo_graphicformat="arrow_data {{0 1 2 0 0 -1} {} 0}
                     width 5 offset 0"
     CDS             5071..5073
                     /codon_start=1
                     /note="TIR 10295"
                     /translation="M"
                     /locus_tag="TIR 10295"
                     /label="TIR 10295"
                     /ApEinfo_label="TIR 10295"
                     /ApEinfo_fwdcolor="pink"
                     /ApEinfo_revcolor="pink"
                     /ApEinfo_graphicformat="arrow_data {{0 1 2 0 0 -1} {} 0}
                     width 5 offset 0"
     rep_origin      1421..2103
                     /locus_tag="Origin of DNA Replication"
                     /label="Origin of DNA Replication"
                     /ApEinfo_label="Origin of DNA Replication"
                     /ApEinfo_fwdcolor="gray50"
                     /ApEinfo_revcolor="gray50"
                     /ApEinfo_graphicformat="arrow_data {{0 1 2 0 0 -1} {} 0}
                     width 5 offset 0"
     rep_origin      12..467
                     /locus_tag="M13 origin"
                     /label="M13 origin"
```

```
                    /ApEinfo_label="M13 origin"
                    /ApEinfo_fwdcolor="gray50"
                    /ApEinfo_revcolor="gray50"
                    /ApEinfo_graphicformat="arrow_data {{0 1 2 0 0 -1} {} 0}
                    width 5 offset 0"
misc_feature    4984..5002
                    /locus_tag="T7prom"
                    /label="T7prom"
                    /ApEinfo_label="T7prom"
                    /ApEinfo_fwdcolor="#8080ff"
                    /ApEinfo_revcolor="#8080ff"
                    /ApEinfo_graphicformat="arrow_data {{0 1 2 0 0 -1} {} 0}
                    width 5 offset 0"
CDS             5083..5100
                    /codon_start=1
                    /product="6xHis"
                    /note="6xHis"
                    /translation="HHHHHH"
                    /locus_tag="6xHis"
                    /label="6xHis"
                    /ApEinfo_label="6xHis"
                    /ApEinfo_fwdcolor="pink"
                    /ApEinfo_revcolor="pink"
                    /ApEinfo_graphicformat="arrow_data {{0 1 2 0 0 -1} {} 0}
                    width 5 offset 0"
rep_origin      29..335
                    /locus_tag="F1 ori"
                    /label="F1 ori"
                    /ApEinfo_label="F1 ori"
                    /ApEinfo_fwdcolor="gray50"
                    /ApEinfo_revcolor="gray50"
                    /ApEinfo_graphicformat="arrow_data {{0 1 2 0 0 -1} {} 0}
                    width 5 offset 0"
CDS             5386..5388
                    /codon_start=1
                    /note="3xGSG"
                    /translation="GSGGSGGSG"
                    /locus_tag="3xGSG"
                    /label="3xGSG"
                    /ApEinfo_label="3xGSG"
                    /ApEinfo_fwdcolor="pink"
                    /ApEinfo_revcolor="pink"
                    /ApEinfo_graphicformat="arrow_data {{0 1 2 0 0 -1} {} 0}
                    width 5 offset 0"
CDS             complement(563..1375)
                    /locus_tag="KanR"
                    /label="KanR"
                    /ApEinfo_label="KanR"
                    /ApEinfo_fwdcolor="yellow"
                    /ApEinfo_revcolor="yellow"
                    /ApEinfo_graphicformat="arrow_data {{0 1 2 0 0 -1} {} 0}
                    width 5 offset 0"
misc_feature    6079..6096
                    /locus_tag="TEVc Site"
                    /label="TEVc Site"
                    /ApEinfo_label="TEVc Site"
                    /ApEinfo_fwdcolor="#ff0000"
                    /ApEinfo_revcolor="green"
```

```
                     /ApEinfo_graphicformat="arrow_data {{0 1 2 0 0 -1} {} 0}
                     width 5 offset 0"
     misc_feature    5137..5139
                     /note="reactive 255K from CnaB2"
                     /locus_tag="reactive 255K from CnaB2"
                     /label="reactive 255K from CnaB2"
                     /ApEinfo_label="reactive 255K from CnaB2"
                     /ApEinfo_fwdcolor="#0080ff"
                     /ApEinfo_revcolor="#008200"
                     /ApEinfo_graphicformat="arrow_data {{0 1 2 0 0 -1} {} 0}
                     width 5 offset 0"
     misc_binding    5003..5025
                     /locus_tag="LacO"
                     /label="LacO"
                     /ApEinfo_label="LacO"
                     /ApEinfo_fwdcolor="#6495ed"
                     /ApEinfo_revcolor="#6495ed"
                     /ApEinfo_graphicformat="arrow_data {{0 1 2 0 0 -1} {} 0}
                     width 5 offset 0"
     misc_feature    complement(3515..4606)
                     /locus_tag="lacI"
                     /label="lacI"
                     /ApEinfo_label="lacI"
                     /ApEinfo_fwdcolor="#8080c0"
                     /ApEinfo_revcolor="#008200"
                     /ApEinfo_graphicformat="arrow_data {{0 1 2 0 0 -1} {} 0}
                     width 5 offset 0"
     misc_feature    5395..6063
                     /locus_tag="I3-01"
                     /label="I3-01"
                     /ApEinfo_label="I3-01"
                     /ApEinfo_fwdcolor="cyan"
                     /ApEinfo_revcolor="green"
                     /ApEinfo_graphicformat="arrow_data {{0 1 2 0 0 -1} {} 0}
                     width 5 offset 0"
     misc_feature    6131..6259
                     /locus_tag="T7 Terminator"
                     /label="T7 Terminator"
                     /ApEinfo_label="T7 Terminator"
                     /ApEinfo_fwdcolor="#0000a0"
                     /ApEinfo_revcolor="#0000a0"
                     /ApEinfo_graphicformat="arrow_data {{0 1 2 0 0 -1} {} 0}
                     width 5 offset 0"
     misc_feature    6103..6120
                     /locus_tag="6xHis Epitope Tag"
                     /label="6xHis Epitope Tag"
                     /ApEinfo_label="6xHis Epitope Tag"
                     /ApEinfo_fwdcolor="#0080c0"
                     /ApEinfo_revcolor="green"
                     /ApEinfo_graphicformat="arrow_data {{0 1 2 0 0 -1} {} 0}
                     width 5 offset 0"
ORIGIN
        1 tggcgaatgg gacgcgccct gtagcggcgc attaagcgcg gcgggtgtgg tggttacgcg
       61 cagcgtgacc gctacacttg ccagcgccct agcgcccgct cctttcgctt tcttcccttc
      121 ctttctcgcc acgttcgccg gctttccccg tcaagctcta aatcgggggc tccctttagg
      181 gttccgattt agtgctttac ggcacctcga ccccaaaaaa cttgattagg gtgatggttc
      241 acgtagtggg ccatcgccct gatagacggt ttttcgccct ttgacgttgg agtccacgtt
      301 ctttaatagt ggactcttgt tccaaactgg aacaacactc aaccctatct cggtctattc
```

361 ttttgattta taagggattt tgccgatttc ggcctattgg ttaaaaaatg agctgattta
421 acaaaaattt aacgcgaatt ttaacaaaat attaacgttt acaatttcag gtggcacttt
481 tcggggaaat gtgcgcggaa cccctatttg tttatttttc taaatacatt caaatatgta
541 tccgctcatg aattaattct tagaaaaact catcgagcat caaatgaaac tgcaatttat
601 tcatatcagg attatcaata ccatattttt gaaaaagccg tttctgtaat gaaggagaaa
661 actcaccgag gcagttccat aggatggcaa gatcctggta tcggtctgcg attccgactc
721 gtccaacatc aatacaacct attaatttcc cctcgtcaaa aataaggtta tcaagtgaga
781 aatcaccatg agtgacgact gaatccggtg agaatggcaa aagtttatgc atttctttcc
841 agacttgttc aacaggccag ccattacgct cgtcatcaaa atcactcgca tcaaccaaac
901 cgttattcat tcgtgattgc gcctgagcga gacgaaatac gcgatcgctg ttaaaaggac
961 aattacaaac aggaatcgaa tgcaaccggc gcaggaacac tgccagcgca tcaacaatat
1021 tttcacctga atcaggatat tcttctaata cctggaatgc tgttttcccg gggatcgcag
1081 tggtgagtaa ccatgcatca tcaggagtac ggataaaatg cttgatggtc ggaagaggca
1141 taaattccgt cagccagttt agtctgacca tctcatctgt aacatcattg gcaacgctac
1201 ctttgccatg tttcagaaac aactctggcg catcgggctt cccatacaat cgatagattg
1261 tcgcacctga ttgcccgaca ttatcgcgag cccatttata cccatataaa tcagcatcca
1321 tgttggaatt taatcgcggc ctagagcaag acgtttcccg ttgaatatgg ctcataacac
1381 cccttgtatt actgtttatg taagcagaca gttttattgt tcatgaccaa aatcccttaa
1441 cgtgagtttt cgttccactg agcgtcagac cccgtagaaa agatcaaagg atcttcttga
1501 gatccttttt ttctgcgcgt aatctgctgc ttgcaaacaa aaaaaccacc gctaccagcg
1561 gtggtttgtt tgccggatca agagctacca actctttttc cgaaggtaac tggcttcagc
1621 agagcgcaga taccaaatac tgtccttcta gtgtagccgt agttaggcca ccacttcaag
1681 aactctgtag caccgcctac atacctcgct ctgctaatcc tgttaccagt ggctgctgcc
1741 agtggcgata agtcgtgtct taccgggttg gactcaagac gatagttacc ggataaggcg
1801 cagcggtcgg gctgaacggg gggttcgtgc acacagccca gcttggagcg aacgacctac
1861 accgaactga gatacctaca gcgtgagcta tgagaaagcg ccacgcttcc cgaagggaga
1921 aaggcggaca ggtatccggt aagcggcagg gtcggaacag gagagcgcac gagggagctt
1981 ccagggggaa acgcctggta tctttatagt cctgtcgggt ttcgccacct ctgacttgag
2041 cgtcgatttt tgtgatgctc gtcagggggg cggagcctat ggaaaaacgc cagcaacgcg
2101 gcctttttac ggttcctggc cttttgctgg ccttttgctc acatgttctt tcctgcgtta
2161 tcccctgatt ctgtggataa ccgtattacc gcctttgagt gagctgatac cgctcgccgc
2221 agccgaacga ccgagcgcag cgagtcagtg agcgaggaag cggaagagcg cctgatgcgg
2281 tattttctcc ttacgcatct gtgcggtatt tcacaccgca tatatggtgc actctcagta
2341 caatctgctc tgatgccgca tagttaagcc agtatacact ccgctatcgc tacgtgactg
2401 ggtcatggct gcgccccgac acccgccaac acccgctgac gcgccctgac gggcttgtct
2461 gctcccggca tccgcttaca gacaagctgt gaccgtctcc gggagctgca tgtgtcagag
2521 gttttcaccg tcatcaccga aacgcgcgag gcagctgcgg taaagctcat cagcgtggtc
2581 gtgaagcgat tcacagatgt ctgcctgttc atccgcgtcc agctcgttga gtttctccag
2641 aagcgttaat gtctggcttc tgataaagcg ggccatgtta agggcggttt tttcctgttt
2701 ggtcactgat gcctccgtgt aagggggatt tctgttcatg ggggtaatga taccgatgaa
2761 acgagagagg atgctcacga tacgggttac tgatgatgaa catgcccggt tactggaacg
2821 ttgtgagggt aaacaactgg cggtatggat gcggcgggac cagagaaaaa tcactcaggg
2881 tcaatgccag cgcttcgtta atacagatgt aggtgttcca cagggtagcc agcagcatcc
2941 tgcgatgcag atccggaaca taatggtgca gggcgctgac ttccgcgttt ccagacttta
3001 cgaaacacgg aaaccgaaga ccattcatgt tgttgctcag gtcgcagacg ttttgcagca
3061 gcagtcgctt cacgttcgct cgcgtatcgg tgattcattc tgctaaccag taaggcaacc
3121 ccgccagcct agccgggtcc tcaacgacag gagcacgatc atgcgcaccc gtggggccgc
3181 catgccggcg ataatggcct gcttctcgcc gaaacgtttg gtggcgggac cagtgacgaa
3241 ggcttgagcg agggcgtgca agattccgaa taccgcaagc gacaggccga tcatcgtcgc
3301 gctccagcga aagcggtcct cgccgaaaat gacccagagc gctgccggca cctgtcctac
3361 gagttgcatg ataaagaaga cagtcataag tgcggcgacg atagtcatgc cccgcgccca
3421 ccggaaggag ctgactgggt tgaaggctct caagggcatc ggtcgagatc ccggtgccta
3481 atgagtgagc taacttacat taattgcgtt gcgctcactg cccgctttcc agtcgggaaa
3541 cctgtcgtgc cagctgcatt aatgaatcgg ccaacgcgcg gggagaggcg gtttgcgtat
3601 tgggcgccag ggtggttttt cttttcacca gtgagacggg caacagctga ttgcccttca
3661 ccgcctggcc ctgagagagt tgcagcaagc ggtccacgct ggtttgcccc agcaggcgaa
3721 aatcctgttt gatggtggtt aacggcggga tataacatga gctgtcttcg gtatcgtcgt
3781 atcccactac cgagatatcc gcaccaacgc gcagcccgga ctcggtaatg gcgcgcattg

3841 cgcccagcgc catctgatcg ttggcaacca gcatcgcagt gggaacgatg ccctcattca
3901 gcatttgcat ggtttgttga aaaccggaca tggcactcca gtcgccttcc cgttccgcta
3961 tcggctgaat ttgattgcga gtgagatatt tatgccagcc agccagacgc agacgcgccg
4021 agacagaact taatgggccc gctaacagcg cgatttgctg gtgacccaat gcgaccagat
4081 gctccacgcc cagtcgcgta ccgtcttcat gggagaaaat aatactgttg atgggtgtct
4141 ggtcagagac atcaagaaat aacgccggaa cattagtgca ggcagcttcc acagcaatgg
4201 catcctggtc atccagcgga tagttaatga tcagcccact gacgcgttgc gcgagaagat
4261 tgtgcaccgc cgctttacag gcttcgacgc cgcttcgttc taccatcgac accaccacgc
4321 tggcacccag ttgatcggcg cgagatttaa tcgccgcgac aatttgcgac ggcgcgtgca
4381 gggccagact ggaggtggca acgccaatca gcaacgactg tttgcccgcc agttgttgtg
4441 ccacgcggtt gggaatgtaa ttcagctccg ccatcgccgc ttccactttt tcccgcgttt
4501 tcgcagaaac gtggctggcc tggttcacca cgcgggaaac ggtctgataa gagacaccgg
4561 catactctgc gacatcgtat aacgttactg gtttcacatt caccaccctg aattgactct
4621 cttccgggcg ctatcatgcc ataccgcgaa aggttttgcg ccattcgatg gtgtccggga
4681 tctcgacgct ctcccttatg cgactcctgc attaggaagc agcccagtag taggttgagg
4741 ccgttgagca ccgccgccgc aaggaatggt gcatgcaagg agatggcgcc caacagtccc
4801 ccggccacgg ggcctgccac catacccacg ccgaaacaag cgctcatgag cccgaagtgg
4861 cgagcccgat cttccccatc ggtgatgtcg gcgatatagg cgccagcaac cgcacctgtg
4921 gcgccggtga tgccggccac gatgcgtccg gcgtagagga tcgagatctc gatcccgcga
4981 aattaatacg actcactata ggggaattgt gagcggataa caattcccct ctagaaataa
5041 ttttgtttaa ctttaagaag gagatatacc atggGCAGCA GCCATCATCA TCATCATCAC
5101 GGCAGCGGCG ATAGTGCTAC CCATATTAAA TTCTCAAAAC GTGATGAGGA CGGCAAAGAG
5161 TTAGCTGGTG CAACTATGGA GTTGCGTGAT TCATCTGGTA AAACTATTAG TACATGGATT
5221 TCAGATGGAC AAGTGAAAGA TTTCTACCTG TATCCAGGAA AATATACATT TGTCGAAACC
5281 GCAGCACCAG ACGGTTATGA GGTAGCAACT GCTATTACCT TTACAGTTAA TGAGCAAGGT
5341 CAGGTTACTG TAAACGGCAA AGCAACTAAA GGTGACGCTC ATATTGGCgt cgacCACCAC
5401 CACCACCACC ACGGCGGCAG CGGCGGCAGC GGCGGTAGCG GCGGTAGCAT GAAGATGGAA
5461 GAGCTGTTCA AGAAACACAA GATCGTTGCC GTGCTGCGTG CCAATAGTGT GGAAGAAGCG
5521 AAAAAGAAAG CGCTGGCGGT TTTCCTGGGC GGCGTTCATC TGATTGAAAT TACCTTTACC
5581 GTGCCGGATG CGGATACCGT GATTAAGGAA CTGAGCTTTC TGAAGGAAAT GGGCGCGATT
5641 ATTGGTGCGG GCACCGTGAC CAGCGTGGAG CAGTGCCGTA AAGCGGTGGA AAGTGGCGCC
5701 GAATTCATTG TGAGTCCGCA CCTGGACGAG GAAATTAGCC AATTTTGCAA GGAGAAGGGT
5761 GTGTTCTATA TGCCAGGCGT TATGACCCCG ACCGAACTGG TGAAAGCCAT GAAACTGGGC
5821 CATACCATCT TAAAACTGTT TCCGGGTGAG GTGGTGGGTC CGCAGTTTGT TAAAGCGATG
5881 AAAGGTCCGT TTCCGAATGT GAAATTTGTG CCAACCGGCG GTGTTAATCT GGACAATGTG
5941 TGCGAATGGT TCAAAGCGGG CGTGCTGGCC GTGGGCGTGG GCAGCGCGTT AGTGAAAGGC
6001 ACCCCGGTGG AAGTGGCGGA AAAGGCCAAG GCGTTCGTTG AGAAGATTCG TGGCTGCACC
6061 GAAcatatgt agctcgagAA CCTGTACTTC CAGGGAgtcg agcaccacca ccaccaccac
6121 tgagatccgg ctgctaacaa agcccgaaag gaagctgagt tggctgctgc caccgctgag
6181 caataactag cataacccct tggggcctct aaacgggtct tgaggggttt tttgctgaaa
6241 ggaggaacta tatccggat
//

```
>pSL1510_SARS-CoV-2-Spike-RBD-NoSpyTag_Expression Plasmid
LOCUS       pSL1510       5498 bp    DNA       circular     12-MAY-2022
DEFINITION  .
ACCESSION
VERSION
SOURCE      .
  ORGANISM  .
COMMENT
COMMENT
COMMENT     ApEinfo:methylated:1
FEATURES             Location/Qualifiers
     rep_origin      complement(3730..4412)
                     /locus_tag="ColE1 origin"
                     /label="ColE1 origin"
                     /ApEinfo_label="ColE1 origin"
                     /ApEinfo_fwdcolor="gray50"
                     /ApEinfo_revcolor="gray50"
                     /ApEinfo_graphicformat="arrow_data {{0 1 2 0 0 -1} {} 0}
                     width 5 offset 0"
     misc_binding    complement(3023..3045)
                     /locus_tag="LacO"
                     /label="LacO"
                     /ApEinfo_label="LacO"
                     /ApEinfo_fwdcolor="#6495ed"
                     /ApEinfo_revcolor="#6495ed"
                     /ApEinfo_graphicformat="arrow_data {{0 1 2 0 0 -1} {} 0}
                     width 5 offset 0"
     misc_feature    3375..3505
                     /locus_tag="SV40 early poly-A signal"
                     /label="SV40 early poly-A signal"
                     /ApEinfo_label="SV40 early poly-A signal"
                     /ApEinfo_fwdcolor="#808000"
                     /ApEinfo_revcolor="green"
                     /ApEinfo_graphicformat="arrow_data {{0 1 2 0 0 -1} {} 0}
                     width 5 offset 0"
     misc_feature    complement(4510..5169)
                     /locus_tag="AmpR"
                     /label="AmpR"
                     /ApEinfo_label="AmpR"
                     /ApEinfo_fwdcolor="#ffff00"
                     /ApEinfo_revcolor="#ffff00"
                     /ApEinfo_graphicformat="arrow_data {{0 1 2 0 0 -1} {} 0}
                     width 5 offset 0"
     misc_feature    3185..3364
                     /locus_tag="SV40 Promoter"
                     /label="SV40 Promoter"
                     /ApEinfo_label="SV40 Promoter"
                     /ApEinfo_fwdcolor="#8080c0"
                     /ApEinfo_revcolor="green"
                     /ApEinfo_graphicformat="arrow_data {{0 1 2 0 0 -1} {} 0}
                     width 5 offset 0"
     misc_feature    complement(776..781)
                     /locus_tag="Ribosome Binding Site"
                     /label="Ribosome Binding Site"
                     /ApEinfo_label="Ribosome Binding Site"
                     /ApEinfo_fwdcolor="cyan"
                     /ApEinfo_revcolor="green"
                     /ApEinfo_graphicformat="arrow_data {{0 1 2 0 0 -1} {} 0}
```

width 5 offset 0"
misc_feature    complement(2918..2923)
/locus_tag="Ribosome Binding Site(1)"
/label="Ribosome Binding Site(1)"
/ApEinfo_label="Ribosome Binding Site"
/ApEinfo_fwdcolor="cyan"
/ApEinfo_revcolor="green"
/ApEinfo_graphicformat="arrow_data {{0 1 2 0 0 -1} {} 0}
width 5 offset 0"
misc_feature    complement(3869..3874)
/locus_tag="Ribosome Binding Site(2)"
/label="Ribosome Binding Site(2)"
/ApEinfo_label="Ribosome Binding Site"
/ApEinfo_fwdcolor="cyan"
/ApEinfo_revcolor="green"
/ApEinfo_graphicformat="arrow_data {{0 1 2 0 0 -1} {} 0}
width 5 offset 0"
misc_feature    complement(3023..3040)
/locus_tag="Minimal LacO"
/label="Minimal LacO"
/ApEinfo_label="Minimal LacO"
/ApEinfo_fwdcolor="#ff80c0"
/ApEinfo_revcolor="green"
/ApEinfo_graphicformat="arrow_data {{0 1 2 0 0 -1} {} 0}
width 5 offset 0"
misc_feature    84..371
/locus_tag="CAG Enhancer / CMV immediate early promoter"
/label="CAG Enhancer / CMV immediate early promoter"
/ApEinfo_label="CAG Enhancer / CMV immediate early
promoter"
/ApEinfo_fwdcolor="#8080c0"
/ApEinfo_revcolor="green"
/ApEinfo_graphicformat="arrow_data {{0 1 2 0 0 -1} {} 0}
width 5 offset 0"
misc_feature    2517..2955
/locus_tag="rb glob PA terminator"
/label="rb glob PA terminator"
/ApEinfo_label="rb glob PA terminator"
/ApEinfo_fwdcolor="#800000"
/ApEinfo_revcolor="green"
/ApEinfo_graphicformat="arrow_data {{0 1 2 0 0 -1} {} 0}
width 5 offset 0"
misc_feature    1734..2465
/locus_tag="SARS-CoV-2 Spike SP+RBD+6xHis"
/label="SARS-CoV-2 Spike SP+RBD+6xHis"
/ApEinfo_label="SARS-CoV-2 Spike SP+RBD+6xHis"
/ApEinfo_fwdcolor="cyan"
/ApEinfo_revcolor="green"
/ApEinfo_graphicformat="arrow_data {{0 1 2 0 0 -1} {} 0}
width 5 offset 0"
misc_feature    385..1630
/locus_tag="CAG promoter"
/label="CAG promoter"
/ApEinfo_label="CAG promoter"
/ApEinfo_fwdcolor="#0080c0"
/ApEinfo_revcolor="green"
/ApEinfo_graphicformat="arrow_data {{0 1 2 0 0 -1} {} 0}
width 5 offset 0"

ORIGIN

```
        1 GTCGACATTG ATTATTGACT AGTTATTAAT AGTAATCAAT TACGGGGTCA TTAGTTCATA
       61 GCCCATATAT GGAGTTCCGC GTTACATAAC TTACGGTAAA TGGCCCGCCT GGCTGACCGC
      121 CCAACGACCC CCGCCCATTG ACGTCAATAA TGACGTATGT TCCCATAGTA ACGCCAATAG
      181 GGACTTTCCA TTGACGTCAA TGGGTGGAGT ATTTACGGTA AACTGCCCAC TTGGCAGTAC
      241 ATCAAGTGTA TCATATGCCA AGTACGCCCC CTATTGACGT CAATGACGGT AAATGGCCCG
      301 CCTGGCATTA TGCCCAGTAC ATGACCTTAT GGGACTTTCC TACTTGGCAG TACATCTACG
      361 TATTAGTCAT CGCTATTACC ATGGTCGAGG TGAGCCCCAC GTTCTGCTTC ACTCTCCCCA
      421 TCTCCCCCCC CTCCCCACCC CCAATTTTGT ATTTATTTAT TTTTTAATTA TTTTGTGCAG
      481 CGATGGGGGC GGGGGGGGGG GGGGCGCGCG CCAGGCGGGG CGGGGCGGGG CGAGGGGCGG
      541 GGCGGGGCGA GGCGGAGAGG TGCGGCGGCA GCCAATCAGA GCGGCGCGCT CCGAAAGTTT
      601 CCTTTTATGG CGAGGCGGCG GCGGCGGCGG CCCTATAAAA AGCGAAGCGC GCGGCGGGCG
      661 GGAGTCGCTG CGCGCGCTGC CTTCGCCCCG TGCCCCGCTC CGCGCCGCCT CGCGCCGCCC
      721 GCCCCGGCTC TGACTGACCG CGTTACTCCC ACAGGTGAGC GGGCGGGACG GCCCTTCTCC
      781 TCCGGGCTGT AATTAGCGCT TGGTTTAATG ACGGCTTGTT TCTTTTCTGT GGCTGCGTGA
      841 AAGCCTTGAG GGGCTCCGGG AGGGCCCTTT GTGCGGGGGG GAGCGGCTCG GGGGGTGCGT
      901 GCGTGTGTGT GTGCGTGGGG AGCGCCGCGT GCGGCTCCGC GCTGCCCGGC GGCTGTGAGC
      961 GCTGCGGGCG CGGCGCGGGG CTTTGTGCGC TCCGCAGTGT GCGCGAGGGG AGCGCGGCCG
     1021 GGGGCGGTGC CCCGCGGTGC GGGGGGGGCT GCGAGGGGAA CAAAGGCTGC GTGCGGGGTG
     1081 TGTGCGTGGG GGGGTGAGCA GGGGGTGTGG GCGCGTCGGT CGGGCTGCAA CCCCCCCCTG
     1141 CACCCCCCTC CCCGAGTTGC TGAGCACGGC CCGGCTTCGG GTGCGGGGCT CCGTACGGGG
     1201 CGTGGCGCGG GGCTCGCCGT GCCGGGCGGG GGGTGGCGGC AGGTGGGGGT GCCGGGCGGG
     1261 GCGGGGCCGC CTCGGGCCGG GGAGGGCTCG GGGGAGGGGC GCGGCGGCCC CCGGAGCGCC
     1321 GGCGGCTGTC GAGGCGCGGC GAGCCGCAGC CATTGCCTTT TATGGTAATC GTGCGAGAGG
     1381 GCGCAGGGAC TTCCTTTGTC CCAAATCTGG CGGAGCCGAA ATCTGGGAGG CGCCGCCGCA
     1441 CCCCCTCTAG CGGGCGCGGG GCGAAGCGGT GCGGCGCCGG CAGGAAGGAA ATGGGCGGGG
     1501 AGGGCCTTCG TGCGTCGCCG CGCCGCCGTC CCCTTCTCCC TCTCCAGCCT CGGGGCTGCC
     1561 GCGGGGGGAC GGCTGCCTTC GGGGGGGACG GGGCAGGGCG GGGTTCGGCT TCTGGCGTGT
     1621 GACCGGCGGC TCTAGAGCCT CTGCTAACCA TGTTCATGCC TTCTTCTTTT TCCTACAGCT
     1681 CCTGGGCAAC GTGCTGGTTA TTGTGCTGTC TCATCATTTT GGCAAAGGCC ACCATGTTCG
     1741 TGTTTCTGGT GCTGCTGCCT CTGGTGTCCA GCCAGCGGGT GCAGCCCACC GAATCCATCG
     1801 TGCGGTTCCC CAATATCACC AATCTGTGCC CCTTCGGCGA GGTGTTCAAT GCCACCAGAT
     1861 TCGCCTCTGT GTACGCCTGG AACCGGAAGC GGATCAGCAA TTGCGTGGCC GACTACTCCG
     1921 TGCTGTACAA CTCCGCCAGC TTCAGCACCT TCAAGTGCTA CGGCGTGTCC CCTACCAAGC
     1981 TGAACGACCT GTGCTTCACA AACGTGTACG CCGACAGCTT CGTGATCCGG GGAGATGAAG
     2041 TGCGGCAGAT TGCCCCTGGA CAGACAGGCA AGATCGCCGA CTACAACTAC AAGCTGCCCG
     2101 ACGACTTCAC CGGCTGTGTG ATTGCCTGGA ACAGCAACAA CCTGGACTCC AAAGTCGGCG
     2161 GCAACTACAA TTACCTGTAC CGGCTGTTCC GGAAGTCCAA TCTGAAGCCC TTCGAGCGGG
     2221 ACATCTCCAC CGAGATCTAT CAGGCCGGCA GCACCCCTTG TAACGGCGTG GAAGGCTTCA
     2281 ACTGCTACTT CCCACTGCAG TCCTACGGCT TTCAGCCCAC AAATGGCGTG GGCTATCAGC
     2341 CCTACAGAGT GGTGGTGCTG AGCTTCGAAC TGCTGCATGC CCCTGCCACA GTGTGCGGCC
     2401 CTAAGAAAAG CACCAATCTC GTGAAGAACA AATGCGTGAA CTTCCACCAT CACCATCACC
     2461 ATTGATAAAA TTCGAGCTCG CGGCCGCATC GATCTTAAGT CGCGACTCGA GCTAGCAGAT
     2521 CTTTTTCCCT CTGCCAAAAA TTATGGGGAC ATCATGAAGC CCCTTGAGCA TCTGACTTCT
     2581 GGCTAATAAA GGAAATTTAT TTTCATTGCA ATAGTGTGTT GGAATTTTTT GTGTCTCTCA
     2641 CTCGGAAGGA CATATGGGAG GGCAAATCAT TTAAAACATC AGAATGAGTA TTTGGTTTAG
     2701 AGTTTGGCAA CATATGCCCA TATGCTGGCT GCCATGAACA AAGGTTGGCT ATAAAGAGGT
     2761 CATCAGTATA TGAAACAGCC CCCTGCTGTC CATTCCTTAT TCCATAGAAA AGCCTTGACT
     2821 TGAGGTTAGA TTTTTTTTAT ATTTTGTTTT GTGTTATTTT TTTCTTTAAC ATCCCTAAAA
     2881 TTTTCCTTAC ATGTTTTACT AGCCAGATTT TTCCTCCTCT CCTGACTACT CCCAGTCATA
     2941 GCTGTCCCTC TTCTCTTATG GAGATCCCTC GACCTGCAGC CCAAGCTTGG CGTAATCATG
     3001 GTCATAGCTG TTTCCTGTGT GAAATTGTTA TCCGCTCACA ATTCCACACA ACATACGAGC
     3061 CGGAAGCATA AAGTGTAAAG CCTGGGGTGC CTAATGAGTG AGCTAACTCA CATTAATTGC
     3121 GTTGCGCTCA CTGCCCGCTT TCCAGTCGGG AAACCTGTCG TGCCAGCGGA TCCGCATCTC
     3181 AATTAGTCAG CAACCATAGT CCCGCCCCTA ACTCCGCCCA TCCCGCCCCT AACTCCGCCC
     3241 AGTTCCGCCC ATTCTCCGCC CCATGGCTGA CTAATTTTTT TTATTTATGC AGAGGCCGAG
     3301 GCCGCCTCGG CCTCTGAGCT ATTCCAGAAG TAGTGAGGAG GCTTTTTTGG AGGCCTAGGC
     3361 TTTTGCAAAA AGCTAACTTG TTTATTGCAG CTTATAATGG TTACAAATAA AGCAATAGCA
```

3421 TCACAAATTT CACAAATAAA GCATTTTTTT CACTGCATTC TAGTTGTGGT TTGTCCAAAC
3481 TCATCAATGT ATCTTATCAT GTCTGGATCC GCTGCATTAA TGAATCGGCC AACGCGCGGG
3541 GAGAGGCGGT TTGCGTATTG GGCGCTCTTC CGCTTCCTCG CTCACTGACT CGCTGCGCTC
3601 GGTCGTTCGG CTGCGGCGAG CGGTATCAGC TCACTCAAAG GCGGTAATAC GGTTATCCAC
3661 AGAATCAGGG GATAACGCAG GAAAGAACAT GTGAGCAAAA GGCCAGCAAA AGGCCAGGAA
3721 CCGTAAAAAG GCCGCGTTGC TGGCGTTTTT CCATAGGCTC CGCCCCCCTG ACGAGCATCA
3781 CAAAAATCGA CGCTCAAGTC AGAGGTGGCG AAACCCGACA GGACTATAAA GATACCAGGC
3841 GTTTCCCCCT GGAAGCTCCC TCGTGCGCTC TCCTGTTCCG ACCCTGCCGC TTACCGGATA
3901 CCTGTCCGCC TTTCTCCCTT CGGGAAGCGT GGCGCTTTCT CATAGCTCAC GCTGTAGGTA
3961 TCTCAGTTCG GTGTAGGTCG TTCGCTCCAA GCTGGGCTGT GTGCACGAAC CCCCCGTTCA
4021 GCCCGACCGC TGCGCCTTAT CCGGTAACTA TCGTCTTGAG TCCAACCCGG TAAGACACGA
4081 CTTATCGCCA CTGGCAGCAG CCACTGGTAA CAGGATTAGC AGAGCGAGGT ATGTAGGCGG
4141 TGCTACAGAG TTCTTGAAGT GGTGGCCTAA CTACGGCTAC ACTAGAAGAA CAGTATTTGG
4201 TATCTGCGCT CTGCTGAAGC CAGTTACCTT CGGAAAAAGA GTTGGTAGCT CTTGATCCGG
4261 CAAACAAACC ACCGCTGGTA GCGGTGGTTT TTTTGTTTGC AAGCAGCAGA TTACGCGCAG
4321 AAAAAAAGGA TCTCAAGAAG ATCCTTTGAT CTTTTCTACG GGGTCTGACG CTCAGTGGAA
4381 CGAAAACTCA CGTTAAGGGA TTTTGGTCAT GAGATTATCA AAAAGGATCT TCACCTAGAT
4441 CCTTTTAAAT TAAAAATGAA GTTTTAAATC AATCTAAAGT ATATATGAGT AAACTTGGTC
4501 TGACAGTTAC CAATGCTTAA TCAGTGAGGC ACCTATCTCA GCGATCTGTC TATTTCGTTC
4561 ATCCATAGTT GCCTGACTCC CCGTCGTGTA GATAACTACG ATACGGGAGG GCTTACCATC
4621 TGGCCCCAGT GCTGCAATGA TACCGCGAGA CCCACGCTCA CCGGCTCCAG ATTTATCAGC
4681 AATAAACCAG CCAGCCGGAA GGGCCGAGCG CAGAAGTGGT CCTGCAACTT TATCCGCCTC
4741 CATCCAGTCT ATTAATTGTT GCCGGGAAGC TAGAGTAAGT AGTTCGCCAG TTAATAGTTT
4801 GCGCAACGTT GTTGCCATTG CTACAGGCAT CGTGGTGTCA CGCTCGTCGT TTGGTATGGC
4861 TTCATTCAGC TCCGGTTCCC AACGATCAAG GCGAGTTACA TGATCCCCCA TGTTGTGCAA
4921 AAAAGCGGTT AGCTCCTTCG GTCCTCCGAT CGTTGTCAGA AGTAAGTTGG CCGCAGTGTT
4981 ATCACTCATG GTTATGGCAG CACTGCATAA TTCTCTTACT GTCATGCCAT CCGTAAGATG
5041 CTTTTCTGTG ACTGGTGAGT ACTCAACCAA GTCATTCTGA GAATAGTGTA TGCGGCGACC
5101 GAGTTGCTCT TGCCCGGCGT CAATACGGGA TAATACCGCG CCACATAGCA GAACTTTAAA
5161 AGTGCTCATC ATTGGAAAAC GTTCTTCGGG GCGAAAACTC TCAAGGATCT TACCGCTGTT
5221 GAGATCCAGT TCGATGTAAC CCACTCGTGC ACCCAACTGA TCTTCAGCAT CTTTTACTTT
5281 CACCAGCGTT TCTGGGTGAG CAAAAACAGG AAGGCAAAAT GCCGCAAAAA AGGGAATAAG
5341 GGCGACACGG AAATGTTGAA TACTCATACT CTTCCTTTTT CAATATTATT GAAGCATTTA
5401 TCAGGGTTAT TGTCTCATGA GCGGATACAT ATTTGAATGT ATTTAGAAAA ATAAACAAAT
5461 AGGGGTTCCG CGCACATTTC CCCGAAAAGT GCCACCTG
//

```
>pSL1515_SARS-CoV-2-Spike-RBD-WithSpyTag_Expression Plasmid
LOCUS       pSL1515               5512 bp    DNA       circular     12-MAY-2022
DEFINITION  .
ACCESSION
VERSION
SOURCE      .
  ORGANISM  .
COMMENT
COMMENT     ApEinfo:methylated:1
FEATURES             Location/Qualifiers
     primer_bind     complement(3011..3031)
                     /locus_tag="M13-rev"
                     /label="M13-rev"
                     /ApEinfo_label="M13-rev"
                     /ApEinfo_fwdcolor="cyan"
                     /ApEinfo_revcolor="green"
                     /ApEinfo_graphicformat="arrow_data {{0 1 2 0 0 -1} {} 0}
                     width 5 offset 0"
     misc_feature    1898..2462
                     /locus_tag="SARS-CoV-2 Spike SP+RBD+6xHis"
                     /label="SARS-CoV-2 Spike SP+RBD+6xHis"
                     /ApEinfo_label="SARS-CoV-2 Spike SP+RBD+6xHis"
                     /ApEinfo_fwdcolor="cyan"
                     /ApEinfo_revcolor="green"
                     /ApEinfo_graphicformat="arrow_data {{0 1 2 0 0 -1} {} 0}
                     width 5 offset 0"
     rep_origin      complement(3744..4426)
                     /locus_tag="ColE1 origin"
                     /label="ColE1 origin"
                     /ApEinfo_label="ColE1 origin"
                     /ApEinfo_fwdcolor="gray50"
                     /ApEinfo_revcolor="gray50"
                     /ApEinfo_graphicformat="arrow_data {{0 1 2 0 0 -1} {} 0}
                     width 5 offset 0"
     misc_binding    complement(3037..3059)
                     /locus_tag="LacO"
                     /label="LacO"
                     /ApEinfo_label="LacO"
                     /ApEinfo_fwdcolor="#6495ed"
                     /ApEinfo_revcolor="#6495ed"
                     /ApEinfo_graphicformat="arrow_data {{0 1 2 0 0 -1} {} 0}
                     width 5 offset 0"
     misc_feature    3389..3519
                     /locus_tag="SV40 early poly-A signal"
                     /label="SV40 early poly-A signal"
                     /ApEinfo_label="SV40 early poly-A signal"
                     /ApEinfo_fwdcolor="#808000"
                     /ApEinfo_revcolor="green"
                     /ApEinfo_graphicformat="arrow_data {{0 1 2 0 0 -1} {} 0}
                     width 5 offset 0"
     misc_feature    complement(4524..5183)
                     /locus_tag="AmpR"
                     /label="AmpR"
                     /ApEinfo_label="AmpR"
                     /ApEinfo_fwdcolor="#ffff00"
                     /ApEinfo_revcolor="#ffff00"
                     /ApEinfo_graphicformat="arrow_data {{0 1 2 0 0 -1} {} 0}
                     width 5 offset 0"
```

misc_feature 3199..3378
/locus_tag="SV40 Promoter"
/label="SV40 Promoter"
/ApEinfo_label="SV40 Promoter"
/ApEinfo_fwdcolor="#8080c0"
/ApEinfo_revcolor="green"
/ApEinfo_graphicformat="arrow_data {{0 1 2 0 0 -1} {} 0}
width 5 offset 0"
misc_feature complement(776..781)
/locus_tag="Ribosome Binding Site"
/label="Ribosome Binding Site"
/ApEinfo_label="Ribosome Binding Site"
/ApEinfo_fwdcolor="cyan"
/ApEinfo_revcolor="green"
/ApEinfo_graphicformat="arrow_data {{0 1 2 0 0 -1} {} 0}
width 5 offset 0"
misc_feature complement(2932..2937)
/locus_tag="Ribosome Binding Site(1)"
/label="Ribosome Binding Site(1)"
/ApEinfo_label="Ribosome Binding Site"
/ApEinfo_fwdcolor="cyan"
/ApEinfo_revcolor="green"
/ApEinfo_graphicformat="arrow_data {{0 1 2 0 0 -1} {} 0}
width 5 offset 0"
misc_feature complement(3883..3888)
/locus_tag="Ribosome Binding Site(2)"
/label="Ribosome Binding Site(2)"
/ApEinfo_label="Ribosome Binding Site"
/ApEinfo_fwdcolor="cyan"
/ApEinfo_revcolor="green"
/ApEinfo_graphicformat="arrow_data {{0 1 2 0 0 -1} {} 0}
width 5 offset 0"
misc_feature complement(3037..3054)
/locus_tag="Minimal LacO"
/label="Minimal LacO"
/ApEinfo_label="Minimal LacO"
/ApEinfo_fwdcolor="#ff80c0"
/ApEinfo_revcolor="green"
/ApEinfo_graphicformat="arrow_data {{0 1 2 0 0 -1} {} 0}
width 5 offset 0"
misc_feature 84..371
/locus_tag="CAG Enhancer / CMV immediate early promoter"
/label="CAG Enhancer / CMV immediate early promoter"
/ApEinfo_label="CAG Enhancer / CMV immediate early
promoter"
/ApEinfo_fwdcolor="#8080c0"
/ApEinfo_revcolor="green"
/ApEinfo_graphicformat="arrow_data {{0 1 2 0 0 -1} {} 0}
width 5 offset 0"
misc_feature 2531..2969
/locus_tag="rb glob PA terminator"
/label="rb glob PA terminator"
/ApEinfo_label="rb glob PA terminator"
/ApEinfo_fwdcolor="#800000"
/ApEinfo_revcolor="green"
/ApEinfo_graphicformat="arrow_data {{0 1 2 0 0 -1} {} 0}
width 5 offset 0"
misc_feature 1734..1897

```
                     /locus_tag="SARS-CoV-2 Spike SP+RBD+6xHis(1)"
                     /label="SARS-CoV-2 Spike SP+RBD+6xHis(1)"
                     /ApEinfo_label="SARS-CoV-2 Spike SP+RBD+6xHis"
                     /ApEinfo_fwdcolor="cyan"
                     /ApEinfo_revcolor="green"
                     /ApEinfo_graphicformat="arrow_data {{0 1 2 0 0 -1} {} 0}
                     width 5 offset 0"
     misc_feature    385..1630
                     /locus_tag="CAG promoter"
                     /label="CAG promoter"
                     /ApEinfo_label="CAG promoter"
                     /ApEinfo_fwdcolor="#0080c0"
                     /ApEinfo_revcolor="green"
                     /ApEinfo_graphicformat="arrow_data {{0 1 2 0 0 -1} {} 0}
                     width 5 offset 0"
ORIGIN
        1 GTCGACATTG ATTATTGACT AGTTATTAAT AGTAATCAAT TACGGGGTCA TTAGTTCATA
       61 GCCCATATAT GGAGTTCCGC GTTACATAAC TTACGGTAAA TGGCCCGCCT GGCTGACCGC
      121 CCAACGACCC CCGCCCATTG ACGTCAATAA TGACGTATGT TCCCATAGTA ACGCCAATAG
      181 GGACTTTCCA TTGACGTCAA TGGGTGGAGT ATTTACGGTA AACTGCCCAC TTGGCAGTAC
      241 ATCAAGTGTA TCATATGCCA AGTACGCCCC CTATTGACGT CAATGACGGT AAATGGCCCG
      301 CCTGGCATTA TGCCCAGTAC ATGACCTTAT GGGACTTTCC TACTTGGCAG TACATCTACG
      361 TATTAGTCAT CGCTATTACC ATGGTCGAGG TGAGCCCCAC GTTCTGCTTC ACTCTCCCCA
      421 TCTCCCCCCC CTCCCCACCC CCAATTTTGT ATTTATTTAT TTTTTAATTA TTTTGTGCAG
      481 CGATGGGGGC GGGGGGGGGG GGGGCGCGCG CCAGGCGGGG CGGGGCGGGG CGAGGGGCGG
      541 GGCGGGGCGA GGCGGAGAGG TGCGGCGGCA GCCAATCAGA GCGGCGCGCT CCGAAAGTTT
      601 CCTTTTATGG CGAGGCGGCG GCGGCGGCGG CCCTATAAAA AGCGAAGCGC GCGGCGGGCG
      661 GGAGTCGCTG CGCGCGCTGC CTTCGCCCCG TGCCCCGCTC CGCGCCGCCT CGCGCCGCCC
      721 GCCCCGGCTC TGACTGACCG CGTTACTCCC ACAGGTGAGC GGGCGGGACG GCCCTTCTCC
      781 TCCGGGCTGT AATTAGCGCT TGGTTTAATG ACGGCTTGTT TCTTTTCTGT GGCTGCGTGA
      841 AAGCCTTGAG GGGCTCCGGG AGGGCCCTTT GTGCGGGGGG GAGCGGCTCG GGGGGTGCGT
      901 GCGTGTGTGT GTGCGTGGGG AGCGCCGCGT GCGGCTCCGC GCTGCCCGGC GGCTGTGAGC
      961 GCTGCGGGCG CGGCGCGGGG CTTTGTGCGC TCCGCAGTGT GCGCGAGGGG AGCGCGGCCG
     1021 GGGGCGGTGC CCCGCGGTGC GGGGGGGGCT GCGAGGGGAA CAAAGGCTGC GTGCGGGGTG
     1081 TGTGCGTGGG GGGGTGAGCA GGGGGTGTGG GCGCGTCGGT CGGGCTGCAA CCCCCCCCTG
     1141 CACCCCCCTC CCCGAGTTGC TGAGCACGGC CCGGCTTCGG GTGCGGGGCT CCGTACGGGG
     1201 CGTGGCGCGG GGCTCGCCGT GCCGGGCGGG GGGTGGCGGC AGGTGGGGGT GCCGGGCGGG
     1261 GCGGGGCCGC CTCGGGCCGG GGAGGGCTCG GGGGAGGGGC GCGGCGGCCC CCGGAGCGCC
     1321 GGCGGCTGTC GAGGCGCGGC GAGCCGCAGC CATTGCCTTT TATGGTAATC GTGCGAGAGG
     1381 GCGCAGGGAC TTCCTTTGTC CCAAATCTGG CGGAGCCGAA ATCTGGGAGG CGCCGCCGCA
     1441 CCCCCTCTAG CGGGCGCGGG GCGAAGCGGT GCGGCGCCGG CAGGAAGGAA ATGGGCGGGG
     1501 AGGGCCTTCG TGCGTCGCCG CGCCGCCGTC CCCTTCTCCC TCTCCAGCCT CGGGGCTGCC
     1561 GCGGGGGGAC GGCTGCCTTC GGGGGGGACG GGGCAGGGCG GGGTTCGGCT TCTGGCGTGT
     1621 GACCGGCGGC TCTAGAGCCT CTGCTAACCA TGTTCATGCC TTCTTCTTTT TCCTACAGCT
     1681 CCTGGGCAAC GTGCTGGTTA TTGTGCTGTC TCATCATTTT GGCAAAGGCC ACCATGTTCG
     1741 TGTTTCTGGT GCTGCTGCCT CTGGTGTCCA GCCAGCGGGT GCAGCCCACC GAATCCATCG
     1801 TGCGGTTCCC CAATATCACC AATCTGTGCC CCTTCGGCGA GGTGTTCAAT GCCACCAGAT
     1861 TCGCCTCTGT GTACGCCTGG AACCGGAAGC GGATCAGCAA TTGCGTGGCC GACTACTCCG
     1921 TGCTGTACAA CTCCGCCAGC TTCAGCACCT TCAAGTGCTA CGGCGTGTCC CCTACCAAGC
     1981 TGAACGACCT GTGCTTCACA AACGTGTACG CCGACAGCTT CGTGATCCGG GGAGATGAAG
     2041 TGCGGCAGAT TGCCCCTGGA CAGACAGGCA AGATCGCCGA CTACAACTAC AAGCTGCCCG
     2101 ACGACTTCAC CGGCTGTGTG ATTGCCTGGA ACAGCAACAA CCTGGACTCC AAAGTCGGCG
     2161 GCAACTACAA TTACCTGTAC CGGCTGTTCC GGAAGTCCAA TCTGAAGCCC TTCGAGCGGG
     2221 ACATCTCCAC CGAGATCTAT CAGGCCGGCA GCACCCCTTG TAACGGCGTG GAAGGCTTCA
     2281 ACTGCTACTT CCCACTGCAG TCCTACGGCT TTCAGCCCAC AAATGGCGTG GGCTATCAGC
     2341 CCTACAGAGT GGTGGTGCTG AGCTTCGAAC TGCTGCATGC CCCTGCCACA GTGTGCGGCC
     2401 CTAAGAAAAG CACCAATCTC GTGAAGAACA AATGCGTGAA CTTCCACCAT CACCATCACC
     2461 ATggttccgg tggagcacat attgtgatgg ttgacgctta caagccaacc aaataatgac
```

2521 tcgagCTAGC AGATCTTTTT CCCTCTGCCA AAAATTATGG GGACATCATG AAGCCCCTTG
2581 AGCATCTGAC TTCTGGCTAA TAAAGGAAAT TTATTTTCAT TGCAATAGTG TGTTGGAATT
2641 TTTTGTGTCT CTCACTCGGA AGGACATATG GGAGGGCAAA TCATTTAAAA CATCAGAATG
2701 AGTATTTGGT TTAGAGTTTG GCAACATATG CCCATATGCT GGCTGCCATG AACAAAGGTT
2761 GGCTATAAAG AGGTCATCAG TATATGAAAC AGCCCCCTGC TGTCCATTCC TTATTCCATA
2821 GAAAAGCCTT GACTTGAGGT TAGATTTTTT TTATATTTTG TTTTGTGTTA TTTTTTTCTT
2881 TAACATCCCT AAAATTTTCC TTACATGTTT TACTAGCCAG ATTTTTCCTC CTCTCCTGAC
2941 TACTCCCAGT CATAGCTGTC CCTCTTCTCT TATGGAGATC CCTCGACCTG CAGCCCAAGC
3001 TTGGCGTAAT CATGGTCATA GCTGTTTCCT GTGTGAAATT GTTATCCGCT CACAATTCCA
3061 CACAACATAC GAGCCGGAAG CATAAAGTGT AAAGCCTGGG GTGCCTAATG AGTGAGCTAA
3121 CTCACATTAA TTGCGTTGCG CTCACTGCCC GCTTTCCAGT CGGGAAACCT GTCGTGCCAG
3181 CGGATCCGCA TCTCAATTAG TCAGCAACCA TAGTCCCGCC CCTAACTCCG CCCATCCCGC
3241 CCCTAACTCC GCCCAGTTCC GCCCATTCTC CGCCCCATGG CTGACTAATT TTTTTTATTT
3301 ATGCAGAGGC CGAGGCCGCC TCGGCCTCTG AGCTATTCCA GAAGTAGTGA GGAGGCTTTT
3361 TTGGAGGCCT AGGCTTTTGC AAAAAGCTAA CTTGTTTATT GCAGCTTATA ATGGTTACAA
3421 ATAAAGCAAT AGCATCACAA ATTTCACAAA TAAAGCATTT TTTTCACTGC ATTCTAGTTG
3481 TGGTTTGTCC AAACTCATCA ATGTATCTTA TCATGTCTGG ATCCGCTGCA TTAATGAATC
3541 GGCCAACGCG CGGGGAGAGG CGGTTTGCGT ATTGGGCGCT CTTCCGCTTC CTCGCTCACT
3601 GACTCGCTGC GCTCGGTCGT TCGGCTGCGG CGAGCGGTAT CAGCTCACTC AAAGGCGGTA
3661 ATACGGTTAT CCACAGAATC AGGGGATAAC GCAGGAAAGA ACATGTGAGC AAAAGGCCAG
3721 CAAAAGGCCA GGAACCGTAA AAAGGCCGCG TTGCTGGCGT TTTTCCATAG GCTCCGCCCC
3781 CCTGACGAGC ATCACAAAAA TCGACGCTCA AGTCAGAGGT GGCGAAACCC GACAGGACTA
3841 TAAAGATACC AGGCGTTTCC CCCTGGAAGC TCCCTCGTGC GCTCTCCTGT TCCGACCCTG
3901 CCGCTTACCG GATACCTGTC CGCCTTTCTC CCTTCGGGAA GCGTGGCGCT TTCTCATAGC
3961 TCACGCTGTA GGTATCTCAG TTCGGTGTAG GTCGTTCGCT CCAAGCTGGG CTGTGTGCAC
4021 GAACCCCCCG TTCAGCCCGA CCGCTGCGCC TTATCCGGTA ACTATCGTCT TGAGTCCAAC
4081 CCGGTAAGAC ACGACTTATC GCCACTGGCA GCAGCCACTG GTAACAGGAT TAGCAGAGCG
4141 AGGTATGTAG GCGGTGCTAC AGAGTTCTTG AAGTGGTGGC CTAACTACGG CTACACTAGA
4201 AGAACAGTAT TTGGTATCTG CGCTCTGCTG AAGCCAGTTA CCTTCGGAAA AAGAGTTGGT
4261 AGCTCTTGAT CCGGCAAACA AACCACCGCT GGTAGCGGTG GTTTTTTTGT TTGCAAGCAG
4321 CAGATTACGC GCAGAAAAAA AGGATCTCAA GAAGATCCTT TGATCTTTTC TACGGGGTCT
4381 GACGCTCAGT GGAACGAAAA CTCACGTTAA GGGATTTTGG TCATGAGATT ATCAAAAAGG
4441 ATCTTCACCT AGATCCTTTT AAATTAAAAA TGAAGTTTTA AATCAATCTA AAGTATATAT
4501 GAGTAAACTT GGTCTGACAG TTACCAATGC TTAATCAGTG AGGCACCTAT CTCAGCGATC
4561 TGTCTATTTC GTTCATCCAT AGTTGCCTGA CTCCCCGTCG TGTAGATAAC TACGATACGG
4621 GAGGGCTTAC CATCTGGCCC CAGTGCTGCA ATGATACCGC GAGACCCACG CTCACCGGCT
4681 CCAGATTTAT CAGCAATAAA CCAGCCAGCC GGAAGGGCCG AGCGCAGAAG TGGTCCTGCA
4741 ACTTTATCCG CCTCCATCCA GTCTATTAAT TGTTGCCGGG AAGCTAGAGT AAGTAGTTCG
4801 CCAGTTAATA GTTTGCGCAA CGTTGTTGCC ATTGCTACAG GCATCGTGGT GTCACGCTCG
4861 TCGTTTGGTA TGGCTTCATT CAGCTCCGGT TCCCAACGAT CAAGGCGAGT TACATGATCC
4921 CCCATGTTGT GCAAAAAAGC GGTTAGCTCC TTCGGTCCTC CGATCGTTGT CAGAAGTAAG
4981 TTGGCCGCAG TGTTATCACT CATGGTTATG GCAGCACTGC ATAATTCTCT TACTGTCATG
5041 CCATCCGTAA GATGCTTTTC TGTGACTGGT GAGTACTCAA CCAAGTCATT CTGAGAATAG
5101 TGTATGCGGC GACCGAGTTG CTCTTGCCCG GCGTCAATAC GGGATAATAC CGCGCCACAT
5161 AGCAGAACTT TAAAAGTGCT CATCATTGGA AAACGTTCTT CGGGGCGAAA ACTCTCAAGG
5221 ATCTTACCGC TGTTGAGATC CAGTTCGATG TAACCCACTC GTGCACCCAA CTGATCTTCA
5281 GCATCTTTTA CTTTCACCAG CGTTTCTGGG TGAGCAAAAA CAGGAAGGCA AAATGCCGCA
5341 AAAAAGGGAA TAAGGGCGAC ACGGAAATGT TGAATACTCA TACTCTTCCT TTTTCAATAT
5401 TATTGAAGCA TTTATCAGGG TTATTGTCTC ATGAGCGGAT ACATATTTGA ATGTATTTAG
5461 AAAAATAAAC AAATAGGGGT TCCGCGCACA TTTCCCCGAA AAGTGCCACC TG

//

### Supp File 2 - Cryo-EM Validation Report

Full wwPDB EM Validation Report (i)

Aug 9, 2022 – 12:48 PM EDT

| | | |
|---|---|---|
| PDB ID | : | 8E01 |
| EMDB ID | : | EMD-27812 |
| Title | : | Structure of engineered nano-cage fusion protein |
| Deposited on | : | 2022-08-08 |
| Resolution | : | 3.40 Å(reported) |

**This wwPDB validation report is for manuscript review**

For Manuscript Review

This is a Full wwPDB EM Validation Report.

This report is produced by the wwPDB biocuration pipeline after annotation of the structure.

We welcome your comments at **
A user guide is available at
https://www.wwpdb.org/validation/2017/EMValidationReportHelp
with specific help available everywhere you see the (i) symbol.

The types of validation reports are described at http://www.wwpdb.org/validation/2017/FAQs#types.

The following versions of software and data (see references (i)) were used in the production of this report:

| | | |
|---|---|---|
| EMDB validation analysis | : | 0.0.1.dev8 |
| MolProbity | : | 4.02b-467 |
| Percentile statistics | : | 20191225.v01 (using entries in the PDB archive December 25th 2019) |
| Ideal geometry (proteins) | : | Engh & Huber (2001) |
| Ideal geometry (DNA, RNA) | : | Parkinson et al. (1996) |
| Validation Pipeline (wwPDB-VP) | : | 2.29 |

### 1 Overall quality at a glance (i)

The following experimental techniques were used to determine the structure:
*ELECTRON MICROSCOPY*

The reported resolution of this entry is 3.40 Å.

Percentile scores (ranging between 0-100) for global validation metrics of the entry are shown in the following graphic. The table shows the number of entries on which the scores are based.

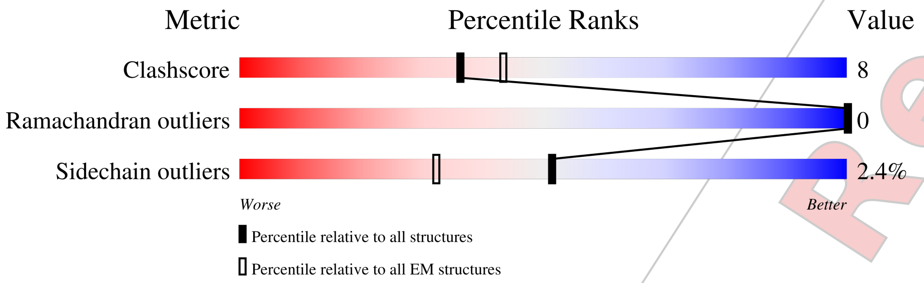

| Metric | Whole archive (#Entries) | EM structures (#Entries) |
|---|---|---|
| Clashscore | 158937 | 4297 |
| Ramachandran outliers | 154571 | 4023 |
| Sidechain outliers | 154315 | 3826 |

The table below summarises the geometric issues observed across the polymeric chains and their fit to the map. The red, orange, yellow and green segments of the bar indicate the fraction of residues that contain outliers for >=3, 2, 1 and 0 types of geometric quality criteria respectively. A grey segment represents the fraction of residues that are not modelled. The numeric value for each fraction is indicated below the corresponding segment, with a dot representing fractions <=5% The upper red bar (where present) indicates the fraction of residues that have poor fit to the EM map (all-atom inclusion < 40%). The numeric value is given above the bar.

| Mol | Chain | Length | Quality of chain |
|---|---|---|---|
| 1 | A | 224 | 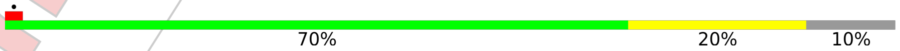  |

For Manuscript Review

#### 2 Entry composition (i)

There is only 1 type of molecule in this entry. The entry contains 1518 atoms, of which 0 are hydrogens and 0 are deuteriums.

In the tables below, the AltConf column contains the number of residues with at least one atom in alternate conformation and the Trace column contains the number of residues modelled with at most 2 atoms.

- Molecule 1 is a protein called 2-dehydro-3-deoxyphosphogluconate aldolase/4-hydroxy-2-oxoglutarate aldolase.

| Mol | Chain | Residues | Atoms | | | | | AltConf | Trace |
|---|---|---|---|---|---|---|---|---|---|
| | | | Total | C | N | O | S | | |
| 1 | A | 201 | 1518 | 988 | 248 | 272 | 10 | 0 | 0 |

There are 24 discrepancies between the modelled and reference sequences:

| Chain | Residue | Modelled | Actual | Comment | Reference |
|---|---|---|---|---|---|
| A | 1 | MET | - | initiating methionine | UNP Q9WXS1 |
| A | 2 | HIS | - | expression tag | UNP Q9WXS1 |
| A | 3 | HIS | - | expression tag | UNP Q9WXS1 |
| A | 4 | HIS | - | expression tag | UNP Q9WXS1 |
| A | 5 | HIS | - | expression tag | UNP Q9WXS1 |
| A | 6 | HIS | - | expression tag | UNP Q9WXS1 |
| A | 7 | HIS | - | expression tag | UNP Q9WXS1 |
| A | 8 | GLY | - | expression tag | UNP Q9WXS1 |
| A | 9 | GLY | - | expression tag | UNP Q9WXS1 |
| A | 10 | SER | - | expression tag | UNP Q9WXS1 |
| A | 11 | GLY | - | expression tag | UNP Q9WXS1 |
| A | 12 | GLY | - | expression tag | UNP Q9WXS1 |
| A | 13 | SER | - | expression tag | UNP Q9WXS1 |
| A | 14 | GLY | - | expression tag | UNP Q9WXS1 |
| A | 15 | GLY | - | expression tag | UNP Q9WXS1 |
| A | 16 | SER | - | expression tag | UNP Q9WXS1 |
| A | 17 | GLY | - | expression tag | UNP Q9WXS1 |
| A | 18 | GLY | - | expression tag | UNP Q9WXS1 |
| A | 19 | SER | - | expression tag | UNP Q9WXS1 |
| A | 45 | LYS | GLU | conflict | UNP Q9WXS1 |
| A | 52 | LEU | GLU | conflict | UNP Q9WXS1 |
| A | 80 | MET | LYS | conflict | UNP Q9WXS1 |
| A | 206 | VAL | ASP | conflict | UNP Q9WXS1 |
| A | 209 | ALA | ARG | conflict | UNP Q9WXS1 |

For Manuscript Review

### 3 Residue-property plots ⓘ

These plots are drawn for all protein, RNA, DNA and oligosaccharide chains in the entry. The first graphic for a chain summarises the proportions of the various outlier classes displayed in the second graphic. The second graphic shows the sequence view annotated by issues in geometry and atom inclusion in map density. Residues are color-coded according to the number of geometric quality criteria for which they contain at least one outlier: green = 0, yellow = 1, orange = 2 and red = 3 or more. A red diamond above a residue indicates a poor fit to the EM map for this residue (all-atom inclusion < 40%). Stretches of 2 or more consecutive residues without any outlier are shown as a green connector. Residues present in the sample, but not in the model, are shown in grey.

- Molecule 1: 2-dehydro-3-deoxyphosphogluconate aldolase/4-hydroxy-2-oxoglutarate aldolase

Chain A: 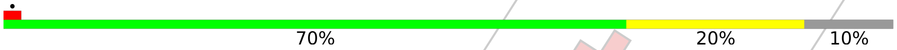

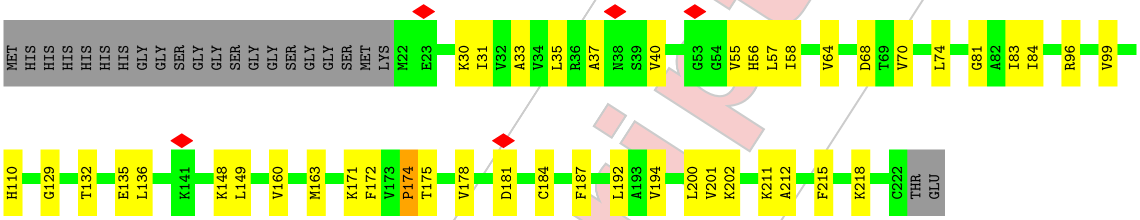

For Manuscript Review

### 4 Experimental information (i)

| Property | Value | Source |
|---|---|---|
| EM reconstruction method | SINGLE PARTICLE | Depositor |
| Imposed symmetry | POINT, I | Depositor |
| Number of particles used | 63430 | Depositor |
| Resolution determination method | FSC 0.143 CUT-OFF | Depositor |
| CTF correction method | PHASE FLIPPING AND AMPLITUDE CORRECTION | Depositor |
| Microscope | FEI TITAN KRIOS | Depositor |
| Voltage (kV) | 300 | Depositor |
| Electron dose ($e^-/\text{Å}^2$) | 44.85 | Depositor |
| Minimum defocus (nm) | 1200 | Depositor |
| Maximum defocus (nm) | 3000 | Depositor |
| Magnification | 59000 | Depositor |
| Image detector | FEI FALCON III (4k x 4k) | Depositor |
| Maximum map value | 6.362 | Depositor |
| Minimum map value | -4.024 | Depositor |
| Average map value | 0.007 | Depositor |
| Map value standard deviation | 0.216 | Depositor |
| Recommended contour level | 0.9 | Depositor |
| Map size (Å) | 462.0, 462.0, 462.0 | wwPDB |
| Map dimensions | 420, 420, 420 | wwPDB |
| Map angles (°) | 90.0, 90.0, 90.0 | wwPDB |
| Pixel spacing (Å) | 1.1, 1.1, 1.1 | Depositor |

For Manuscript Review

### 5 Model quality (i)

#### 5.1 Standard geometry (i)

The Z score for a bond length (or angle) is the number of standard deviations the observed value is removed from the expected value. A bond length (or angle) with $|Z| > 5$ is considered an outlier worth inspection. RMSZ is the root-mean-square of all Z scores of the bond lengths (or angles).

| Mol | Chain | Bond lengths | | Bond angles | |
|---|---|---|---|---|---|
| | | RMSZ | #\|Z\| >5 | RMSZ | #\|Z\| >5 |
| 1 | A | 0.25 | 0/1547 | 0.45 | 0/2086 |

There are no bond length outliers.

There are no bond angle outliers.

There are no chirality outliers.

There are no planarity outliers.

#### 5.2 Too-close contacts (i)

In the following table, the Non-H and H(model) columns list the number of non-hydrogen atoms and hydrogen atoms in the chain respectively. The H(added) column lists the number of hydrogen atoms added and optimized by MolProbity. The Clashes column lists the number of clashes within the asymmetric unit, whereas Symm-Clashes lists symmetry-related clashes.

| Mol | Chain | Non-H | H(model) | H(added) | Clashes | Symm-Clashes |
|---|---|---|---|---|---|---|
| 1 | A | 1518 | 0 | 1587 | 26 | 0 |
| All | All | 1518 | 0 | 1587 | 26 | 0 |

The all-atom clashscore is defined as the number of clashes found per 1000 atoms (including hydrogen atoms). The all-atom clashscore for this structure is 8.

All (26) close contacts within the same asymmetric unit are listed below, sorted by their clash magnitude.

| Atom-1 | Atom-2 | Interatomic distance (Å) | Clash overlap (Å) |
|---|---|---|---|
| 1:A:202:LYS:O | 1:A:211:LYS:NZ | 2.31 | 0.60 |
| 1:A:148:LYS:NZ | 1:A:175:THR:OG1 | 2.35 | 0.59 |
| 1:A:171:LYS:HD2 | 1:A:192:LEU:HD22 | 1.85 | 0.58 |
| 1:A:136:LEU:HD11 | 1:A:172:PHE:HZ | 1.72 | 0.55 |
| 1:A:178:VAL:HG11 | 1:A:194:VAL:HG11 | 1.89 | 0.55 |

Continued on next page...

For Manuscript Review

*Continued from previous page...*

| Atom-1 | Atom-2 | Interatomic distance (Å) | Clash overlap (Å) |
|---|---|---|---|
| 1:A:37:ALA:H | 1:A:64:VAL:HG22 | 1.73 | 0.54 |
| 1:A:35:LEU:HA | 1:A:201:VAL:HG11 | 1.96 | 0.48 |
| 1:A:160:VAL:HG12 | 1:A:172:PHE:CD2 | 2.48 | 0.48 |
| 1:A:33:ALA:HB2 | 1:A:55:VAL:HG11 | 1.94 | 0.48 |
| 1:A:56:HIS:ND1 | 1:A:81:GLY:O | 2.28 | 0.48 |
| 1:A:31:ILE:HG21 | 1:A:55:VAL:HG22 | 1.97 | 0.47 |
| 1:A:149:LEU:HB3 | 1:A:174:PRO:HA | 1.98 | 0.46 |
| 1:A:149:LEU:HD23 | 1:A:174:PRO:HB3 | 1.98 | 0.45 |
| 1:A:184:CYS:SG | 1:A:218:LYS:NZ | 2.89 | 0.45 |
| 1:A:132:THR:OG1 | 1:A:135:GLU:OE2 | 2.35 | 0.45 |
| 1:A:200:LEU:HD11 | 1:A:212:ALA:HA | 1.98 | 0.44 |
| 1:A:31:ILE:HG22 | 1:A:55:VAL:HG13 | 2.00 | 0.44 |
| 1:A:57:LEU:HD23 | 1:A:83:ILE:HB | 2.00 | 0.44 |
| 1:A:68:ASP:OD2 | 1:A:68:ASP:N | 2.51 | 0.43 |
| 1:A:163:MET:O | 1:A:163:MET:HG3 | 2.19 | 0.43 |
| 1:A:40:VAL:HG13 | 1:A:70:VAL:HG22 | 2.01 | 0.42 |
| 1:A:129:GLY:HA2 | 1:A:148:LYS:HB3 | 2.01 | 0.42 |
| 1:A:30:LYS:HB3 | 1:A:187:PHE:HZ | 1.84 | 0.42 |
| 1:A:58:ILE:O | 1:A:84:ILE:HA | 2.20 | 0.42 |
| 1:A:74:LEU:O | 1:A:74:LEU:HD23 | 2.21 | 0.40 |
| 1:A:96:ARG:HA | 1:A:99:VAL:HG22 | 2.02 | 0.40 |

There are no symmetry-related clashes.

#### 5.3 Torsion angles (i)

##### 5.3.1 Protein backbone (i)

In the following table, the Percentiles column shows the percent Ramachandran outliers of the chain as a percentile score with respect to all PDB entries followed by that with respect to all EM entries.

The Analysed column shows the number of residues for which the backbone conformation was analysed, and the total number of residues.

| Mol | Chain | Analysed | Favoured | Allowed | Outliers | Percentiles | |
|---|---|---|---|---|---|---|---|
| 1 | A | 199/224 (89%) | 195 (98%) | 4 (2%) | 0 | 100 | 100 |

There are no Ramachandran outliers to report.

For Manuscript Review

##### 5.3.2 Protein sidechains (i)

In the following table, the Percentiles column shows the percent sidechain outliers of the chain as a percentile score with respect to all PDB entries followed by that with respect to all EM entries.

The Analysed column shows the number of residues for which the sidechain conformation was analysed, and the total number of residues.

| Mol | Chain | Analysed | Rotameric | Outliers | Percentiles | |
|---|---|---|---|---|---|---|
| 1 | A | 164/179 (92%) | 160 (98%) | 4 (2%) | 49 | 74 |

All (4) residues with a non-rotameric sidechain are listed below:

| Mol | Chain | Res | Type |
|---|---|---|---|
| 1 | A | 110 | HIS |
| 1 | A | 174 | PRO |
| 1 | A | 181 | ASP |
| 1 | A | 215 | PHE |

Sometimes sidechains can be flipped to improve hydrogen bonding and reduce clashes. There are no such sidechains identified.

##### 5.3.3 RNA (i)

There are no RNA molecules in this entry.

#### 5.4 Non-standard residues in protein, DNA, RNA chains (i)

There are no non-standard protein/DNA/RNA residues in this entry.

#### 5.5 Carbohydrates (i)

There are no monosaccharides in this entry.

#### 5.6 Ligand geometry (i)

There are no ligands in this entry.

#### 5.7 Other polymers (i)

There are no such residues in this entry.

For Manuscript Review

#### 5.8 Polymer linkage issues (i)

There are no chain breaks in this entry.

For Manuscript Review

### 6 Map visualisation (i)

This section contains visualisations of the EMDB entry EMD-27812. These allow visual inspection of the internal detail of the map and identification of artifacts.

Images derived from a raw map, generated by summing the deposited half-maps, are presented below the corresponding image components of the primary map to allow further visual inspection and comparison with those of the primary map.

#### 6.1 Orthogonal projections (i)

##### 6.1.1 Primary map

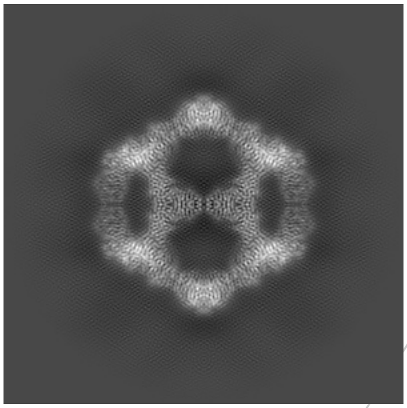

X

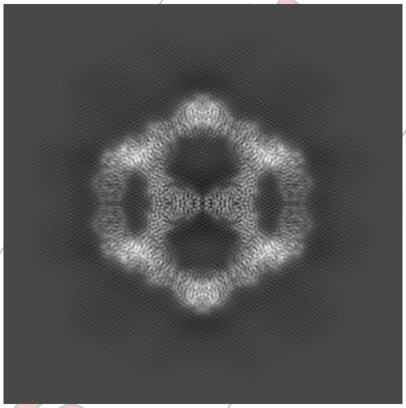

Y

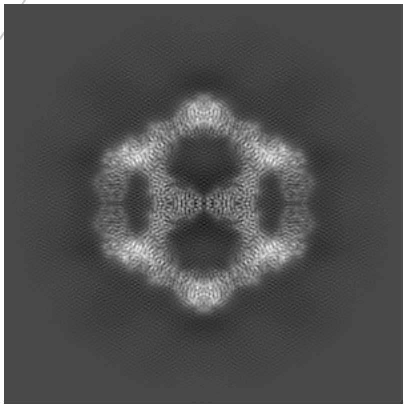

Z

##### 6.1.2 Raw map

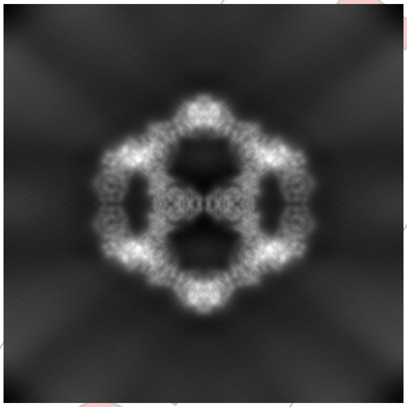

X

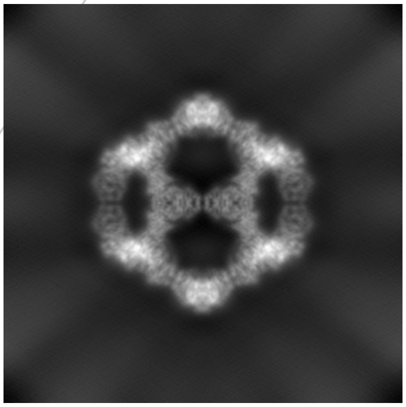

Y

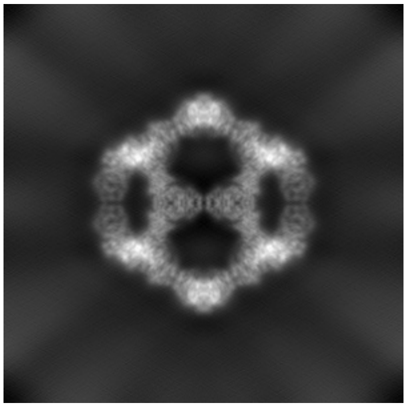

Z

The images above show the map projected in three orthogonal directions.

For Manuscript Review

#### 6.2 Central slices (i)

##### 6.2.1 Primary map

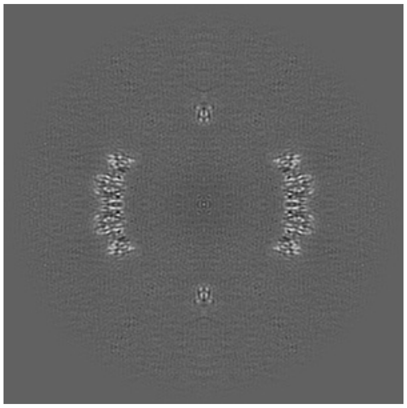
X Index: 210

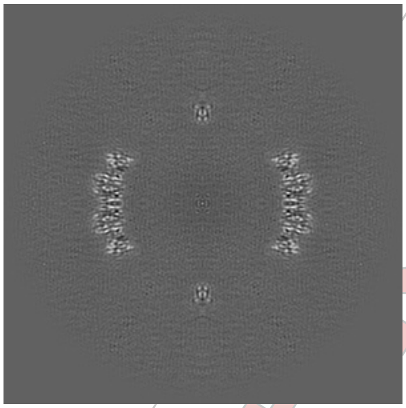
Y Index: 210

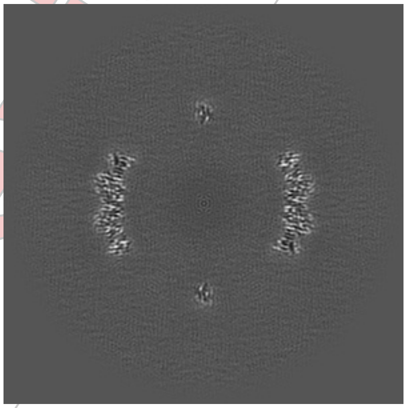
Z Index: 210

##### 6.2.2 Raw map

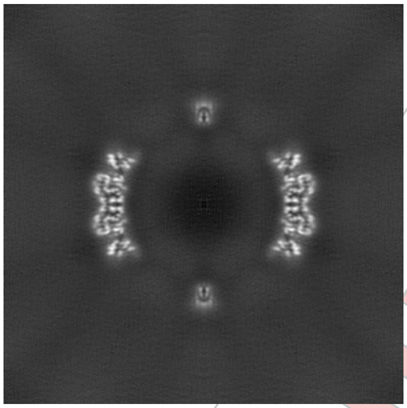
X Index: 210

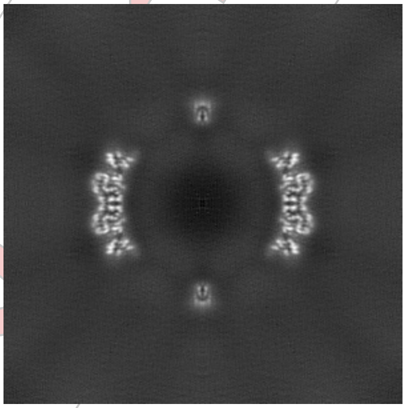
Y Index: 210

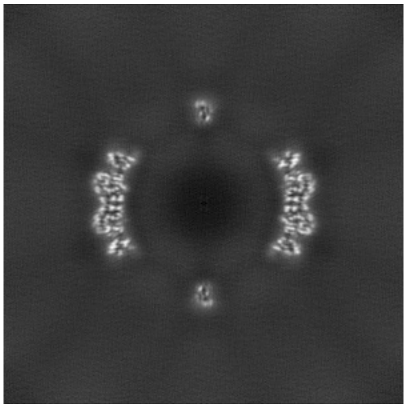
Z Index: 210

The images above show central slices of the map in three orthogonal directions.

For Manuscript Review

#### 6.3 Largest variance slices (i)

##### 6.3.1 Primary map

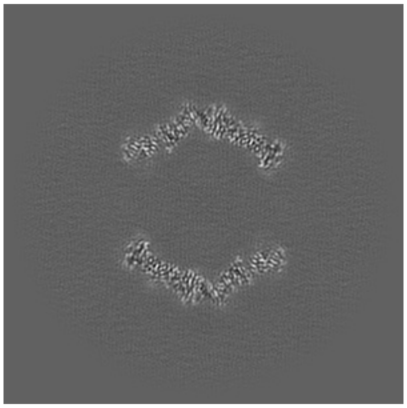
X Index: 162

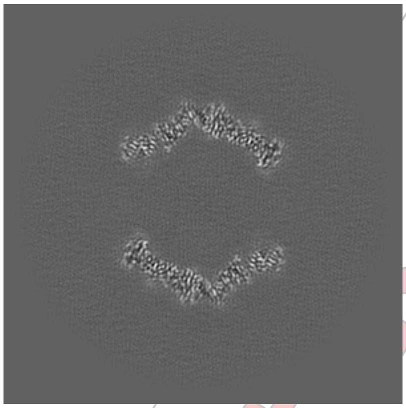
Y Index: 162

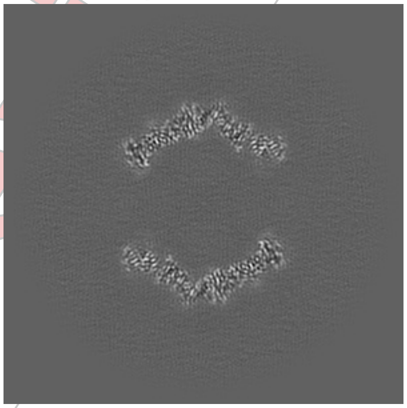
Z Index: 257

##### 6.3.2 Raw map

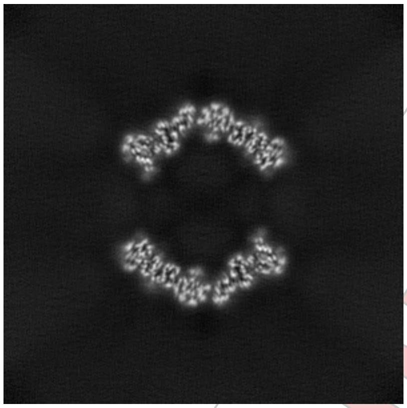
X Index: 158

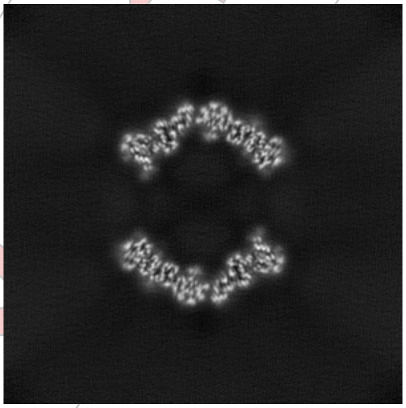
Y Index: 158

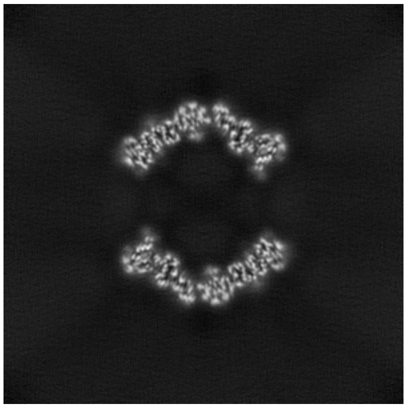
Z Index: 261

The images above show the largest variance slices of the map in three orthogonal directions.

For Manuscript Review

#### 6.4 Orthogonal surface views (i)

##### 6.4.1 Primary map

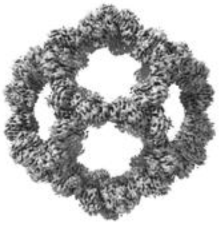

X

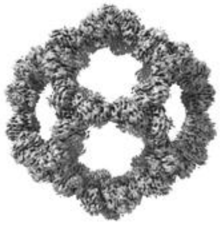

Y

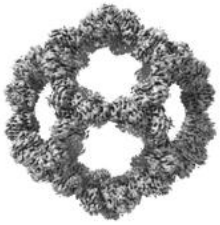

Z

The images above show the 3D surface view of the map at the recommended contour level 0.9. These images, in conjunction with the slice images, may facilitate assessment of whether an appropriate contour level has been provided.

##### 6.4.2 Raw map

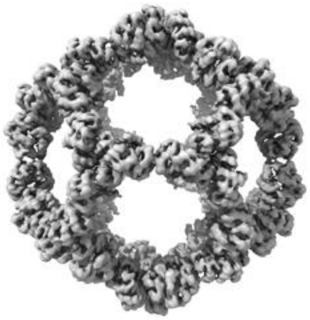

X

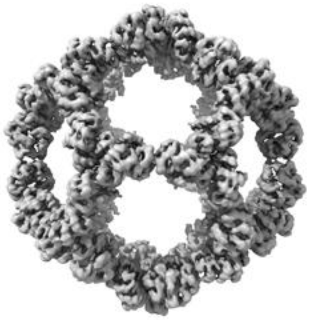

Y

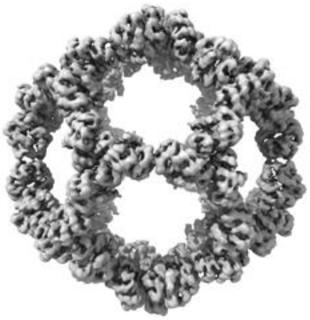

Z

These images show the 3D surface of the raw map. The raw map's contour level was selected so that its surface encloses the same volume as the primary map does at its recommended contour level.

For Manuscript Review

#### 6.5 Mask visualisation (i)

This section shows the 3D surface view of the primary map at 50% transparency overlaid with the specified mask at 0% transparency

A mask typically either:

- Encompasses the whole structure
- Separates out a domain, a functional unit, a monomer or an area of interest from a larger structure

##### 6.5.1 D_1000267547_em-mask-volume_P1.map.V3 (i)

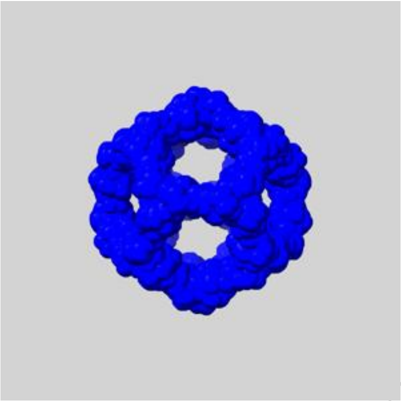

X

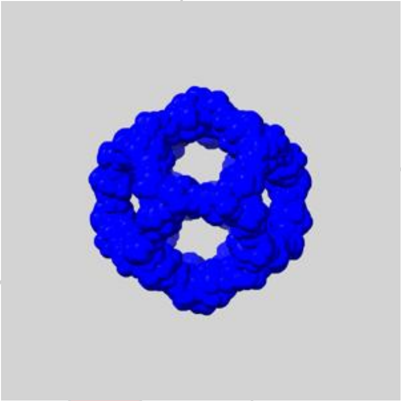

Y

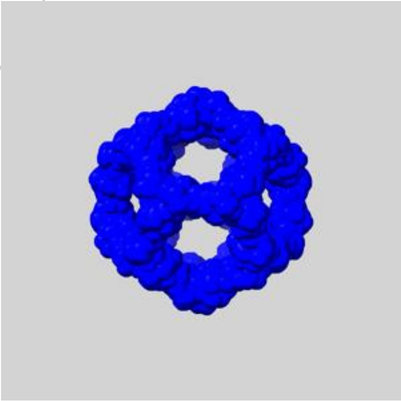

Z

For Manuscript Review

### 7 Map analysis (i)

This section contains the results of statistical analysis of the map.

#### 7.1 Map-value distribution (i)

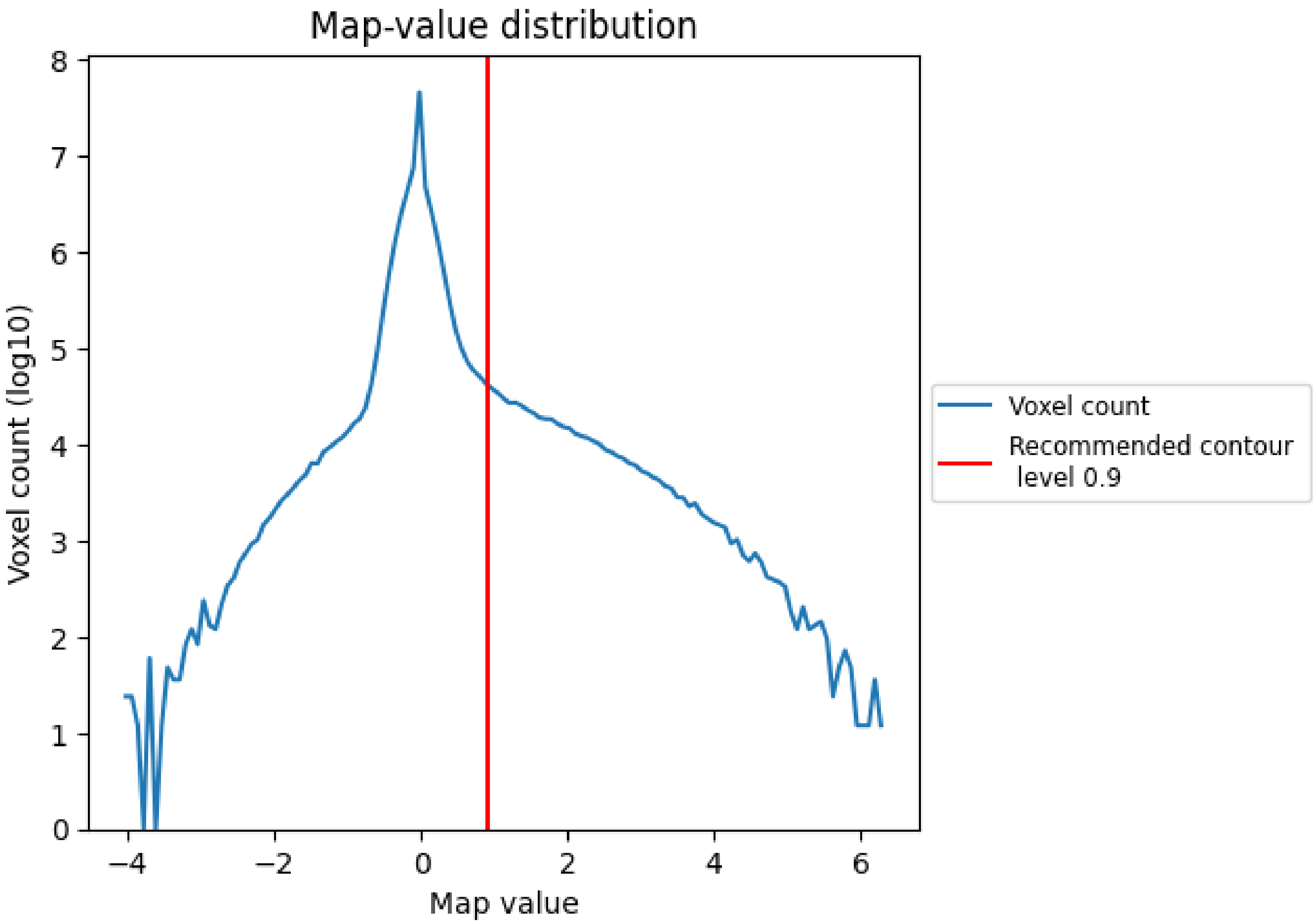

The map-value distribution is plotted in 128 intervals along the x-axis. The y-axis is logarithmic. A spike in this graph at zero usually indicates that the volume has been masked.

For Manuscript Review

#### 7.2 Volume estimate (i)

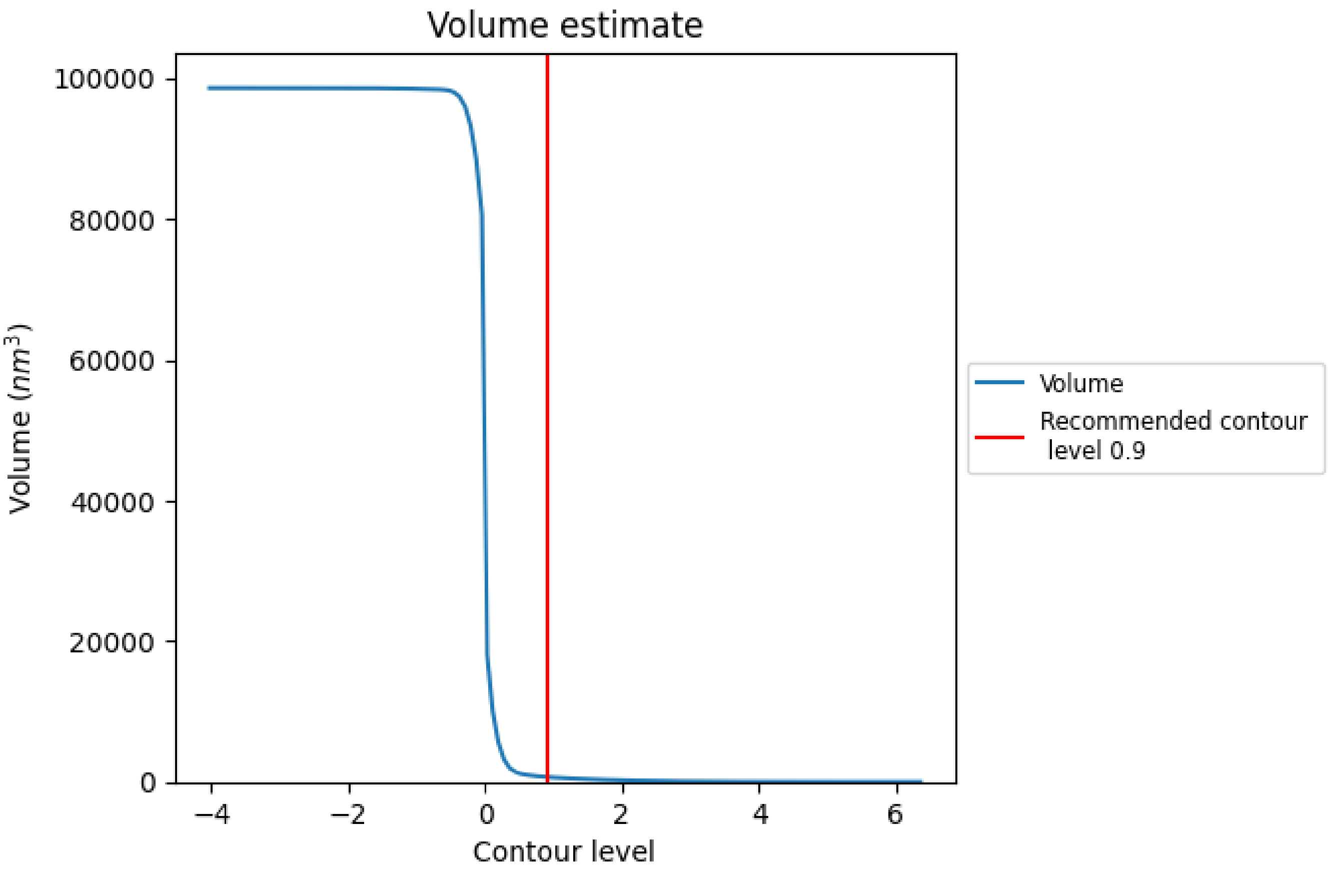

The volume at the recommended contour level is 694 $nm^3$; this corresponds to an approximate mass of 627 kDa.

The volume estimate graph shows how the enclosed volume varies with the contour level. The recommended contour level is shown as a vertical line and the intersection between the line and the curve gives the volume of the enclosed surface at the given level.

For Manuscript Review

#### 7.3 Rotationally averaged power spectrum (i)

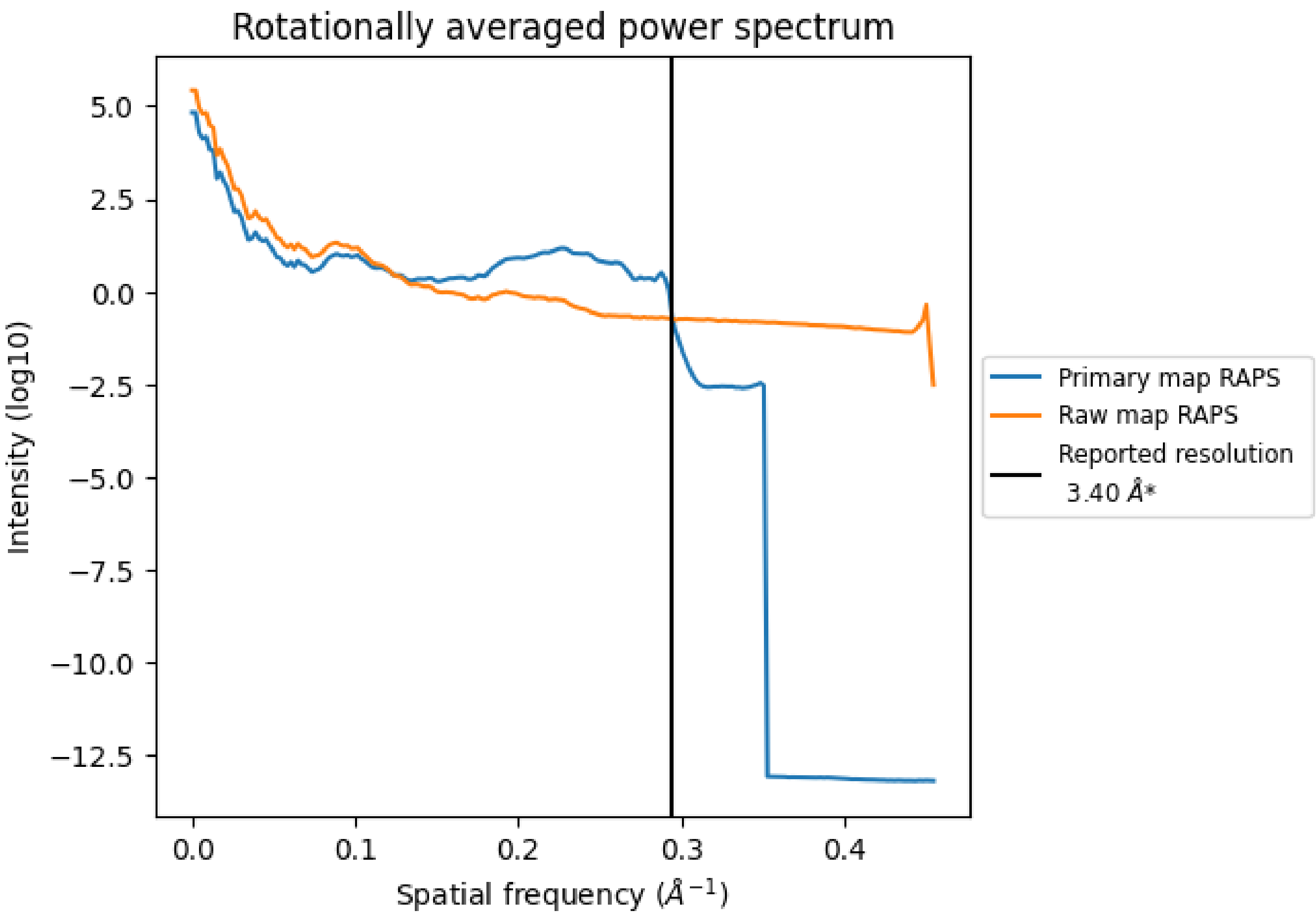

*Reported resolution corresponds to spatial frequency of 0.294 $Å^{-1}$

For Manuscript Review

### 8 Fourier-Shell correlation ⓘ

Fourier-Shell Correlation (FSC) is the most commonly used method to estimate the resolution of single-particle and subtomogram-averaged maps. The shape of the curve depends on the imposed symmetry, mask and whether or not the two 3D reconstructions used were processed from a common reference. The reported resolution is shown as a black line. A curve is displayed for the half-bit criterion in addition to lines showing the 0.143 gold standard cut-off and 0.5 cut-off.

#### 8.1 FSC ⓘ

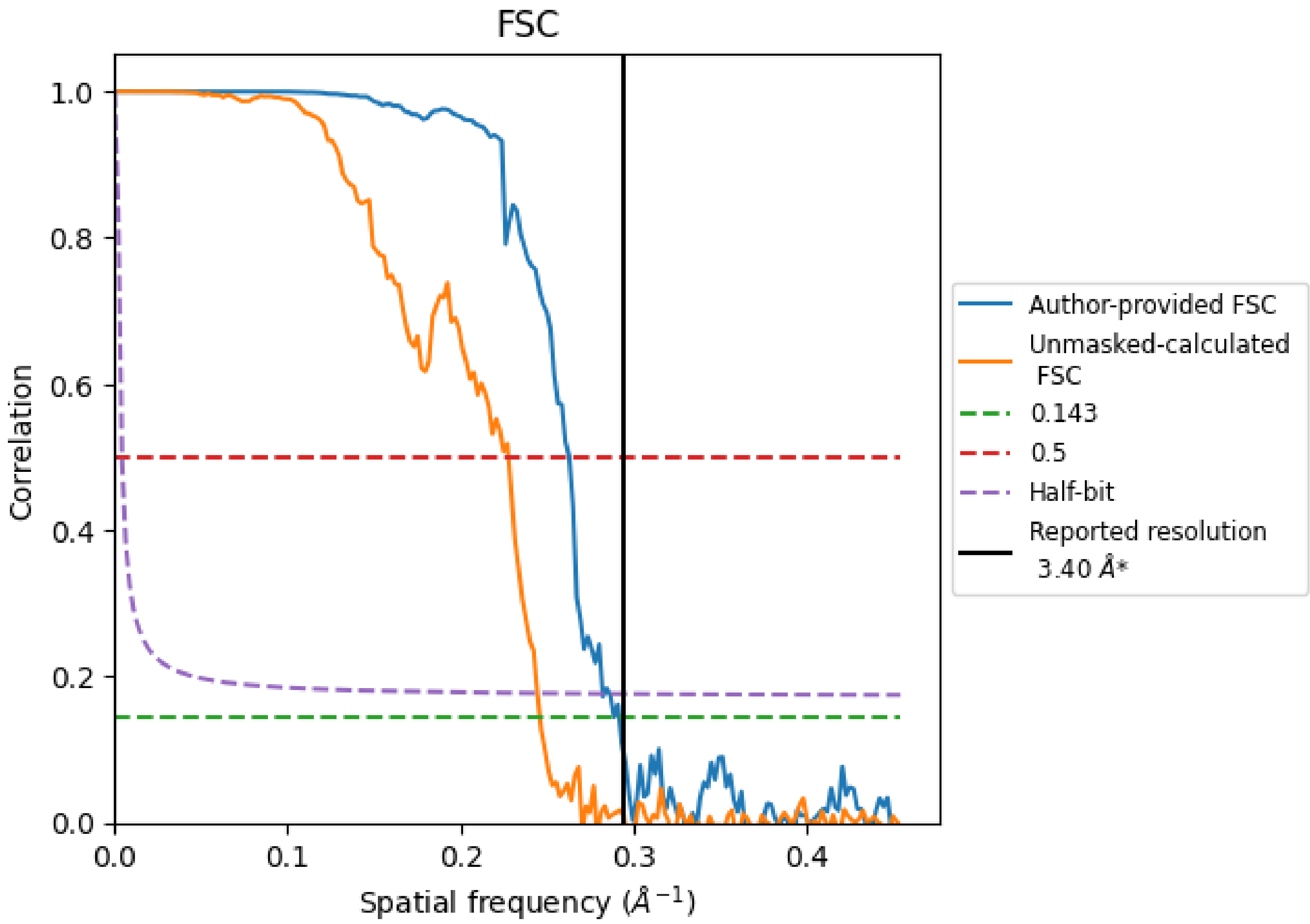

*Reported resolution corresponds to spatial frequency of 0.294 Å$^{-1}$

For Manuscript Review

#### 8.2 Resolution estimates (i)

| Resolution estimate (Å) | Estimation criterion (FSC cut-off) | | |
|---|---|---|---|
| | 0.143 | 0.5 | Half-bit |
| Reported by author | 3.40 | - | - |
| Author-provided FSC curve | 3.43 | 3.80 | 3.54 |
| Unmasked-calculated* | 4.07 | 4.39 | 4.09 |

*Resolution estimate based on FSC curve calculated by comparison of deposited half-maps. The value from deposited half-maps intersecting FSC 0.143 CUT-OFF 4.07 differs from the reported value 3.4 by more than 10 %

For Manuscript Review

### 9 Map-model fit (i)

This section contains information regarding the fit between EMDB map EMD-27812 and PDB model 8E01. Per-residue inclusion information can be found in section 3 on page 4.

##### 9.0.1 Map-model overlay (i)

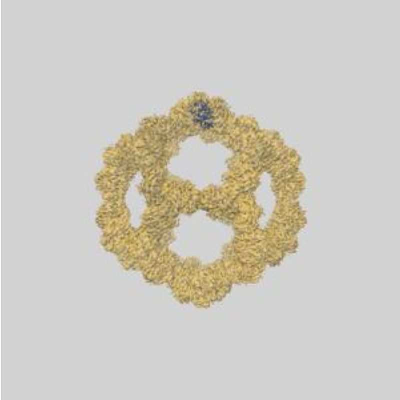

X

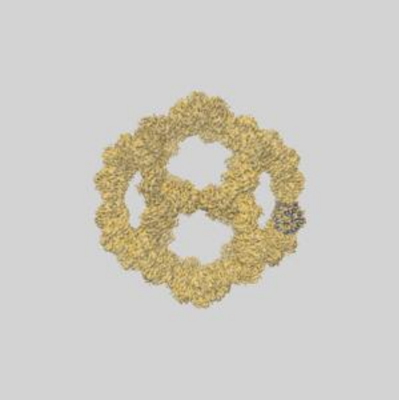

Y

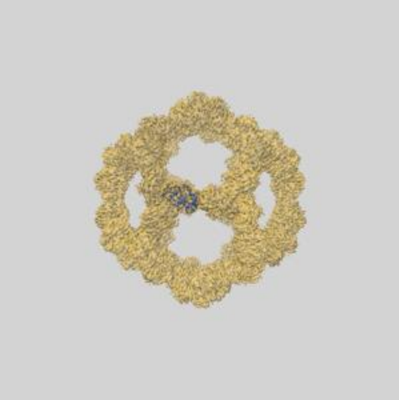

Z

##### 9.0.2 Map-model assembly overlay (i)

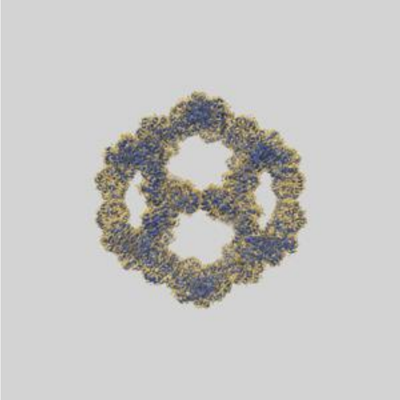

X

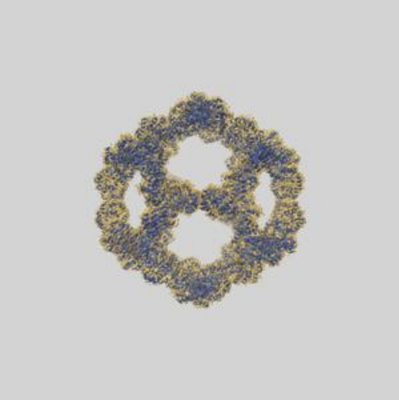

Y

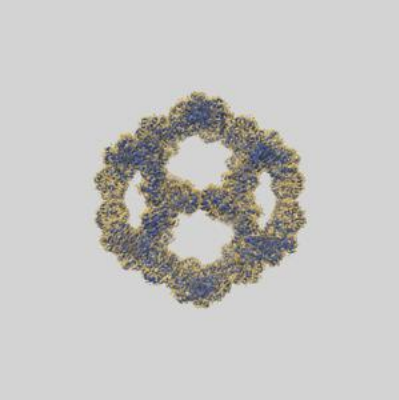

Z

The images above show the 3D surface view of the map at the recommended contour level 0.9 at 50% transparency in yellow overlaid with a ribbon representation of the model coloured in blue. These images allow for the visual assessment of the quality of fit between the atomic model and the map.

For Manuscript Review

#### 9.1 Atom inclusion (i)

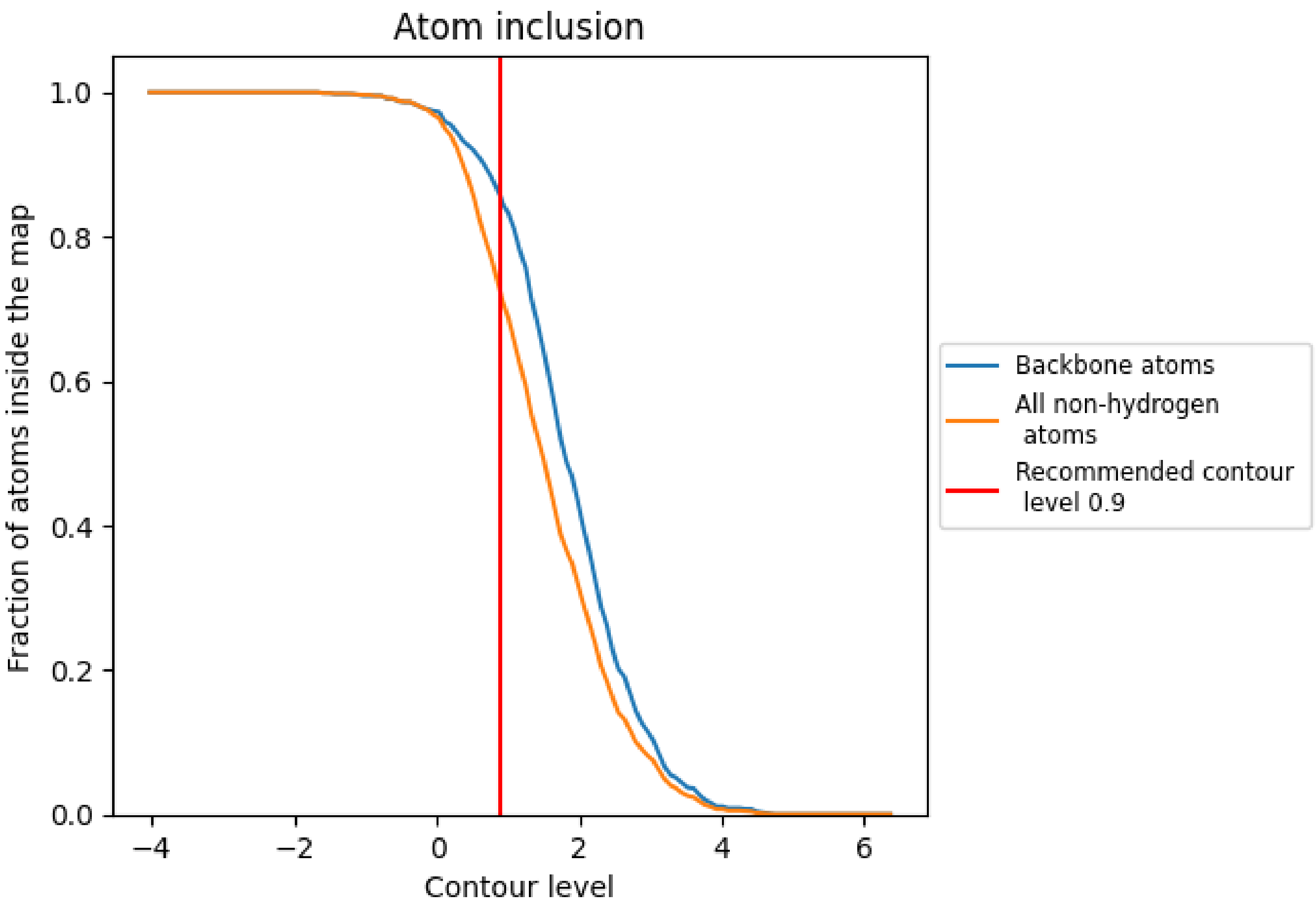

At the recommended contour level, 85% of all backbone atoms, 72% of all non-hydrogen atoms, are inside the map.

For Manuscript Review
